# Loss of the Coronary Artery Disease Risk Gene *Leiomodin1* in Vascular Smooth Muscle Cells Triggers Rapid Onset Coronary Atherosclerosis

**DOI:** 10.64898/2026.02.15.705944

**Authors:** Amr R. Salem, Jaser Doja, Chunyu Ge, Alshimaa Wally, Orazio J. Slivano, Susan H. Griffin, Brendan Marshall, Elizabeth Perry, Erin H. Seeley, Kunzhe Dong, Bhupesh Singla, Malgorzata Boczkowska, Gabor Csanyi, Roberto I. Vazquez-Padron, Vivek Nanda, Ajay Kumar, William B. Bryant, Roberto Dominguez, Xiaochun Long, Joseph M. Miano

## Abstract

**Background:** Atherosclerosis is the primary underlying cause of coronary artery disease (CAD). *Leiomodin1* is a vascular smooth muscle cell (VSMC)-restricted CAD risk gene whose role in coronary artery pathophysiology is unknown. Global loss of *Leiomodin1* causes lethal neonatal visceral myopathy, requiring unique approaches for study in VSMCs.

**Methods:** Several distinct *Leiomodin1* mutant mouse models were generated by clustered regularly interspaced short palindromic repeats (CRISPR). Control (*Lmod1^WT^*) and VSMC-restricted *Lmod1* knockout (*Lmod1^SMKO^*) mice were subjected to various atherogenic regimens. Atherosclerosis and LMOD1 expression in mouse and human coronary arteries were assessed by histopathology and confocal immunofluorescence microscopy. Coronary arteries from *Lmod1^WT^* and *Lmod1^SMKO^* mice were analyzed with assorted stains and antibodies, immunogold lineage tracing, spatial metabolomics/transcriptomics, and single-cell RNA sequencing (scRNA-seq). Mouse aortic SMCs from *Lmod1^WT^* and *Lmod1^SMKO^* mice were subjected to lipid loading with lentiviruses expressing wild-type *Lmod1*, a nucleation deficient *Leiomodin1* (*Lmod1^ND^*), or a short hairpin RNA (shRNA) targeting *Thrombospondin* (*Thbs1*).

**Results:** Under atherogenic conditions, *Lmod1^SMKO^* mice displayed unremarkable vessels in several organs but developed diffuse and occlusive coronary atherosclerosis. No such disease was observed in *Lmod1^WT^* mice. Time-course studies documented lipid insudation and VSMC foam cell formation in the coronary arteries of *Lmod1^SMKO^* mice as early as six days post-regimen. Immunogold lineage tracing demonstrated 46% of coronary plaque cells being of VSMC origin, with most showing evidence of lipid uptake. An intronic deletion of *Lmod1*, containing a conserved region where the single nucleotide variant associated with CAD exists, showed attenuated LMOD1 expression; heterozygous *Lmod1^SMKO^* mice, with a similar reduction in LMOD1, showed no CAD. Spatial metabolomics uncovered multiple lipid species within coronary atheromata of *Lmod1^SMKO^* mice, and spatial/scRNA-seq of similar coronary lesions disclosed altered lipid pathways with a consistent elevation in *Thbs1*. In vitro mechanistic studies revealed lipid accumulation in *Lmod1^SMKO^* VSMCs that was rescued by *Lmod1^WT^*, *Lmod1^ND^*, and *Thbs1* shRNA. VSMC-restricted expression of *Lmod1^ND^* in mice resulted in negligible coronary atherosclerosis.

**Conclusions:** Under proatherogenic conditions, *Lmod1^SMKO^* mice present with rapidly manifesting coronary atherosclerosis that appears to be independent of the actin nucleation function of LMOD1. Targeting *Thbs1* represents a viable strategy to mitigate VSMC foam cell formation.

**Clinical Perspective:** *What is new?:*

- Vascular smooth muscle cell (VSMC) loss of *Leiomodin1* (*Lmod1*) causes diffuse and occlusive coronary atherosclerosis in mice, with little or no such disease in other vascular beds.
- A novel immunogold lineage tracing assay shows VSMC migration to the intima as early as six days following an atherogenic regimen, and quantitative studies demonstrate that 46% of coronary plaque cells are of SMC origin.
- The coronary phenotype appears to be independent of LMOD1’s actin nucleation activity, but VSMC lipid uptake is thrombospondin-dependent.

*What are the clinical implications?:*

- LMOD1 is an annotated smooth muscle cell-restricted risk allele for human coronary artery disease (CAD), offering new insight into the role of smooth muscle cells in atherogenesis.
- The rapidly manifesting CAD phenotype in *Lmod1* knockout mice enables expedited testing of novel therapeutics to mitigate disease progression.
- New insight into LMOD1 pathobiology will help inform further SNV interrogation of the *LMOD1* locus for CAD risk in patients.

## Background

Ischemic heart disease continues to be the leading cause of age-standardized death in the world (https://www.healthdata.org/data-tools-practices/interactive-visuals/gbd-compare). ^1^ The underlying pathology in ischemic heart disease involves coronary artery atherosclerosis, a condition of lipid infiltration and modification, chronic inflammation, and lipid uptake by macrophages and vascular smooth muscle cells (VSMC) within the arterial intima, leading to atheromatous plaque formation, erosion or rupture, and consequent myocardial infarction. ^2,3^ While modifiable risk factors such as smoking, hypercholesterolemia, hypertension, and diabetes can be managed, non-modifiable factors, such as age and sex cannot. Further, genome wide association studies (GWAS) have identified heritable single nucleotide variants (SNVs) around 346 risk alleles for coronary artery disease (CAD). ^4^ Thus, there is established consensus for a genetic predisposition to CAD and, by extension, mortality due to ischemic heart disease. A small number of GWAS risk alleles have known mechanisms of action (eg, *LDLR*, *APOE*, *PCSK9*); however, the vast majority have poorly understood mechanisms for CAD risk. ^4–6^ The extensive historic and ongoing literature related to GWAS for CAD will require enormous efforts to experimentally elucidate individual and combined effects of SNVs on a patient’s risk of CAD.

Incontrovertible evidence demonstrates dynamic fate and state changes in VSMCs that contribute to atherogenesis. ^7–11^ Notably, three CAD risk genes (*FHL5*, *LMOD1*, and *MYH11*) are highly restricted to SMCs, but their roles in atherosclerosis have largely been limited to expression-based studies. ^12–14^ Two SMC-restricted CAD risk genes (*LMOD1*, *MYH11*) harbor coding mutations linked to visceral SMC diseases of the intestine and bladder. ^15,16^ Predictably, global inactivation of *Lmod1* or *Myh11* in mice results in early neonatal death due to visceral myopathy. ^15,17^ Conditional inactivation of these CAD risk genes using conventional SMC Cre mouse lines is likely to be problematic due to broad Cre activity across both vascular and visceral SMC lineages. ^18,19^ Recently, a more VSMC-restrictive *Itga8-CreER^T2^* mouse line was developed that circumvents lethal gastrointestinal phenotypes, expected with more traditional smooth muscle Cre drivers. ^20^ More recent studies have reported similar evasion of confounding visceral myopathies that would otherwise preclude or complicate assessment of VSMC phenotypes. ^21–23^

Here, we describe a rapidly manifesting coronary atherosclerosis mouse model, enabled by *Itga8-CreER^T2^*-mediated inactivation of *Lmod1* under various atherogenic conditions. No such phenotype was observed in control mice or in mice where the *Lmod1* gene is inactivated using the standard *Myh11-CreER^T2^* mouse. ^24^ In fact, the latter animals developed acute intestinal distention and died within one week of tamoxifen administration. Several assays – some never previously applied in mouse models of coronary atherosclerosis – underscore a critical role for normal LMOD1 expression and function in VSMCs to safeguard the coronary vasculature under hypercholesterolemic conditions.

## Materials and Methods

### Human coronary artery collection

The study cohort consisted of 12 deceased organ donors whose tissues were donated for research in collaboration with the Life Alliance Organ Recovery Agency (no informed consent required). Cross-sectional samples of the left coronary artery, approximately 2 cm in length, were obtained post-mortem following organ procurement procedures. Tissue was fixed in 10% neutral buffered formalin, embedded in paraffin, and sectioned at a thickness of 5 μm.

### Analysis of human coronary artery scRNA-seq and scATAC-seq data

Count matrix of scRNA-seq data generated from human coronary arteries from four cardiac transplant recipients was downloaded from GEO database (GSE131778) ^7^ and was analyzed as we previously described. ^25^ Briefly, data were processed using R package Seurat v4.1.1 ^26^ for normalization, dimensionality reduction, and clustering. Genes detected in fewer than five cells, and cells expressing fewer than 500 or more than 3,500 genes, or containing >7.5% mitochondrial genes, were excluded.

Cluster-specific genes were identified by using Seurat build-in “FindAllMarkers” function. Uniform manifold approximation and projection (UMAP) plots were generated to visualize cell clusters of *LMOD1* gene expression. The scATAC-seq count matrix from coronary artery segments of 41 human patients with varying stages of CAD was obtained from GEO database (GSE175621) ^27^ and processed using Signac v1.1.0. ^28^ Peaks from individual samples were merged into a unified peak set for downstream analysis. Cells with peak_region_fragments >3000, peak_region_fragments <20000, pct_reads_in_peaks >15, blacklist_ratio <0.05, nucleosome_signal <4 and TSS.enrichment >2 were kept for subsequent analysis. Peak annotation was performed with “GetGRangesFromEnsDb” (ensdb=EnsDb.Hsapiens.v86). Gene activity scores were calculated using “GeneActivity” function and visualized with “FeaturePlot”. Integration of scRNA-seq data of human atherosclerotic tissues ^7^ was conducted using “FindTransferAnchors” function and cell cluster was annotated by predicted labels and cluster-specific genes with a high gene activity score identified by “FindAllMarkers” function. Chromatin accessibility at the *LMOD1* gene locus was visualized using the “CoveragePlot” function.

### Mouse models

A total of 293 male and 220 female experimental mice were used in this study over a three-year period following ARRIVE 2.0 guidelines. ^29^ All parental mouse lines were of strain C57BL6/J and were maintained in a heterozygous state via repeated back-crossing to wild type C57BL6/J mice (RRID:IMSR_JAX:000664). Wild type breeders were “refreshed” every five generations to mitigate genetic drift. Experimental mice were inter-crossed no more than five generations to further minimize genetic drift. All mice were maintained in microisolator cages containing bedding enrichment within a temperature- and humidity-controlled environment under a 12-hour light-dark cycle. Clean water via lixit and a normal diet comprising 24% kcal protein, 58% kcal carbohydrate, and 18% kcal fat (Inotiv TD.2918, Lafayette, IN) were provided *ad libitum*. Mice carrying one copy of *Itga8-CreER^T2^* ^20^ (RRID:MMRRC_069930-UNC MMRRC) or *Myh11-CreER^T2^* ^24^ (RRID:IMSR_JAX:019079) were bred to a newly floxed *Lmod1* mouse, with Lox*P* sequences upstream of the core *Lmod1* promoter containing two SRF-binding CArG elements ^30^ and downstream of the splice donor site following exon 1 (full strategy available upon request) to generate homozygous floxed *Lmod1*/heterozygous *Itga8-CreER^T2^* (*Lmod1^fl/fl^/Itga8Cre*) or homozygous floxed *Lmod1*/heterozygous *Myh11-CreER^T2^* (*Lmod1^fl/fl^/Myh11Cre*) mice. Three cohorts of *Lmod1^fl/fl^/Itga8Cre* mice carried the *mTmG* reporter (RRID:IMSR_JAX:007676) for lineage tracing studies. To control for Cre toxicity ^31^, *Itga8-CreER^T2^* or *Myh11-CreER^T2^* siblings from the same litter of mice used to create homozygous null mice carrying a Cre allele, were simultaneously crossed to C57BL6/J breeders to generate wild type *Lmod1* mice carrying one of the Cre alleles (abbreviated *Lmod1^KO-Itga8^* or *Lmod1^KO-Myh11Cre^*). Group randomized *Lmod1^KO-Itga8^* or *Lmod1^KO-Myh11Cre^* mice and their respective controls of age 7-8 weeks were administered tamoxifen (Sigma #T5648; St. Louis, MO) at a dose of 50 mg/kg (ip) for five consecutive days followed by a 12-day washout period. In one experiment, we used *Lmod1^KO-Itga8^* mice treated with oil instead of tamoxifen. Another cohort of mice was studied as *Lmod1^fl/+^*/*Itga8Cre* heterozygous knockouts to test for haploinsufficiency. Adeno-associated virus serotype 8 (AAV8) carrying a gain-of-function mutation (D377Y) in mouse PCSK9 ^32^ (Penn Vector, University of Pennsylvania) was delivered to male and female mice by retro-orbital puncture. Because female mice exhibit less hypercholesterolemia than male mice following a lower dose of AAV8-PCSK9, ^33^ we injected all mice with 10e10 viral particles per mouse. Immediately thereafter, mice were fed a Clinton/Cybulsky high fat diet (HFD) consisting of 40% kcal carbohydrate (corn starch and granulated sucrose), 20% kcal protein, and 40% kcal fat (mainly cocoa butter), supplemented with 1.25% cholesterol (Research Diets, D12108C) from six days to 29 weeks. In some experiments, the initiation of the PCSK9/HFD was followed, two weeks later, by implantation of a mini-osmotic pump containing either saline or angiotensin II delivered at a rate of 1,000ng/kg/min for five days or up to four weeks at which time the experiment was terminated. Prior to pump implantation, mice were anesthetized with isoflurane inhalation followed by slow-release Buprenorphine to manage pain.

In a separate study carried out in Dr. Vivek Nanda’s lab, *Lmod1^KO-Itga8^* and *Apoe^-/-^* (strain #002052; The Jackson Laboratory, Bar Harbor, ME) double knockout mice were administered the same dose of tamoxifen (or equal volume of sunflower oil) and washout as above and then transitioned to a Western diet (WD) comprising 42.7% kcal carbohydrate, 15.2% kcal protein, and 42% kcal fat (anhydrous milkfat), supplemented with 0.2% cholesterol (Inotiv TD.88137) for 11 weeks. All cohorts of mice (**Table S1**) were euthanized at the terminal end point (or if ill-health was evident) by isoflurane anesthesia/cervical dislocation followed by PBS perfusion and rapid isolation of tissues for biochemical, histological, metabolic, immunological, transcriptomic, and ultrastructural analyses as detailed below.

The inducible *Lmod1* nuclease deficient knockin mouse (designated *Lmod1^iND^*), with five amino acid substitutions within the leucine-rich repeat domain of LMOD1, ^34^ was generated by Biocytogen Boston Corp (Waltham, MA). This mouse was crossed into the *Itga8-CreER^T2^* strain to generate homozygous *Lmod1^iND^*/heterozygous *Itga8CreER^T2^* mice (designated inducible nuclease deficient *Lmod1*, or *Lmod1^iND^*), which were treated with Tmx and the PCSK9/HFD regimen as above for 10 weeks. Heart, aorta, and blood were collected for subsequent Oil-Red-O (ORO) staining, qRT-PCR and Western blotting, and cholesterol measures, respectively. The *Lmod1* intron one 11,350 bp deletion mouse (excising the orthologous human sequences containing the lead CAD SNV, rs34091558 and two conserved CArG boxes) was generated by two-component CRISPR ^35^ and bred to homozygosity for Western blot and confocal immunofluorescence microscopy studies (CIFM).

All mice were genotyped with rigorous PCR protocols. Briefly, ear punch biopsies – used for individual mouse identification and PCR genotyping – were incubated at 55°C in digestion buffer (50 mM KCl, 10 mM Tris-HCl [pH9], 0.1% Triton X-100, and 1 μl of Proteinase K [NEB #P8107S]) for 2 hours followed by heat inactivation at 95°C for 10 minutes. A 1 μl sample of the digested biopsy was PCR amplified using GoTaq^®^ Green Master Mix (Promega #712) and specific primer sequences (**Table S2**). Optimized genotyping strategies and some strains of mice are available upon request. PCR products were resolved in 2.5% agarose gels and, in some cases, purified for Sanger sequencing (GENEWIZ^®^, Research Triangle Park, NC) to validate sequence integrity.

Institutional Animal Care and Use Committees reviewed and approved all mouse procedures conducted at Augusta University (#20191000) and the University of Alabama, Birmingham (#21997).

### Blood pressure

Blood pressure was measured using the CODA® High Throughput Tail-Cuff System (Kent Scientific, Torrington, CT), following the manufacturer’s guidelines. The system contains six activated channels and volume pressure recording sensor technology to record blood pressure in real time from a cuff placed around the tail of mice. Patency of the occlusion and volume pressure recording cuffs were checked routinely before the start of the experiments. Blood pressure measurements were conducted in a designated quiet area (22 ± 2 °C). Prior to data collection, mice were acclimated to the testing environment and trained in restraining tubes with 20 cycles of cuff inflation/deflation for three consecutive days to reduce stress and ensure reliable measurements. Blood pressure measurements were then recorded over two subsequent days. To ensure consistent measurement conditions, the tail temperature of each mouse was maintained between 32 °C and 37 °C. All trials were performed at the same time of day to minimize effects of diurnal arterial pressure variation. A maximum occlusion pressure of 250 mm Hg was applied, followed by a 20-second deflation period. Tail volume changes were surveilled during each cycle as blood flow returned to the tail post-cuff deflation, with a minimum volume change threshold set at 12 μL. Any cycles where the recorded blood flow was below 12 μL were excluded to ensure data accuracy. Each session consisted of 20 measurement cycles, with a minimum of 10 valid readings required per session. The final blood pressure values were reported as the mean ± standard error of the mean (SEM) derived from the measurements collected during the last two days of assessment.

### Blood assays

For serum lipid analysis, approximately 100 μL of blood was collected from live mice using Microvette® CB 300 Capillary Blood Collection Tubes (Sarstedt, #1060089). The samples were kept on ice and then centrifuged at 5,000 rpm for 15 minutes at 4 °C and the serum was isolated. Total cholesterol, LDL, and HDL levels were quantified using Cholesterol E (Wako Chemicals, #999-02601) and the HDL and LDL Assay Kit (Cell Biolabs, #STA-391), following the manufacturer’s instructions.

For complete blood count (CBC), electrolyte measurements, and liver and kidney function assays, blood was collected at the terminal time point via orbital exsanguination. Approximately 700–1,000 μL of blood was collected into Vacuette K3EDTA tubes (Greiner Bio-One, #454021). Plasma was separated by centrifugation under the same conditions as above, transferred into Eppendorf tubes, and stored at −80 °C until analysis. Samples were analyzed at the University of Rochester Medical Center (URMC) Clinical Trials Laboratory.

### Histochemical staining

Tissues were immersion fixed in 4% paraformaldehyde for 24-28 hours at room temperature (RT), followed by processing in a Leica TP1020 tissue processor, embedded with paraffin, and sectioned at 5 μm with an automated microtome (HM355S, ThermoScientific). Sections were initially stained with hematoxylin and eosin (H&E) using an automated stainer (Leica Autostainer XL). Masson Trichrome staining was performed on paraffin sections of heart using StatLab (#KTMTR2LT) according to the manufacturer’s instructions. Briefly, 5 μm cross-sections were deparaffinized in xylene twice for five minutes each, followed by immersion in absolute alcohol three times for one minute each. The slides were then immersed in running water. After deparaffinization, slides were fixed in Bouin’s solution for 2 hours at 60 °C, followed by staining with Wright’s hematoxylin for 5 minutes at 60 °C, and then washed under running water for 2 minutes. Next, slides were immersed in Biebrich Scarlet-Acid Fuchsin for 16 minutes at 60 °C, followed by rinsing in running water for 2 minutes. The slides were then immersed in phosphomolybdic/phosphotungstic acid for 15 minutes at 60 °C, and then stained with aniline blue for 5 minutes at 60 °C. Finally, stained slides were immersed in 1% acetic acid for 5 minutes, followed by dehydration in absolute alcohol three times for 1 minute each. The slides were then cleared in xylene twice for 2 minutes each and mounted with Cytoseal 60 (#8310-4). Images were captured using an Olympus BX43 microscope with cellSens Standard 4.4 software (Olympus, Center Valley, PA).

For Oil-Red-O (ORO) staining, tissues (aorta, brain, heart, kidney, liver, lung, mesentery, and spleen) were rapidly removed from the mouse and fixed in 4% paraformaldehyde for 24-48 hours at 4°C. Samples were removed from fix and placed in 30% sucrose for 24-48 hours at 4°C to cryoprotect each tissue. Next, tissues were embedded in optical cutting temperature compound at −25°C and allowed to set for two minutes. Sections (7 μm) were made with a Leica CM1950 Cryostat and placed on RT slides. ORO staining was performed using StatLab reagent (#KTOROLT). The sections were fixed in Bouin’s solution for five minutes, followed by rinsing in running water for three minutes. Next, the slides were immersed in propylene glycol for two minutes at RT and then incubated in preheated ORO solution at 60 °C for six minutes. Differentiation was performed by immersing the slides in 85% propylene glycol for two minutes, followed by rinsing with running deionized (DI) water for two minutes. The slides were then counterstained with modified hematoxylin for two minutes at RT, rinsed in running tap water for one minute, and mounted using ProLong Antifade (Thermo Scientific, #P36981). Images were acquired as above.

To visualize collagen deposition, paraffin sections were deparaffinized with xylene using the standard protocol described above, followed by hydration in running water for 3 minutes. The slides were then immersed in Picrosirius Red (Lab Chemical #SRS500) for one hour at RT. After staining, the slides were washed in 1% acetic acid twice for one minute each, followed by dehydration in absolute alcohol through three changes, each for one minute. The slides were then cleared with xylene twice for one minute each and mounted with Cytoseal 60 (#8310-4). Images were captured using an Olympus BX43 microscope and areas of Picrosirius Red staining calculated with ImageJ. To stain for calcification, paraffin sections were deparaffinized as described above and then immersed in Alizarin Red stain (StatLab, #STARE100) for five minutes at RT. Sections were then dehydrated twice for 30 seconds each in acetone, cleared in xylene (3x for 1 minute each) and mounted with Cytoseal 60 (Thermo Scientific, #8310-4). Stained sections were visualized with an Olympus BX43 microscope and images taken as above.

### Western blotting

Aortas were isolated, cleaned, and lysed in RIPA buffer (Sigma-Aldrich #R0278) supplemented with protease inhibitor cocktail (Roche #04693159001) and PMSF (Sigma-Aldrich #10837091001).

Protein concentrations were determined using the Protein Assay Kit II (Bio-Rad #5000112), and an average of 5 μg of protein was mixed with Laemmli Sample Buffer (Bio-Rad #1610747), boiled, cooled on ice for 10 minutes, and separated on a 4% to 15% Criterion TGX stain-free gel (Bio-Rad #5678083) using Tris-Glycine-SDS Buffer (10X, Bio Basic #A0030). Proteins were transferred onto PVDF membranes (Bio-Rad #1620177) using the Trans-Blot Turbo Transfer System (Bio-Rad #1704150) with the Trans-Blot Turbo Transfer Pack (Bio-Rad #1704275). Membranes were blocked and incubated overnight at 4 °C with primary antibodies against LMOD1 (Proteintech #15117-1-AP), SRF (Cell Signaling Technology #5147S), ACTA2 (Sigma-Aldrich #A2547-0.2ML), ACTB (Affinity #T0022), GAPDH (Millipore #MAB374), ITGA8 (Santa Cruz #SC-365798), TAGLN (Abcam #ab14106), and Thrombospondin-1 (THBS1) (Abcam #ab85762) (**Table S3**). After incubation, membranes were washed three times with PBST (Boston BioProducts #IBB-171) for 10 minutes each, followed by a 2-hour incubation at RT with secondary antibodies at a 1:5000 dilution, including Goat anti-Rabbit (Novus Biologicals #NB7183), Goat anti-Mouse (Novus Biologicals #NBP1-75130), and Goat anti-Rat (Novus Biologicals #AF005). After additional washes, membranes were developed using a chemiluminescent detection kit (Advansta #K-12043-D20) and imaged using a BioRad Chemi-Doc^™^ MP Imaging System. Western blot bands were quantified in ImageJ. Relative band density was normalized to the loading control (GAPDH or ACTB).

### Vascular histomorphometry and quantitative measures of atheromatous disease

Aortic segments from ascending aorta, between the 3^rd^ and 4^th^ intercostal space, and just below the left renal artery were immersion fixed in 4% paraformaldehyde, processed, embedded in wax, and sectioned (horizontally) at 5 μm for H&E staining to calculate aortic circumference and luminal area.

Hearts (aortic root to apex) were similarly fixed and processed for horizontal sectioning. Varying levels of heart were sectioned (5 μm) from apex to root at 300 μm to 1 mm intervals, stained with H&E, and imaged for coronary artery circumference, luminal area, and plaque area. All quantitative measures were done using ImageJ. We used the Stary classification of atherosclerosis lesion types ^36,37^ in 340 and 164 vessels from 68 *Lmod1^fl/fl^/Itga8-CreER^T2^* knockouts and 49 *Lmod1^+/+^/Itga8-Cre* controls, respectively.

### Confocal immunofluorescence microscopy (CIFM)

Paraffin-embedded heart sections were deparaffinized as described in the H&E staining protocol, followed by heat-mediated antigen retrieval using either Dako Target Retrieval Solution, pH 9 (10x) (Agilent Technologies #S236784-2) or Dako Target Retrieval Solution, Citrate pH 6 (10x) (Agilent Technologies #S236984-2) in a pressure cooker for seven minutes, depending on the antibody manufacturer’s instructions. After antigen retrieval, slides were left to cool at RT and then washed three times with 1x PBS. Next, sections were blocked with Protein Block, Serum-Free (Liquid form, 110 mL; Agilent Technologies #X090930-2) for 30 minutes at RT. Primary antibodies (**Table S3**) were diluted in Antibody Diluent (Abcam #ab64211) and applied to the slides. After incubation, slides were washed three times in 1x PBS, followed by incubation with secondary antibody for one hour at RT. Afterward, slides were washed three times in 1x PBS and mounted with ProLong Antifade (Thermo Scientific #P36981). Images were captured using a Zeiss LSM900 confocal microscope and analyzed with Zen blue software 3.5 (Zeiss Microscopy, White Plains, NY).

For frozen sections, tissues were fixed in ice cold acetone for 10 minutes and left to dry for another 10 minutes at RT. Slides were then washed three times in 1x PBS (one minute each) and permeabilized with 1x SDS for five minutes at RT. After permeabilization, slides were washed three times in 1x PBS (five minutes each), followed by blocking, primary and secondary antibody incubation, mounting, and visualization, as described for paraffin sections.

### Quantitative IFM and ORO

Two methods were employed to quantify immunofluorescence microscopy (IFM) staining area and colocalization. First, IFM staining for markers such as oxLDL was evaluated based on the total area above a set threshold. Analysis was performed using Zeiss Zen blue software 3.5 software. In the image-analysis panel, thresholds were set to 18–255 for AF568 (oxLDL) and 41–255 for DAPI. Individual nuclei were then counted by Zeiss Zen blue 3.5 software using these parameters. Data tables containing nuclei counts and total oxLDL staining area were exported, and total oxLDL per cell was calculated in GraphPad Prism. Second, colocalization assessments for CD68, SPP1, LGALS3, and CD45 with either GFP or ACTA2 were conducted using the colocalization feature in ZEN Blue. These markers were evaluated from the internal elastic lamina (IEL) inward, including the neointima but excluding the medial layer. THBS1 and MYH11 were quantified in the same fashion; however, for these two proteins, the medial layer was also included by quantifying from the external elastic lamina inward. The thresholds for each marker were set as follows: THBS1 (13:13), CD68 (30:30), SPP1 (30:30), LGALS3 (30:30), CD45 (30:30), and MYH11 (20:20). Tables containing the total colocalized fluorescence area with either GFP or ACTA2 were exported and used in GraphPad Prism to generate bar graphs.

### RNA isolation and quantitative RT-PCR

Total RNA was extracted from aortas using the RNeasy Plus Mini Kit (Qiagen #74136) following the manufacturer’s instructions. Reverse transcription was performed using the iScript Reverse Transcription Supermix (Bio-Rad #1708841BUN) in the presence of DNAse I. Quantitative RT-PCR was conducted in 10 μL reactions using iQ™ SYBR Green Supermix (Bio-Rad #1708886) on a Bio-Rad CFX Connect detection system, following the manufacturer’s specifications. Final data were calculated using the 2^-ΔΔCt^ method, with *Gapdh* or *Actb* as reference genes. Results are shown as the relative level of normalized expression to the control group. A list of all PCR primers is provided in **Table S2**.

### Dual reporter lineage tracing

To map the state and fate of the VSMC, *Lmod1^WT^* (carrying *Itga8-CreER^T2^*) and *Lmod1^SMKO^* were crossed with Rosa26 mTmG mice. Mice carrying all alleles were induced at eight weeks of age using the same tamoxifen regimen described above. Heart and aortas were harvested and fixed in 4% paraformaldehyde (PFA), embedded in OCT, and cryosectioned. Sections were briefly dipped in 95% ethanol (5 min), rinsed in running H₂O, mounted using Antifade NucBlue with DAPI (Thermo Fisher P36981), and imaged by confocal microscopy (Zeiss). Medial VSMCs and VSMC-derived cells of atheromata are membrane-GFP (green), whereas non-recombined (non-VSMC) cells remain membrane-tdTomato (red). Co-immunostaining of lineage-traced heart sections included antibodies to MYH11, CD68, SPP1, and LGALS3. Briefly, primary antibodies were incubated on slides overnight at 4 °C, washed 3x in 1xPBST, and incubated with the Alexa 647 secondary antibody for 30 min at RT. Sections were washed twice, treated with an autofluorescence quenching reagent (Vector Laboratories SP-8500-15) according to the manufacturer’s instructions, mounted, and imaged by confocal microscopy.

### Proliferation and cell death assays

To assess the in vivo proliferation of coronary VSMCs, mice were injected ip with 100 μg of EdU (Thermo Scientific, #C10337) either once 24 hours prior to termination or every other day for a total of 13 doses, starting on day eight after the initiation of PCSK9/high-fat diet. 12 hours following the final injection, mice were euthanized and perfused with PBS. Hearts and intestine (positive control) were isolated and fixed in 4% PFA, embedded in paraffin, and cut at 7 μm. Sections were deparaffinized, permeabilized with 0.5% TritonX-100 for five minutes, and then prepared for testing with the Click-iT^TM^ EdU Alexa Fluor^TM^ 488 imaging kit (Thermo Scientific, #C10337) according to the manufacturer’s instructions. For apoptosis, we followed a similar series of steps, including the inclusion of intestine as a positive control and followed the protocol in the Click-iT^TM^ Plus TUNEL assay kit for in situ detection of apoptotic cells (Thermo Fisher, #C10619). Images were captured using a Zeiss LSM900 confocal microscope and analyzed with Zen blue software 3.5 (Zeiss Microscopy, White Plains, NY).

### Mouse aortic SMC cultures

Mouse aortic smooth muscle cells (MASMCs) were isolated following a previously established protocol with some modifications. ^38^ Briefly, aortas were digested in Hank’s balanced salt solution containing 1mg/ml of Collagenase II (Worthington, LS004174) and 1 mg/ml Soybean Trypsin Inhibitor (Worthington, LS003570) for 20 minutes to remove the outer adventitial layer. Aortas were cut open and the endothelial layer was gently removed using a cotton swab. The cleaned aortas were minced into small pieces and incubated in 67 µL Elastase; (Worthington, LS002279) at a concentration of 24.8 mg/mL for 30 minutes at 37^0^C. Aortic pieces were then dissociated with a P1000 pipette. Dispersed aortic pieces were re-incubated in the elastase for an additional 30 minutes to prepare single cell suspensions. The digestion was terminated by adding complete media (DMEM-F12 supplemented with 10% mL fetal bovine serum (Thermo Fisher, A3160601), 1× Smooth Muscle Growth Supplement (SMGS) (Thermo Fisher, S00725) and antibiotics (Penicillin-Streptomycin). Cells were centrifuged at 1,000 rpm for five minutes to pellet the cells. The entire cell suspension was plated onto a single well of a 6-well plate. After approximately three days, the top 2/3 of the medium (∼1.6 mL) was carefully removed and an equal volume (1.6 mL) of fresh, pre-warmed media was gently added. Primary MASMC were passaged no more than five times.

### Actin nucleation deficient LMOD1

The cDNA encoding human *LMOD1* (UniProt: P29536-1) was purchased from Open Biosystems (Huntsville, AL). The fragment corresponding to amino acids 313–600 was cloned between the *NotI* and *HindIII* sites of a modified pMAL-c2x vector. ^34^ Point mutations corresponding to K449D, H453D, R460D, R475D, and R478D (equivalent to K444D, H448D, R455D, R470D, and R473D in mouse *Lmod1*) were introduced using the QuickChange mutagenesis kit (Agilent Technologies, Santa Clara, CA). Proteins were expressed in ArcticExpress (DE3) cells (Invitrogen), grown in Terrific Broth at 37 °C to an OD_600_ of 1.5–2.0, and induced with 0.5 mM isopropyl-β-D-thiogalactoside followed by incubation for 24 h at 10 °C. Cells were harvested by centrifugation, resuspended in 20 mM HEPES pH 7.5, 200 mM NaCl, 1 mM EDTA, and 100 μM phenylmethylsulfonyl fluoride, and lysed using a microfluidizer (Microfluidics).

Proteins were purified by amylose affinity chromatography according to the manufacturer’s protocol (New England Biolabs), followed by ion-exchange chromatography on a MonoS column (Cytiva) using a linear gradient from 50 to 500 mM NaCl in 20 mM MES pH 6.5.

### Actin polymerization assay

Actin polymerization was monitored as the increase in fluorescence resulting from the incorporation of pyrene-labelled actin into filaments, using a Cary Eclipse fluorescence spectrophotometer (Varian Medical Systems, Palo Alto, CA). Prior to data acquisition, 200 μl Mg–ATP–actin (2 μM; 6% pyrene-labelled) was mixed with 5 μl LMOD1 in 10 mM Tris (pH 8.0), 1 mM MgCl₂, 50 mM KCl, 1 mM EGTA, 0.2 mM ATP, 0.5 mM DTT, and 0.1 mM NaN₃. Final LMOD1 concentrations in the polymerization reactions are indicated below (**Figure 8B**). Data acquisition was initiated 20 seconds after mixing. All measurements were performed at 25 °C. Control reactions were performed by addition of 5 μl buffer.

### Lentivirus transduction

All Lentiviruses (shRNA and overexpression) were designed and prepared by a commercial vendor (Vector Builders, Chicago, IL; **Table S3**). MASMCs were seeded and the following day transduced at 50-60% confluency. Indicated lentivirus (5 vp/cell) was added in fresh culture media in the presence of 5 μg/ml of Polybrene. Seventy-two hours later, cells were selected in Puromycin (2.5 ug/ml) for an additional three days. Overexpression or knockdown was confirmed by protein expression (Immunoblot/Immunostaining).

### Lipid uptake studies

Oxidized LDL (Dil-oxLDL; L34357, ThermoFisher) uptake studies were conducted in MASMCs isolated from *Lmod1^WT^* (carrying *Itga8-CreER^T2^*) and *Lmod1^SMKO^* mice following Tmx administration.

Briefly, cells were plated in eight-well chamber slides at 60-70% confluency. The following day, cells were serum-starved in basal DMEM-F12 media supplemented with 0.3% bovine serum albumin (BSA) for four hours. Cells were then primed with 10 ng/ml TNFα (R&D Systems, #210-TA-005/CF) for 30 minutes before adding 10 μg/ml Dil-oxLDL in DMEM-F12 media with 0.3% BSA. Cells were incubated for an additional four hours at 37 °C. At the end of incubation, cells were washed three times with PBS and mounted with NucBlue. In some experiments, cells were subjected to immunostaining following oxLDL uptake. Briefly, at the last PBS wash step, cells were permeabilized with 0.5% Triton X-100 for 10 minutes, blocked with 3% BSA in PBST followed by primary antibody incubation at 4 °C overnight.

### Transmission electron microscopy

A total of 11 control (Tmx-treated heterozygous *Itga8-CreER^T2^*) and 14 conditional *Lmod1* knockout (heterozygous *Itga8-CreER^T2^*/*Lmod1^fl/fl^*) mice were perfused with heparinized PBS followed by fixative comprising 4% paraformaldehyde and 2% glutaraldehyde in 0.1 M sodium cacodylate buffer, pH 7.4. Hearts were dissected, immersion fixed in the same fixative overnight, post-fixed in 2% osmium tetroxide in sodium cacodylate for two hours and stained en bloc with 2% uranyl acetate and Reynolds lead citrate. Following dehydration with a graded ethanol series (25%, 50%, 70%, 80%, 95%, 100%), tissues were treated with propylene oxide (as an intermediary solvent) and then embedded in Embed 812/Araldite resin (Electron Microscopy Sciences, Ft. Washington, PA). Semi-thin sections (750 nm thick) were cut, mounted on glass slides, and stained with Toluidine Blue to map areas of interest. Thin sections (50 nm) were then cut with a diamond knife (Diatome/Electron Microscopy Sciences) on a Leica EM UC7 ultramicrotome (Leica Microsystems, Inc, Bannockburn, IL) and collected on Pioloform coated Synaptek copper slot grids (Electron Microscopy Sciences, Ft. Washington, PA). Grids were stained with 4% aqueous uranyl acetate and Reynolds lead citrate. Sections were observed in a JEM 1400Flash transmission electron microscope (JEOL USA Inc., Peabody, MA) at 120 kV and imaged with a OneView CCD Digital Camera (Gatan Inc., Pleasanton, CA). TEM data shown here were selected from a total of 109 (wild type) and 343 (knockout) images. A large powerpoint deck of annotated images is available upon request.

### Immunogold electron microscopy lineage tracing (IEMLT)

A total of five control (Tmx-treated double heterozygous *mTmG/Itga8-CreER^T2^*) and 16 *Lmod1* knockout (Tmx-treated *Lmod1^fl/fl^/* double heterozygous *Itga8-CreER^T2^/mTmG*) mice were perfused with heparinized PBS followed by fixative comprising 4% paraformaldehyde and 0.2% glutaraldehyde in 0.1 M sodium cacodylate sodium cacodylate buffer, pH 7.4. Following immersion fixation in the same 4% paraformaldehyde, 0.2% glutaraldehyde fix, hearts were processed through a graded series of ethanol (25%, 50%, 70%, 80%, 95%). Hearts were then embedded in LR White Resin (Electron Microscopy Sciences, Ft Washington, PA). Semi-thin sections, 750 nm in thickness, were cut and stained with Toluidine Blue to map areas of interest. Thin sections (50 nm) were cut with a diamond knife (Diatome/Electron Microscopy Sciences) on a Leica EM UC7 ultramicrotome (Leica Microsystems, Inc, Bannockburn, IL) and collected on Pioloform coated nickel slot grids. Grids, section-side down, were floated on drops of etching solution (5% Sodium Metaperiodate) in PBS for 5 minutes, then washed with PBS. Aldehydes were quenched 30 minutes with 1M ammonium chloride in PBS. Grids were then blocked in Aurion Blocking Solution (Electron Microscopy Sciences, Ft. Washington, PA), in PBS for 2-4 hours at RT. Grids were floated on drops of primary anti-GFP antibody (Novus Biologicals, Minneapolis, MN), diluted 1:100 in Aurion BSA-C incubation solution (Electron Microscopy Sciences) overnight at 4 °C and washed in PBS. Grids were then floated on drops of anti-primary species-specific Aurion Ultrasmall gold reagent (Electron Microscopy Sciences) diluted 1:200 in Aurion BSA-C for 2 hours at RT, then washed in PBS, followed by deionized water. Ultrasmall particles were silver enhanced for 19 minutes using HQ Silver™ (Nanoprobes, Yaphank, NY), then washed in ice cold deionized water to halt enhancement. Grids were then stained with 4% uranyl acetate and lead citrate to increase contrast.

Sections were observed in a JEOL 1400 Flash transmission electron microscope (JEOL USA Inc., Peabody, MA) at 120 kV and imaged with One View Digital Camera Controller (Gatan Inc., Pleasanton, CA). IEMLT images shown here were selected from a total of 24 wild type and >200 knockout images. A powerpoint deck of annotated images is available upon request.

To more accurately quantitate the percentage of GFP-derived cells in plaques, tiled low-magnification (800x) TEM images around the entire circumference of each of eight vessels (from eight knockout hearts) were generated and membrane-labeled GFP cells manually counted versus non-labeled cells. Only clearly-labeled GFP positive cells were counted (at least 50% of the membrane); GFP aggregates and fragments of GFP-labeled cells were excluded to rule out efferocytosis. These strict criteria, coupled with lower recombination efficiency of the *mTmG* reporter, ^39^ likely resulted in an underestimate of the percentage of coronary SMC-derived plaque cells. From these tiled images, we also calculated the percentage of GFP positive cells at the luminal border versus the core of each lesion.

### Bulk RNA-seq

Aortas from *Lmod^SMKO^* and *Lmod1^WT^* mice at baseline (15 days post-Tmx washout) and six weeks after PCSK9/HFD initiation were dissected, cleaned of perivascular tissue and endothelium, and snap-frozen at −80 °C. Total RNA was isolated using TRIzol (Thermo Fisher #15-596-018), chloroform (Sigma-Aldrich #C2432-1L), and isopropanol (Fisher Scientific #BP2618500). To eliminate DNA contamination, RNase-Free DNase (Promega #M6101) was used. A minimum of 2 μg of RNA was used for mRNA sequencing (Ribo-minus RNA-seq) at the University of Rochester Genomics Research Center.

Raw 150 bp paired-end sequences (Sanger/Illumina 1.9 encoding) were quality-controlled using FastQC v0.11.4 (http://www.bioinformatics.babraham.ac.uk/projects/fastqc/). Low-quality bases (quality scores <30) and adapter contamination (if present) were removed using Trimmomatic v0.36. High-quality reads were aligned to the *Mus musculus* mm10 primary assembly genome using CLC Genomics Workbench. Uniquely mapped reads aligned to exons were counted within the same tool and analyzed for differentially expressed genes (DEGs) using the DESeq2 R package v1.44.0. Genes with a false discovery rate (FDR)-adjusted *P*-value < 0.01 were considered DEGs. The sequencing data (FASTQ files) associated with this study will be deposited in the Gene Expression Omnibus (GEO). PCA plots were generated using ggplot2 4.0.0 and volcano plots using EnhancedVolcano v1.22 in R. Figures were generated in Adobe Illustrator.

### Spatial mass spectrometry imaging (SMI)

Spatial mass spectrometry imaging (SMI) was carried out as described previously. ^40^ Frozen heart tissues were sectioned at 12 μm thickness using an Epredia NX50 cryostat (Epredia, Kalamazoo, MI); one section was collected onto a slide for mass spectrometry imaging while a serial section was collected for H&E staining. Digital images of the stained sections were collected using a Hamamatsu NanoZoomerSQ digital slide scanner (Hamamatsu Photonics, Bridgewater, NJ). The H&E images were annotated to indicate regions corresponding to coronary arteries in the heart section. GIMP was used to co-register the annotated images to optical images of the serial unstained sections for SMI that had been acquired at 4800 dpi using an Epson Perfection V600 flatbed document scanner (Epson US, Los Alamitos, CA). Heart sections were coated with 10 mg/mL 1,5-diaminonaphthalene in 50% acetonitrile using an HTX M5 Robotic Reagent sprayer over five passes. The following spray parameters were employed: a nozzle temperature of 75 °C, a track speed of 1200 mm/min, a flow rate of 100 μL/min, a track spacing of three mm, a nitrogen pressure of 10 psi, and nozzle height of 40 mm. SMI was performed on a Bruker timsTOF fleX QTOF mass spectrometer (Bruker Daltonics, Billerica, MA) in negative ion mode at 20 μm resolution using FlexImaging 7.0. The co-registered images were used for fiducial alignment to the slide and data was only collected from the annotated vessels. Instrument parameters in timsControl 3.1 were as follows: *m/z* range of 50-1000, 200 laser shots per pixel, a Funnel 1 RF of 165.0 Vpp, a Funnel 2 RF of 190.0 Vpp, a Multipole RF of 215.0 Vpp, a Collision Energy or 5.0 eV, a Collision RF of 575.0 Vpp, a Transfer Time of 70.0 μs, a Pre Pulse Storage of 6.0 μs. Data was visualized using SCiLS Lab Pro 2025a (Bruker). Peaks were manually selected using an integration window of ± 15 ppm. Peaks were putatively identified using the MetaboScape 2025 (Bruker) plugin for SCiLS allowing a 10 ppm mass tolerance.

### Spatial-scRNA-seq

For spatial transcriptomics, transverse formalin-fixed, paraffin-embedded (FFPE) mouse heart tissue sections (5 μm) were processed using the 10x Genomics Visium CytAssist Spatial Gene Expression platform, following manufacturer’s instructions. H&E-stained images were tiled using FIJI (ImageJ) and aligned with CytAssist output images in the 10x Genomics Loupe Browser (v7) to generate JSON alignment files. These JSON files were paired with FASTQ files for downstream processing with the 10x Space Ranger pipeline. RNA quality and library construction quality control were performed with Agilent 4150 TapeStation system. Libraries were barcoded and sequenced on Illumina NovaSeq platform and demultiplexed at the University of Rochester Genomics Research Center. FASTQ files were fed into 10x Genomics’ Cell Ranger (for FLEX libraries) and Space Ranger (for Visium libraries) pipelines with default parameters.

For the scRNA-seq assay, transverse sections of FFPE mouse heart tissue sections from *Lmod1^WT^* and *Lmod1^SMKO^* mice treated nine weeks with PCSK9/HFD, were processed to liberate individual cells with dissociation enzyme mix containing Liberase TH. Dissociated cells were then loaded into the 10x Genomics Chromium X system using the Fixed RNA Profiling FLEX kit (Singleplex protocol) to generate Gel Beads-in-Emulsion (GEMs), following manufacturer’s instructions. To target ∼10,000 cells per sample, eight tissue sections at 50 μm thickness were collected per genotype. Between ∼5,000-7,000 cells were captured per sample, with mean reads per cell ranging from ∼45,000-65,000 and a median of ∼1,300-1,800 genes per cell. Mapping rates ranged from ∼90%-91.5%, and total reads per sample ranged from ∼492,000,000-494,000,000. For spatial libraries, the number of captured 55 μm voxels ranged from ∼2,200-2,400, with a mean of ∼207,000 reads per voxel and a median of ∼1,000 genes per voxel. Mapping rates ranged from 90%- 95%, with total reads per sample ranging from ∼469,000,000-497,000,000. Other library metrics are available upon request.

### Next generation library sequencing

For scRNA-seq data, initial processing with Seurat v4 ^41^ revealed technical artifacts consistent with ambient RNA contamination, which were removed using CellBender. ^42^ The CellBender-filtered .h5 file was reformatted using ptrepack from PyTables to match the structure of a Cell Ranger v3 .h5 file to load into Seurat using Read10x_h5(). Downstream scRNA-seq analysis was performed with Seurat v4, including quality control, clustering, cell-type annotation based on known markers, and use of Seurat’s ‘anchor based’ approach to integrate samples and account for potential batch effects. For pathway analysis, fully annotated Seurat object was used as input for single cell pathway analysis (SCPA), a specific package tailored to handle noise inherent in scRNA-seq datasets. ^43^ Gene Ontology Biological Process (GOBP) database was queried for SCPA pathway analysis.

For spatial analysis, initial processing was done with Seurat v4. To query spatial voxel spots specific to coronary arteries, barcodes from *Myh11*-positive libraries were obtained by use of 10X Genomics Loupe Browser (v7). These coronary artery-specific barcodes were then used to enrich for coronary-specific spatial voxels via Seurat’s subset() function and subsequently integrated using Seurat’s ‘anchor based’ approach. The integrated object was then normalized and processed for downstream differential gene expression analysis. To deconvolute spatial voxels, Seurat’s ‘anchor’-based integration workflow was used to map cell types onto spatial assay in a genotype-matched manner. Briefly, spatial and scRNA-seq modalities were processed separately. For spatial assay, library coordinates were extracted and then added to Seurat object metadata to enable downstream spatial plotting. To restrict deconvolution to spatial regions of interest, coronary artery-associated voxel spots were enriched using Seurat’s subset() function. To aid in generating more accurate cell type predictions, clusters in scRNA-seq reference were reorganized prior to integration. One dominant cardiomyocyte cluster was removed, and SMCs and pericytes were grouped into a mural cluster to improve the quality of the resulting ‘predictions.assay’. Modality integration of reorganized scRNA-seq reference- and spatial-assays was performed with Seurat’s ‘FindTransferAnchors()’ function, generating a ‘predictions.assay’. As Seurat’s ‘SpatialFeaturePlot()’ function displays only one predicted cell at a time, data from ‘predictions.assay’ were extracted and reformatted for visualization using tidyr and scatterpie R packages to generate composite pie chart representations of multiple cell types per spatial voxel. Spatial and single-cell transcriptomics data will be available in NCBI’s GEO upon acceptance of the manuscript.

### Statistical analyses

More than 300 mice were studied over the course of three years in more than 10 separate cohorts. Based on early studies, there was no need to perform normality tests for most animal studies given the binary nature of the CAD phenotype (ie, no disease in controls). Where multiple groups were analyzed (eg, Western blotting or Lentiviral shRNA transductions), a one-way ANOVA was performed followed by a post-hoc Tukey’s test. When only two groups were compared, statistical significance was determined using a two-tailed t-test (unpaired or paired, as specified). An *a priori* statistical measure of significance was set at 0.05. Analyses were performed using SPSS® software V.29 (Chicago, IL) and GraphPad Prism (versions 9-10; GraphPad Software, La Jolla, CA, USA). The latter was also used for creating statistical illustrations. Some histological and immunofluorescence experiments were quantified by two independent investigators who were blinded to the experimental design as indicated in the figure legends with inter-observer variances not exceeding 10%.

## Results

### LMOD1 is enriched in SMCs of human coronary artery and reduced with atherosclerosis

We probed previous scRNA-seq data ^7^ and found abundant expression of *LMOD1* mRNA in SMC and pericyte clusters of human coronary artery, with lower levels in modulated SMCs and fibroblasts, and barely detectable expression in endothelial cell and immune cell clusters (**Figure S1A, S1B**). An analysis of scATAC-seq data ^27^ showed a similar trend in chromatin accessibility, with most *LMOD1* gene activity clustered in human coronary artery SMCs (**Figure S1C-E**). Consistent with these genomic findings, confocal immunofluorescence microscopy (CIFM) demonstrated enriched LMOD1 protein within ACTA2-positive medial SMCs of human coronary arteries, with much lower levels in overlying atheromatous plaque (**Figure S2**). Collectively, these results are consistent with the preferential expression of LMOD1 in differentiated SMCs ^30,44^ and its attenuated expression in human atherosclerotic lesions. ^13,27,45^

### *Itga8-CreER^T2^* prevents lethal intestinal myopathy caused by *Lmod1* loss

Global *Lmod1* loss-of-function through CRISPR-mediated disruption of exon one, or installation of pathologic substitutions that result in a premature termination codon in exon two, results in neonatal death due to visceral myopathy of the intestine and bladder (**Figure S3A, S3B**). ^15^ Thus, defining chronic vascular phenotypes in global *Lmod1* null mice is impossible. Accordingly, the promoter containing two SRF-binding CArG boxes ^30^ and first exon of *Lmod1* were floxed for conditional gene loss experiments (**Figure S3C**). Mice homozygous for Lox*P* showed no change in expression of *Lmod1* mRNA or LMOD1 protein suggesting intact regulatory elements in the *Lmod1* promoter and intronic region (**Figure S3D, S3E**). A comparable loss of aortic LMOD1 protein was observed upon *Myh11-CreER^T2^-* or *Itga8-CreER^T2^*-mediated recombination of the floxed *Lmod1* gene (**Figure 1A, 1B**), with an estimated recombination efficiency of ∼80% using *Itga8-CreER^T2^* (**Figure S3F**). Quantitative RT-PCR revealed the conditional knockout of *Lmod1* to be a true null allele (**Figure S3G**). ^46^ Tamoxifen-treated *Lmod1^KO-Myh^*^11^ mice showed loss of LMOD1 protein in intestinal SMC (**Figure 1A, 1C**) and exhibited distension of the intestine leading to rapid death within a week of the last dose of tamoxifen (**Figure 1D, left; Figure S3H**). Moreover, all oil-treated *Lmod1^KO-Myh11^* mice died by 12 weeks of age, likely from cumulative unprogrammed recombination (**Figure 1D, left**). ^20,47^ In contrast, tamoxifen-treated *Lmod1^KO-Itga8^* mice showed normal levels of LMOD1 protein in the intestine (**Figure 1A, 1C**), and the majority lived for at least six months (**Figure 1D, right**). LMOD1 protein levels were reduced by more than 90% in the aorta of *Lmod1^KO-Itga8^* mice after six months of Tmx administration with little variation in TAGLN or ITGA8 protein (**Figure 1E, 1F**) and limited evidence of intestinal pathology (**Figure S3H**).

**Figure 1.**
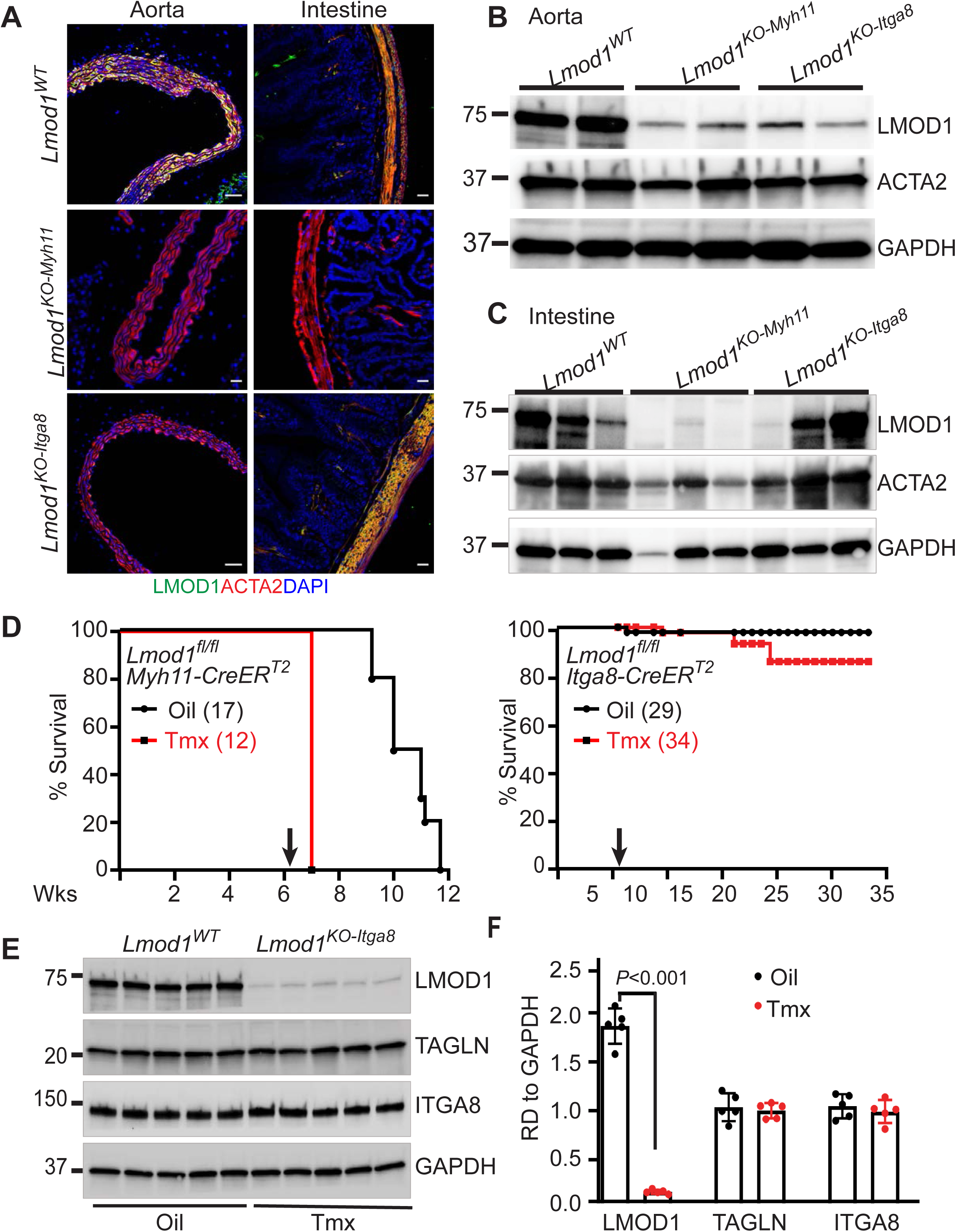
Conditional knockout of *Lmod1* with *Myh11-CreER^T2^* vs *Itga8-CreER^T2^*. (**A**) Confocal immunofluorescence microscopy (CIFM) of LMOD1 and ACTA2 in aorta and intestine of the indicated mice. Scale bars are 20 μm. Western blotting for LMOD1 in aorta (**B**) and intestine (**C**) of indicated mouse genotype. (**D**) Survival curves for Oil versus Tamoxifen (Tmx) treated floxed *Lmod1*/*Myh11-CreER^T2^* (left) and floxed *Lmod1*/*Itga8-CreER^T2^* (right) mice. The black arrow indicates the time of Oil or Tmx administration. Western blotting (**E**) and quantitation (**F**) of indicated proteins after six months of Oil or Tmx administration; n=5 independent mice per genotype (*Lmod1^WT^* is *Lmod1^fl/fl^*).

Females deficient in *Lmod1* had little change in body weight over a six-month period, but males began to slow in weight at week 31 (**Figure S3I**). Both male and female *Lmod1^KO-Itga8^* mice showed a reduction in systolic blood pressure versus oil-administered controls (**Figure S3J**). Interestingly, serum levels of bacteria-derived lipopolysaccharide were elevated in *Lmod1^KO-Myh11^* mice, but not in *Lmod1^KO-Itga8^* mice, suggesting sepsis-mediated death in the former animals (**Figure S3K**). Taken together, these findings extend previous reports demonstrating evasion of lethal visceral myopathies using *Itga8-CreER^T2^*, thus allowing for exploration of vascular phenotypes that otherwise would not be possible. ^20–23^

### *Lmod1^SMKO^* mice have enlarged blood vessels and reduced baseline blood pressure

After only 10 days of Tmx administration, the aorta of *Lmod1^KO-Itga8^* mice (hereafter abbreviated as *Lmod1^SMKO^*) was measurably larger than *Lmod1^WT^* controls (carrying *Itga8-CreER^T2^* and receiving Tmx), from ascending to abdominal aorta (**Figure S4A-S4C**). Further, compared to *Lmod1^WT^* mice, the carotid artery of *Lmod1^SMKO^* mice was enlarged (**Figure S4D**) with a significant increase in circumference and an attending decrease in medial thickness (**Figure S4E**). We next interrogated the hearts from apex to base (**Figure S4F**) and found a significant increase in the circumference of coronary arteries of *Lmod1^SMKO^* mice (**Figure S4G, S4H**). The attenuated systolic blood pressure seen at six months post-Tmx (**Figure S3J**) was observed as early as 10 days post-Tmx in both male and female *Lmod1^SMKO^* mice (**Figure S4I**). Despite the above morpho-physiological changes in the vasculature and a significant decrease in LMOD1 protein in the aorta of *Lmod1^SMKO^* mice, there was no significant reduction in several SMC contractile proteins, including MYH11, ACTA2, and TAGLN (**Figure S4J, S4K**). These findings established a baseline hypotensive phenotype with outward vascular remodeling and normal SMC contractile protein expression in *Lmod1^SMKO^* mice.

### *Lmod1^SMKO^* mice exhibit angiotensin II-independent occlusive CAD after an atherogenic diet

Based on the enlarged vessel phenotype, we surmised *Lmod1^SMKO^* mice would be susceptible to aneurysm formation following a PCSK9/high fat diet (HFD)/angiotensin II regimen (**Figure S5A**). All mice exhibited similar hypercholesterolemia and body weights (**Figure S5B, S5C**), but only *Lmod1^WT^* mice showed evidence of abdominal aneurysm (**Figure S5D**). Although there was some death in the *Lmod1^WT^* control arm, far more *Lmod1^SMKO^* mice died following the combined atherogenic/angiotensin II regimen (**Figure S5E**), despite showing no evidence of cardiac hypertrophy (**Figure S5F**) or cardiac fibrosis (**Figure S5J**). However, gross inspection of the heart revealed a tofu-like appearance of the coronary vasculature in *Lmod1^SMKO^* mice (**Figure S5G**). Histological analysis demonstrated occlusive CAD with marked fibrous caps (**Figure S5H, S5I**) and lipid droplet formation (**Figure S5K**) in both males and females (below). In no instance were any of these coronary artery phenotypes manifested in *Lmod1^WT^* mice.

To ascertain whether the CAD phenotype in *Lmod1^SMKO^* mice required angiotensin II, we conducted a study in the same manner as above only with no angiotensin II (**Figure 2A**). All mice had comparable hypercholesterolemia at 11.5 weeks of age or ∼2 weeks post-PCSK9/HFD (**Figure 2B left**), and the levels of cholesterol, triglycerides, LDL, and HDL were similar between genotypes at the termination (16 weeks of age) of the study (**Figure 2B right, 2C**). *Lmod1^SMKO^* mice had ∼35% mortality over the course of the study (**Figure 2D**), considerably less than that seen with angiotensin II (**Figure S5E**). Blood cell and blood chemistry measures revealed higher circulating lymphocyte and lower neutrophil counts as well as a decrease plasma chloride level in *Lmod1^SMKO^* mice (**Table S4**). ORO staining of the aorta was greater in *Lmod1^SMKO^* mice than *Lmod1^WT^* mice (**Figure 2E, 2F**). As with the experiment above, the atherogenic regimen provoked a tofu-like appearance of coronary arteries in *Lmod1^SMKO^* mice, with no such appearance in any of the *Lmod1^WT^* mice (**Figure 2G**). Histological staining revealed occlusive coronary lesions in *Lmod1^SMKO^* mice (**Figure 2H**) with evidence of a fibrous cap as shown by ACTA2 staining (**Figure 2I**). CIFM consistently demonstrated virtually no detectable LMOD1 staining in the coronary arteries of *Lmod1^SMKO^* mice (**Figure 2I; Figure S6**). To address whether the CAD phenotype was unique to the PCSK9/HFD model, we crossed the floxed *Lmod1* mouse with *Apoe* null mice to generate *Lmod1*/*Apoe* double knockouts and subjected these mice to either Tmx or Oil treatment, followed by a WD for 11 weeks. The results of this study showed similar tofu-like coronary arteries in *Lmod1*/*Apoe* double knockout mice (not observed in any oil control mice). Histological sectioning revealed occlusive lesions closely resembling those seen in *Lmod1^SMKO^* mice subjected to a PCSK9/HFD regimen (**Figure S7**). Taken together, these findings demonstrate an angiotensin II independent-occlusive CAD phenotype in *Lmod1^SMKO^* mice subjected to two distinct atherogenic models.

**Figure 2.**
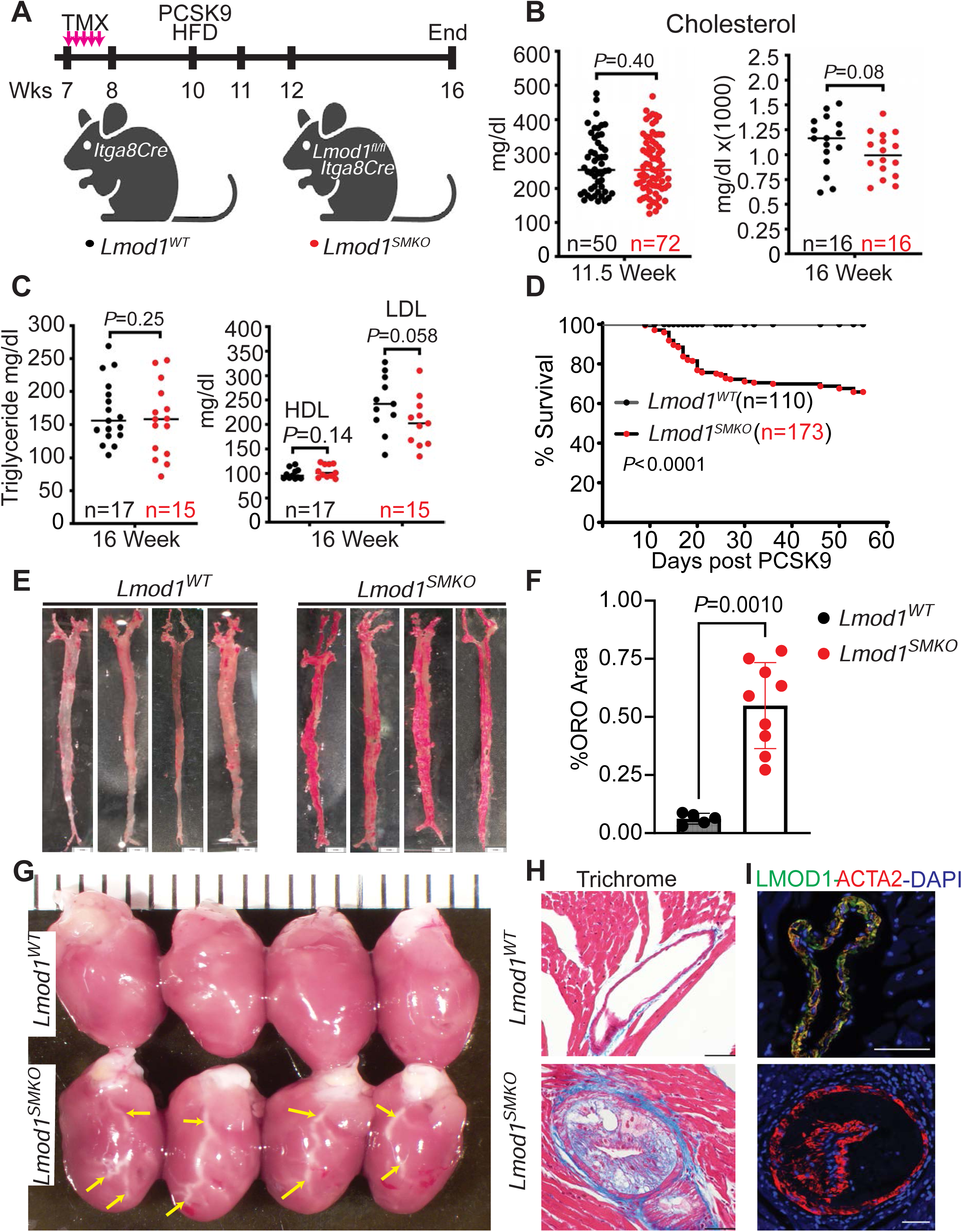
AngII-independent CAD phenotype in *Lmod1^SMKO^*mice. (**A**) Experimental study starting at seven weeks of age. Beginning here, all *Lmod1^WT^* mice carried the *Itga8-CreER^T2^* allele and were treated with the same atherogenic regimen as *Lmod1^SMKO^* mice. (**B**) Total serum cholesterol in *Lmod1^WT^* (black dots) and *Lmod1^SMKO^* (red dots) mice at indicated ages. (**C**) Total serum triglycerides and HDL and LDL. (**D**) Cumulative survival curves for *Lmod1^WT^* and *Lmod1^SMKO^* mice across multiple cohorts. Oil-Red-O (ORO) staining (**E**) and quantitation of percent ORO staining (**F**) across the aorta of *Lmod1^WT^* (n=5) and *Lmod1^SMKO^* (n=9) mice. (**G**) Gross cardiac images showing tofu-like appearance of coronary arteries (yellow arrows) in *Lmod1^SMKO^* mice. Masson trichrome staining (**H**) and CIFM imaging of LMOD1 and ACTA2 protein (**I**) in sections of coronary artery from *Lmod1^WT^* (top panels) and *Lmod1^SMKO^* mice (bottom) under the PCSK9/HFD regimen for 52 days. Note the prominent ACTA2 positive fibrous cap in *Lmod1^SMKO^* section. Scales bars, 50 μm.

### CAD lesions in *Lmod1^SMKO^* mice are heterogeneous and manifested early following PCSK9/HFD

Time course studies were performed to define the onset of CAD and the physicochemical properties of lesions under a PCSK9/HFD regimen. Mild ORO-positive lesions could be readily detected in *Lmod1^SMKO^* after only eight days of the PCSK9/HFD regimen (**Figure 3A-3C; Figure S8**). After 16 days, H&E and ORO+ lesions were larger (**Figure 3D-3F; Figure S8**) and by 63 days, lipid-laden, nearly occluded coronary lesions were evident with a prominent fibrous cap as revealed by Masson trichrome staining (**Figure 3G-3I; Figure S8**). The latter finding was further substantiated by quantitative picrosirius red staining (**Figure S9A-9E**). Cumulative quantitative measures of ORO area (**Figure 3J**), plaque area (**Figure 3K**), and coronary circumference (**Figure 3L**) all were highly significant in *Lmod1^SMKO^* mice. Of note, ORO occlusive lesions were occasionally observed in small (40 μm) coronary arterioles but were infrequent in larger coronary arteries (>150 μm) at the base of the heart (**Figure S10**). The latter may explain why no evidence of myocardial infarction was observed in >100 *Lmod1^SMKO^* mice analyzed. The necrotic core of advanced lesions occupied ∼35% of plaque area (**Figure S11**); however, while human coronary atheroma showed obvious calcification, no such change was observed in *Lmod1^SMKO^* mice under the experimental conditions reported (**Figure S9F, 9G**). Surprisingly, despite clear evidence of a loss in LMOD1 staining, atherosclerosis was rarely observed in the vasculature of brain, kidney, liver, lung, mesentery, and spleen (**Figure S12**).

**Figure 3.**
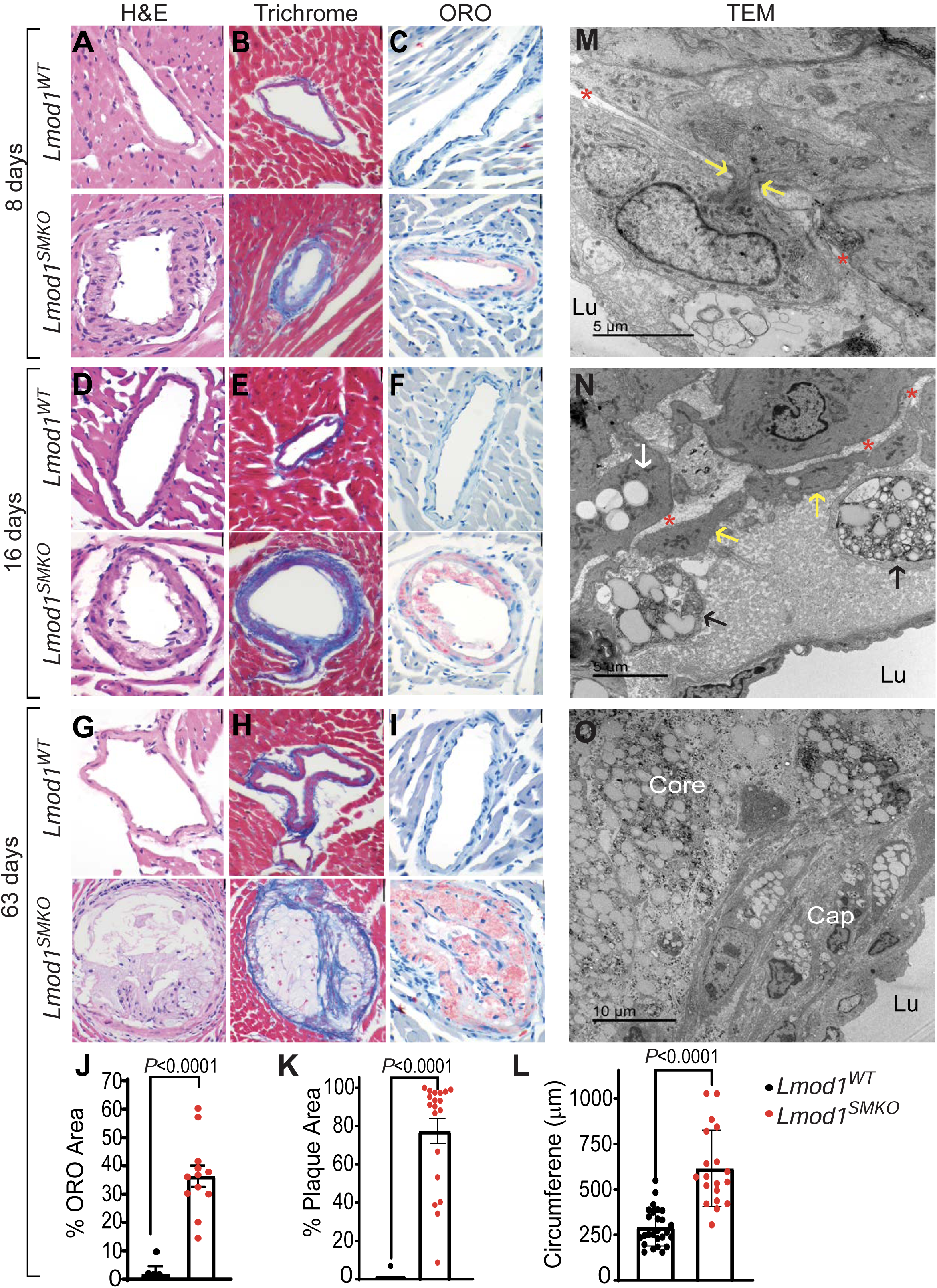
Time course of coronary lesion development in *Lmod1^SMKO^* mice. Representative staining of coronary vessels from *Lmod1^WT^* (n=5 females and 10 males) and *Lmod1^SMKO^* (n=6 females and 11 males) mice after 8 (**A-C**), 16 (**D-F**), and 63 (**G-I**) days of PCSK9/HFD. Quantitative measures of ORO staining (**J**), plaque area (**K**), and circumference (**L**) of coronary arteries from *Lmod1^WT^* and *Lmod1^SMKO^* mice after 63 days of PCSK9/HFD. Data in panel **J** represents 10 and 12 coronary arteries from 5 *Lmod1^WT^* and 5 *Lmod1^SMKO^* mice, respectively. Data in panels **K** and **L** represent 26 and 18 coronary arteries from 5 *Lmod1^WT^* and 5 *Lmod1^SMKO^* mice, respectively. Scales bars (located in upper right corner of each panel) are 20 μm save lower panel **H** (50 μm). Transmission electron micrographs (TEM) showing a presumptive medial SMC migrating through a fenestra (yellow arrows) of the IEL (marked in red asterisks) at 8 days (**M**); two intimal foam cells (black arrows), two recently migrated intimal SMC (yellow arrows, one of which has a lipid droplet), and a medial SMC foam cell (white arrow) at 16 days (**N**); and a more complex plaque with lipid rich foam cell core and foam cell-containing fibrous cap at 63 days (**O**). See **Figures S15-S17** for additional electron micrographs of *Lmod1^WT^* and *Lmod1^SMKO^* coronary arteries. Lu, lumen.

There was no obvious bias for atheromatous lesions between right, left, or septal coronary artery of *Lmod1^SMKO^* mice and no sexual dimorphism after nine weeks of PCSK9/HFD (**Figure S13; Table S5**). Next, we adopted the Stary classification ^37^ of atherosclerotic lesion types in a comprehensive and blinded evaluation of >500 coronary sections across 116 mice of equal sex. This analysis demonstrated a predominance of Type I and Type II lesions at early time points, with a transition to predominantly Type IV and Type V lesions at later stages of the PCSK9/HFD regimen in *Lmod1^SMKO^* mice (**Figure S14; Table S6**). Only 3/161 coronary sections from *Lmod1^WT^* mice showed Type I lesions with no other advanced lesions recorded, even as late as 203 days following the start of PCSK9/HFD (**Figure S14; Table S6**).

On the other hand, essentially all coronary artery sections from *Lmod1^SMKO^* mice showed lesions of Types I-V after 100 days of PCSK9/HFD (**Figure S14; Table S6**). Notably, there were no complex Type VI lesions, consistent with the absence of myocardial infarction in this coronary atherosclerosis model.

Finally, while Type I-III lesions were approximately equal between sexes, females showed more Type IV lesions and males more Type V lesions at later time points of the experiment (**Table S6**). Taken together, these results demonstrate a highly penetrant, coronary-specific occlusive atherosclerosis phenotype, of varying physicochemical nature, in both male and female *Lmod1^SMKO^* mice, with the earliest lesions manifested as early as eight days following a PCSK9/HFD regimen.

### Coronary ultrastructure in *Lmod1^SMKO^* mice reveals phenotypically diverse VSMCs

Surprisingly, few comprehensive studies exist on the ultrastructure of mouse coronary arteries. ^48,49,50^ We found that the ultrastructure of coronary arteries from *Lmod1^WT^* mice was indistinguishable from those of wild type chow-fed mice, with both having abundant myofilaments, dense bodies, myoendothelial junctions, and no lipid droplets (**Figure S15**). In contrast, coronary arteries from *Lmod1^SMKO^* mice showed presumptive SMCs migrating through the fenestra of the IEL into a thickened intima as early as eight days post PCSK9/HFD (**Figure 3M; Figure S16**). As shown below, these migrating cells were proven to be SMC using lineage tracing. By 16 days, the intima was thicker with medial and intimal SMC (defined by dense bodies) containing lipid droplets; the intimal SMC were found below presumptive macrophage-derived foam cells at this stage (**Figure 3N**). At later stages of coronary atherogenesis, plaques were progressively larger with extracellular lipid, cholesterol crystals, and an increasing number of SMC-derived foam cells in the intima and media (**Figure 3O; Figure S17**).

Extensive profiling revealed considerable heterogeneity in SMC lipid droplets, suggesting active lipid metabolism (**Figure S17**). Indeed, SMI profiling revealed a pronounced increase in an array of lipid-derived molecules, including sulfated cholesterol, ceramides, and sphingomyelin species within atheromatous coronary arteries of *Lmod1^SMKO^* mice (**Figure S18; Table S7**). Taken together, these results suggest metabolically active, migrating coronary SMCs in *Lmod1^SMKO^* mice that contribute to the foam cell population in an evolving coronary atheroma.

### Coronary artery SMC of *Lmod1^SMKO^* mice contribute to atheromatous plaque formation

Previous lineage tracing studies have documented a variable number of SMC-derived foam cells in atherosclerotic lesions of the aorta and brachiocephalic artery.^8,9,51^ However, to our knowledge, no lineage tracing has been reported in the adult coronary vasculature. Accordingly, *Lmod1^WT^* and *Lmod1^SMKO^* mice (carrying the *mTmG* reporter) underwent the PCSK9/HFD regimen and coronary arteries analyzed by CIFM for the presence of SMC-derived GFP positive cells in atheromatous plaques. As expected, *Lmod1^WT^* coronaries showed no evidence of disease and all GFP+ SMCs were confined to the media (**Figure 4A, 4B**). In contrast, SMC-derived GFP+ cells were found in coronary plaques of *Lmod1^SMKO^* mice (**Figure 4D, 4E, 4G, 4H**). To accurately quantify the percentage of GFP+ plaque-derived cells, we developed a novel immunogold electron microscopic lineage tracing assay (IEMLT).

**Figure 4.**
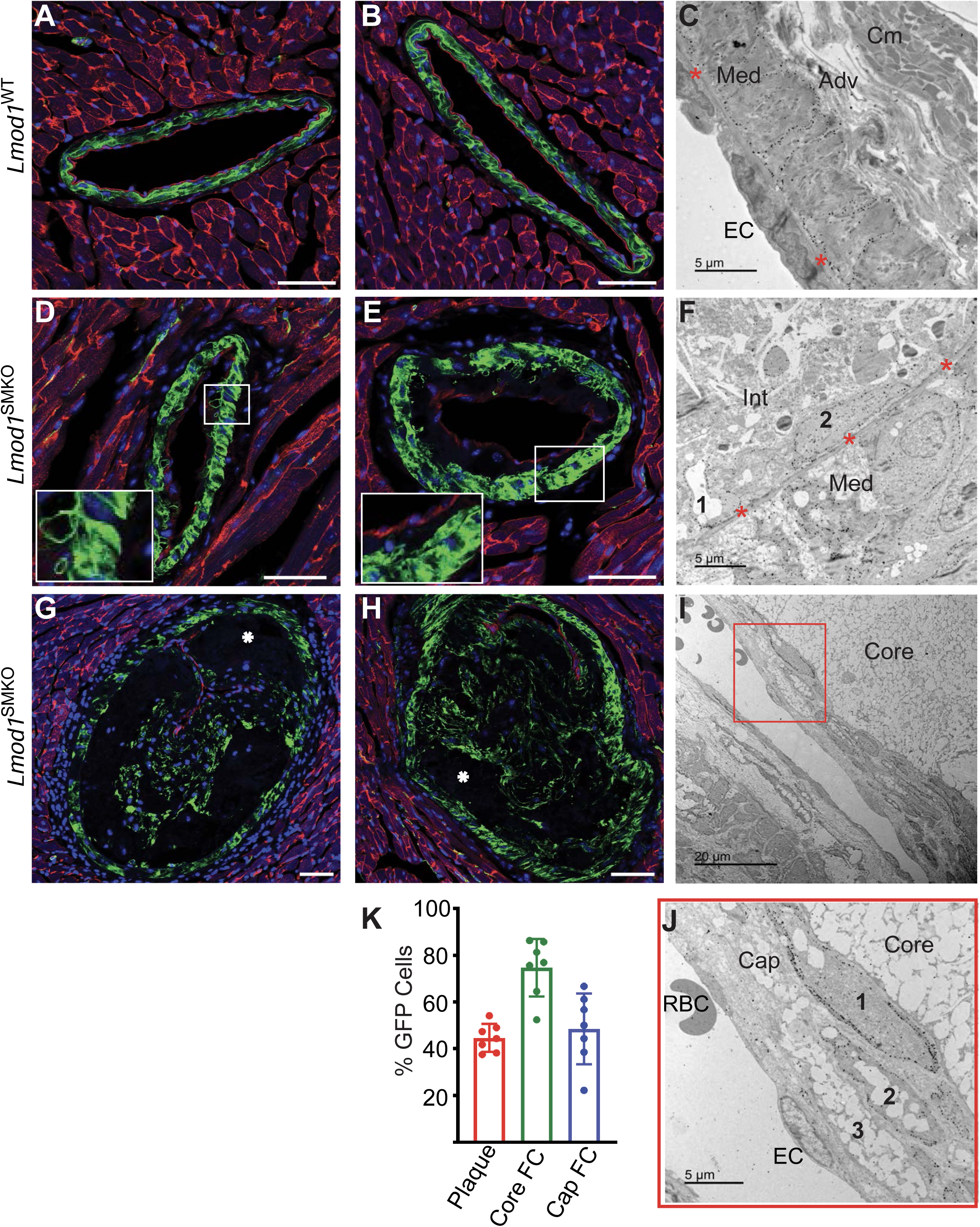
Coronary artery SMC lineage tracing in *Lmod1^SMKO^*mice. *Lmod1^WT^* (**A-C**) and *Lmod1^SMKO^* (**D-J**) coronary arteries from mice carrying the membrane tomato/membrane GFP (mTmG) reporter. Red fluorescence (**A,B,D,E,G,H**) indicates surrounding cardiomyocytes and endothelial and immune cells as well as fibroblasts whose mTmG reporter did not undergo recombination (i.e., *Itga8-CreER^T2^* was inactive in these cells). Conversely, green fluorescence represents medial and plaque cells of SMC origin (i.e., *Itga8-CreER^T2^* was active in these cells and recombined out the tomato reporter allowing for membrane GFP expression). Note GFP+ cells beginning to populate early atheroma (**D,E**; see also, **Figure S20E-S20F**) and comprising a large portion of advanced plaques (**G,H**). White asterisks indicate necrotic core. Immunogold electron microscopy lineage tracing (IEMLT) of a *Lmod1^WT^* control coronary artery (**C**); an *Lmod1^SMKO^* coronary from 16-day lesion with two intimal core-centric GFP+ cells (labeled 1 and 2) (**F**); and an advanced, 63-day lesion showing several GFP+ cells at the fibrous cap (I). (**J**) Higher magnification of red boxed region in panel **I** showing three fibrous cap cells (labeled 1-3), two of which (1 and 2) are GFP+. (**K**) Percentage of GFP+ cells in plaques of seven independent *Lmod1^SMKO^* coronaries (red bar) and the percentage of Core (green bar) and Cap (blue bar) derived foam cells that were GFP+. Scale bars are 50 μm. Adv, adventitia; CM, cardiomyocyte; EC, endothelial cell; Int, intima; Med, media; RBC, red blood cell. See Supplemental Figures 19 **and 20** for additional IEMLT images.

Extensive testing of IEMLT in control mice showed unambiguous and specific labelling of medial coronary artery SMC membranes (**Figure 4C**; **Figure S19**). The IEMLT method demonstrated obvious distinction between GFP+ and GFP-cells (**Figure 4F, 4I, 4J; Figure S20A-20F**), with GFP+ SMCs migrating through the IEL as early as six days post-PCSK9/HFD (**Figure S20E-20F**). Rigorous quantitation of cells within advanced atheromatous lesions revealed >40% of plaque cells to be of coronary SMC origin (**Figure 4K**). Interestingly, 75% of GFP+ cells in the lower core of lesions were foam cells (**Figure 4K; Figure S20A, S20C**) and a surprising 50% of GFP+ cap cells also contained lipid droplets (**Figure 4I-4K; Figure S20B, S20D**). These results provide the first lineage tracing analysis of coronary atherosclerosis, further corroborating a primary role of medial-derived SMCs in atherogenesis.

SMCs undergo diverse fate and state transitions during atherogenesis. ^10,11,52,53^ CIFM colocalization of GFP+ cells with markers of previously described SMC phenotypic states showed a very small percentage of coronary plaque-derived SMCs expressing CD68, SPP1, or LGALS3 (**Figure S21**). Further, despite carefully controlled studies, no proliferating GFP+ cells were found within coronary plaques at early or late stages of atherogenesis; however, as expected, TUNEL positive cells (GFP-) were found in the necrotic core of some plaques (**Figure S22**). These results suggest that, within the context of experiments here, the majority of coronary SMC-derived plaque cells exhibit little, if any, proliferation or apoptosis.

### Reduced LMOD1 does not confer CAD in mice

Two single nucleotide variants (SNVs) within the first intron of human *LMOD1* have been associated with vascular disease, including CAD, ^5,54–57^ and one of them (rs34091558) has been validated in vitro. ^58^ We used a two-component CRISPR approach ^35^ to delete a portion of intron 1 in *Lmod1* encompassing the orthologous intronic SNV corresponding to rs34091558 (**Figure 5A**). Positive founders were bred for germline transmission, and mice homozygous for the 11.35 kilobase deletion were overtly normal despite a ∼50% reduction in LMOD1 protein as measured by Western blotting (**Figure 5B**) and CIFM (**Figure 5C**). A similar reduction in LMOD1 protein was observed in *Lmod1^SM-Het^* mice lacking one *Lmod1* allele (**Figure 5D, 5E**). Interestingly, *Lmod1^SM-Het^* mice showed little evidence of coronary atherosclerosis after nine weeks of PCSK9/HFD, and the minimal ORO staining was no different than that observed in *Lmod1^WT^* mice (**Figure 5F, 5H**) despite an obvious reduction in coronary SMC LMOD1 staining (**Figure 5G, 5I**). Thus, by using *Lmod1^SM-Het^* mice as a proxy for the intronic deletion model, we infer that a 50% reduction in LMOD1 protein is insufficient to cause CAD in mice, at least under the experimental conditions here.

**Figure 5.**
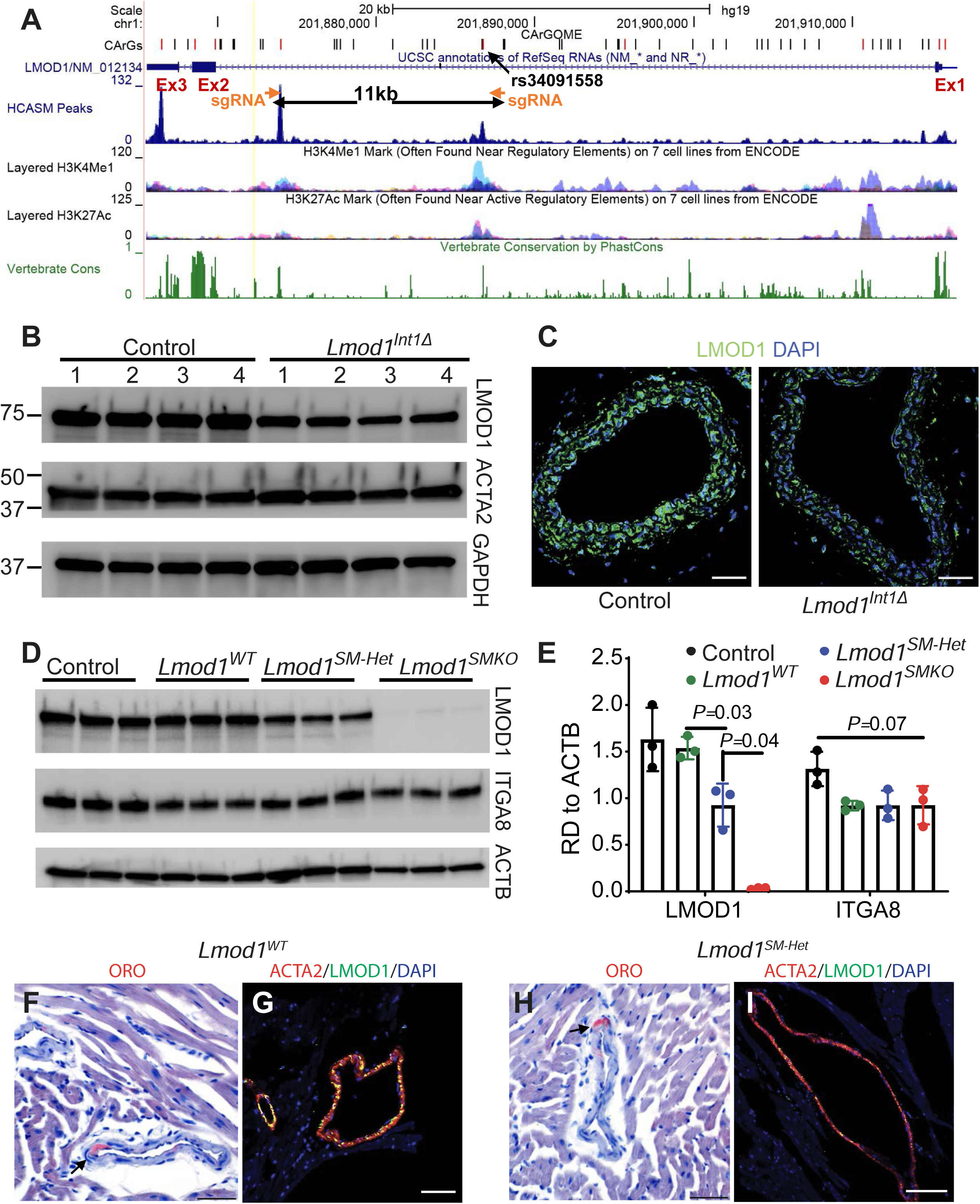
Reduced LMOD1 expression does not confer a CAD phenotype. (**A**) UCSC genome browser screenshot of human *LMOD1* gene structure and position of annotated SNV. The approximate position of sgRNAs used to generate an 11.35 kb intronic deletion in mice is shown. Note that this deletion removes the rs34091558 SNV and an SRF-binding consensus CArG box (indicated by ChIP-seq peak in human coronary artery SMCs and an arrow pointing to a red [conserved CArG] line on the “CArGs track”). This region also coincides with active chromatin marks. Western blot (**B**) and CIFM (**C**) of aortic LMOD1 in mice homozygous for the intronic deletion. Western blot (**D**) and quantitation (**E**) of aortic LMOD1 and ITGA8 in the indicated genotypes. Oil-Red-O staining (**F,H**) and CIFM of LMOD1 (**G,I**) in coronary arteries from *Lmod1^WT^* (**F,G**) and *Lmod1^SMKO^* (**H,I**) mice. Scale bars, 50 μm.

### Bulk, spatial, and scRNA-seq reveal a common pathway of SMC activation in *Lmod1^SMKO^* mice

To gain unbiased insight into coronary SMC states that contribute to the CAD phenotype, we integrated data sets from multiple transcriptomic profiling studies. Bulk RNA-seq with principal component analysis (PCA) of aortic samples revealed distinct transcriptional profiles in *Lmod1^SMKO^* versus *Lmod1^WT^* mice, with divergence further amplified under the HFD/PCSK9 regimen (**Figure S23A, S23B**). Interestingly, the matricellular protein *Thbs1* was upregulated under both conditions (**Figure S23C, S23D; Table S8**). Overrepresentation pathway analysis demonstrated an upregulation of genes associated with microtubule-binding and cell growth under baseline conditions (**Figure S23E**) and an increase in genes linked to lipid biology and vascular stress in the PCSK9/HFD condition (**Figure S23F**).

Next, we obtained comparable cross-sections of the heart from *Lmod1^WT^* and *Lmod1^SMKO^* mice for scRNA-seq and spatial modalities nine weeks after PCSK9/HFD (**Figure 6A**). Ambient RNA removal and object integration with Seurat yielded ten cell types (**Figure 6Bi, 6Bii, Figure S24A-C**) with SMCs accounting for <1% of total cells (96 of 13,925 cells), and cardiomyocytes (CMs) and endothelial cells (ECs) accounting for >60% (**Figure 6Bi**). We focused on coronary arterial (CA) regions by enriching for *Myh11*-positive spatial voxels, with alignment between *Myh11* spatial expression and H&E-defined coronary artery structures confirming accurate CA region capture (**Figure 6Ci**, **Figure 6D**). *Lmod1^SMKO^* hearts possessed a greater CA spatial footprint (∼6.5% for *Lmod1^SMKO^* [128 of 1971 voxels] versus ∼4.3% for *Lmod1^WT^* [998 of 2278 voxels)] (**Figure 6Ci**). Differential expression analysis of *Myh11*-positive CA voxels identified significant up-regulation of *Spp1*, *Lgals3*, and *Thbs1* (**Figure 6Cii**), with *Thbs1* showing coronary artery-enriched expression (**Fig. 6Ciii**).

**Figure 6.**
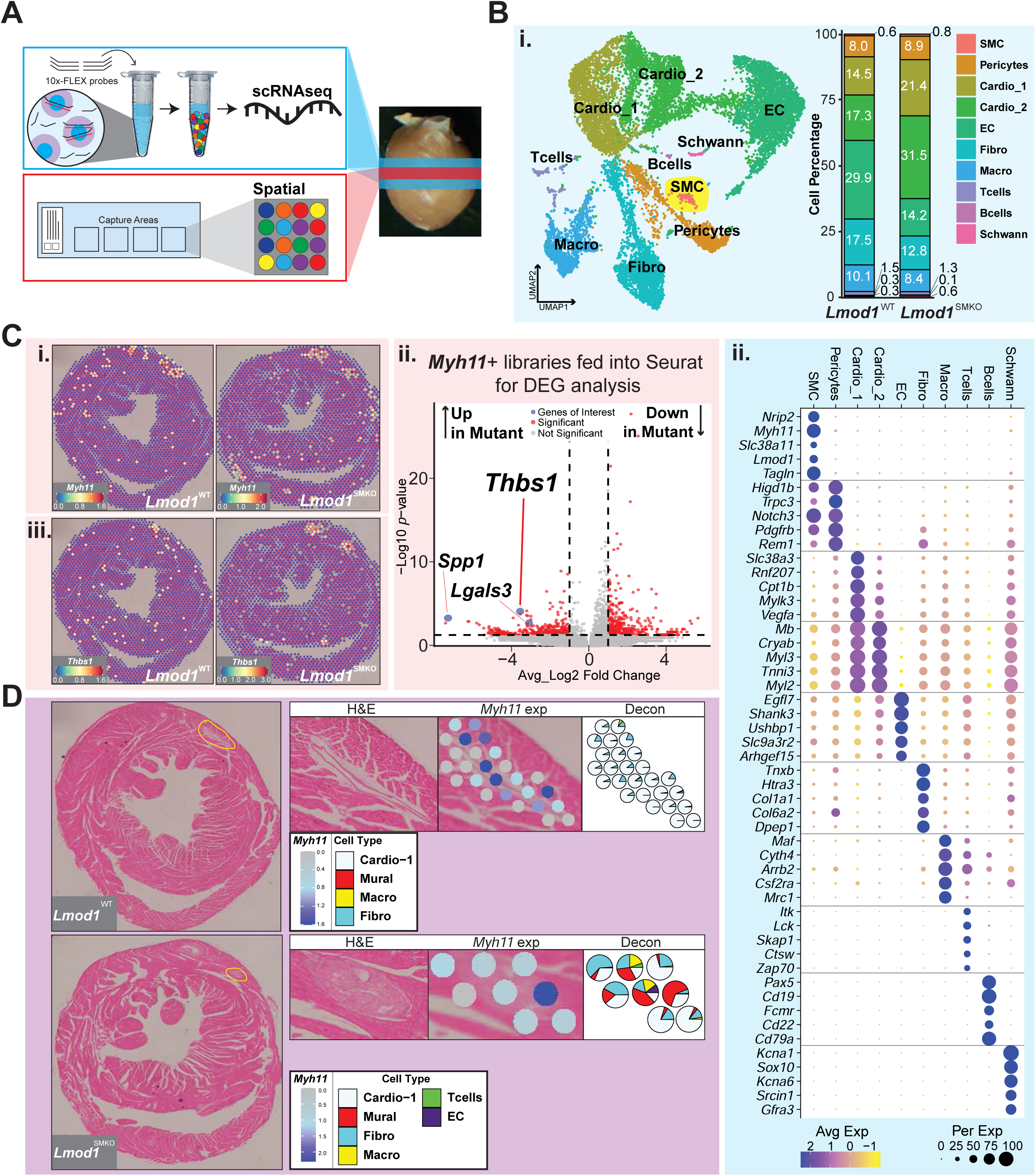
Multi-modal transcriptomic approach to interrogate the coronary artery phenotype in *Lmod1^SMKO^* mice. (**A**) Schematic for scRNA-seq (10x-FLEX, blue) and spatial (10x-Visium, red). Libraries interrogate midline cross-sections of right and left ventricles of mouse hearts. (**B**) scRNA-seq modality: (**i**) UMAP with accompanying stacked bar chart for cell type distribution across *Lmod1*^WT^ and *Lmod1*^SMKO^; and (**ii**) dot plot of top 5 gene markers per cell type. (**C**) Spatial modality: (**i**) spatial feature plot of *Myh11* (used as proxy for coronary artery-enriched spatial libraries); (**ii**) volcano plot of differentially expressed genes (DEG) for *Lmod1*^WT^ and *Lmod1*^SMKO^ *Myh11*+ libraries (cutoffs: average log2 fold change ≥ 1, *p* ≤ 0.05);, (**iii**) spatial feature plot of up-regulated *Thbs1* showing coronary-artery enriched expression. (**D**) Spatial spot deconvolution (purple) reflects the integration of scRNA-seq (red) and spatial (blue) modalities. Coronary artery-specific spatial feature plot of *Myh11* expression with corresponding spatial spot deconvolution with cell type composition (pie-chart) using genotype-matched scRNA-seq reference.

Spatial voxels capture multiple cell types within a defined x-y coordinate, necessitating voxel deconvolution. Given the severe coronary artery phenotype in *Lmod1^SMKO^* mice, the two modalities were genotype-matched, enabling proper genotype context-dependent deconvolution. Representative deconvoluted CA regions, with *Myh11* expression to give CA region context, are shown (**Figure 6D; Figure S24D**). Deconvolution of *Lmod1^WT^* found uniform cellular composition with voxels dominated by CMs, sparse fibroblasts, and low mural (SMC and pericyte) and immune cells (**Figure 6D, Figure S24D**). In contrast, *Lmod1^SMKO^* CA regions exhibited mosaic cell type composition patterns with increased mural cell presence, (>50–60%), reduced CMs (<35%), and increased fibroblasts (∼10–60%), consistent with the elevated fibrosis found in Trichrome stained atheromata (**Figure 2H; Figure S5I; Figure 3H**). Finally, there was an increased presence of immune cells, mostly of the macrophage type (**Figure 6D, Figure S24D**).

### Thrombospondin mediates lipid uptake in SMC of *Lmod1^SMKO^* mice

SCPA of the above scRNA-seq data focused on VSMCs and found enrichment of numerous lipid-associated pathways, including ‘GOBP Regulation of Lipid Transport’, which contains *Thbs1* (**Figure 7A, 7B**), a gene also found up-regulated in *Lmod1^SMKO^* coronary artery voxels in spatial (**Figure 6Cii, 6Ciii**). A dot plot of up-regulated vascular genes from spatial modality (**Figure 6Cii**) revealed *Thbs1* expression localized in SMCs, whereas *Spp1* and *Lgals3* localized to macrophages (**Figure 7C**). We then compared coronary artery spatial modality with two aortic bulk RNA-seq datasets (baseline and PCSK9/high-fat diet), and this analysis revealed consistent up-regulation of *Armc3*, *Igfbp2*, *Cenpf*, *Cdca5*, *Cdk1*, *Slfn9*, *Tns3*, and *Thbs1* in *Lmod1^SMKO^* (**Figure 7D**). These findings led us to prioritize *Thbs1* for further study give its controversial role in atherosclerosis (see Discussion). Accordingly, we assessed THBS1 protein expression in mouse and human coronary atherosclerotic lesions. Lineage tracing in coronary arteries of *Lmod1^SMKO^* mice under PCSK9/HFD showed 10% GFP+/THBS1+ cells of SMC origin (**Figure 7E**). Interestingly, immunogold EM suggested a redistribution of THBS1 protein between *Lmod1^WT^* (diffuse cytoplasmic) and *Lmod1^SMKO^* (membrane-associated) coronary arteries (**Figure S25**). We also observed THBS1 in sections of human coronary artery where little LMOD1 staining could be found (**Figure 7F**). Conversely, LMOD1 expression was most abundant in coronary artery that showed low level THBS1 (**Figure 7F**). This reciprocal pattern of expression was also demonstrated in cultured SMCs from *Lmod1^WT^* and *Lmod1^SMKO^* mice (**Figure 7G**). Further, THBS1 was expressed in the atheroprone inner curvature of the human aortic arch, where intense ORO staining was observed (**Figure 7H, 7I**). The latter result prompted additional in vitro studies with primary mouse aortic SMCs (MASMCs) from *Lmod1^WT^* and *Lmod1^SMKO^* mice. Such cells were primed with TNFα and then treated with brief exposure to oxidized LDL (oxLDL). Consistent with the in vivo VSMC foam cell phenotype, *Lmod1^SMKO^* MASMCs exhibited a threefold increase in oxLDL staining compared with *Lmod1^WT^* MASMCs. (**Figure 7J**). Importantly, this elevation in oxLDL staining was virtually abolished by a shRNA targeting *Thbs1* (**Figure 7K, 7L**). These findings suggest a role for THBS1 in lipid uptake or intracellular trafficking in VSMCs.

**Figure 7.**
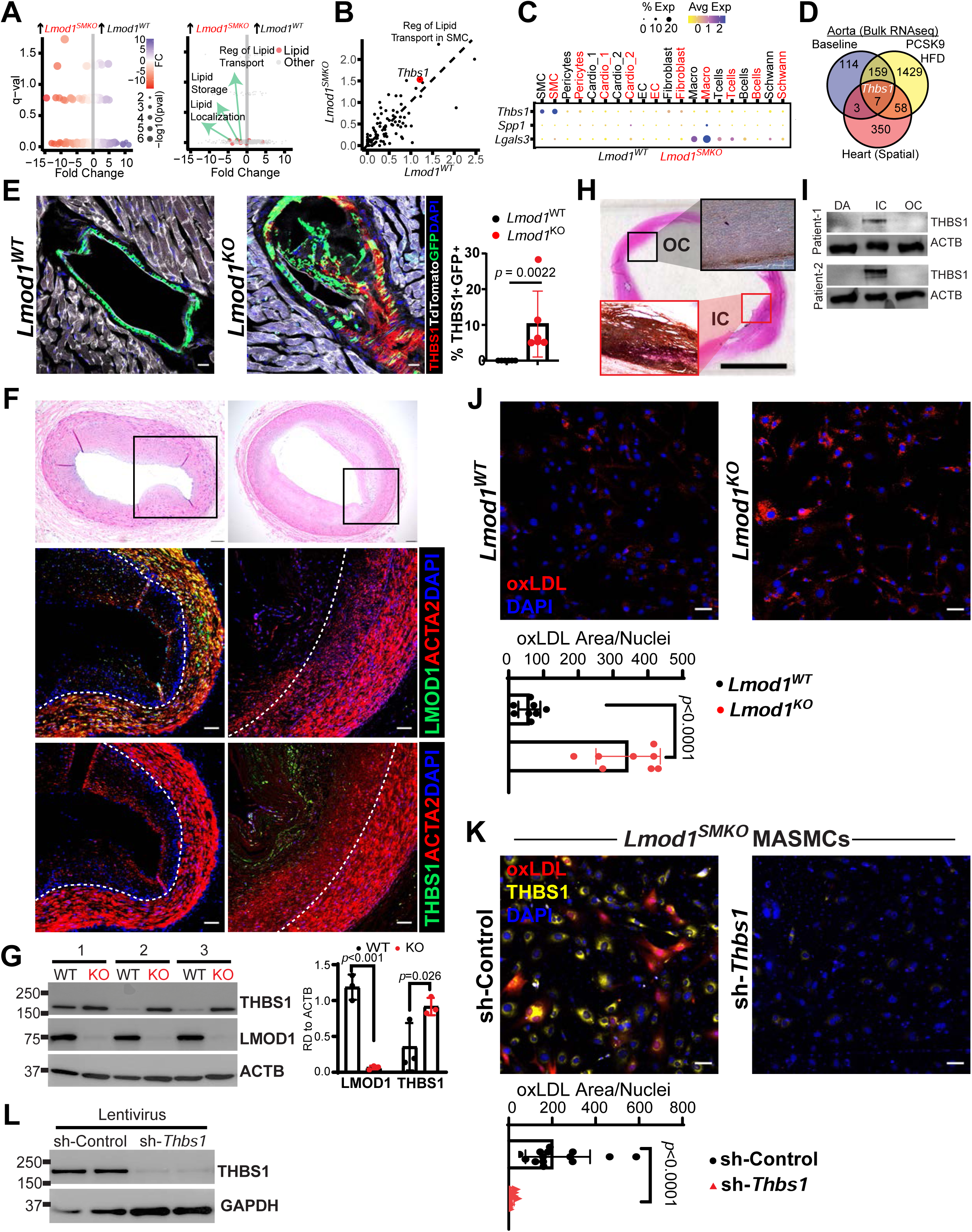
THBS1 in mouse and human atherosclerosis. **(A)** Smooth muscle cell SCPA activity shown by volcano plots with log₂ fold change (x-axis) over q-value (y-axis). First plot displays all pathways, and second plot highlights lipid-associated pathways (red) relative to non-lipid pathways (grey). **(B)** Scatterplot of genes within *GOBP Regulation of Lipid Transport* comparing *Lmod1^WT^* over *Lmod1^SMKO^* with *Thbs1* highlighted in red. **(C)** Dot plot of up-regulated genes in vascular for scRNA-seq modality. **(D)** Venn diagram illustrating shared up-regulated genes in *Lmod1^SMKO^* across two bulk RNA-seq datasets and spatial modality. (**E**) THBS1 protein colocalization with GFP+ cells in the media and plaque of *Lmod1^SMKO^* coronary artery. The tdTomato reporter was pseudo-colored white (marking mainly cardiomyocytes and endothelial cells). Scale bars, 20 μm. (**F**) Two sections of human coronary artery stained with H&E (**top**) or for the indicated proteins (**below**) magnified from H&E boxed regions. Scale bars, 50 μm. (**G**) Western blots of indicated proteins in cultured *Lmod1^WT^* or *Lmod1^SMKO^* and quantitation. (**H**) Representative histological analysis of the human aortic arch. Hematoxylin and eosin (H&E) staining shows overall tissue morphology. Oil-Red-O staining highlights lipid accumulation, particularly in the inner curvature (IC). Scale bars are 1 cm for the aorta and 400 μm for the ORO stained IC/IC. (**I**) Western blot analysis of THBS1 in segments of the descending aorta (DA), IC, and OC of the human atherosclerotic aortic arch from two patients. Quantification of THBS1 expression normalized to ACTB. Data are presented as mean ± SEM (n = 3). *p*<0.01. (**J**) OxLDL staining of primary mouse aortic SMC (MASMC) from each genotype with quantitation below. (**K**) *Lmod1^SMKO^* MASMC transduced with shRNA-control or shRNA-*Thbs1* and treated with oxLDL. Similar findings were observed in an independent experiment. (**L**) Western blot validation of the Lentiviral-mediated knockdown of THBS1 protein with shRNA-*Thbs1* in MASMC.

### The CAD phenotype is lost in mice with low level expression of a mutant LMOD1

To date, actin nucleation remains the only known and validated function of LMOD. ^34,59^ Amino acid residues 313–600 of human LMOD1, comprising the leucine rich repeat (LRR) domain and the C-terminal WH2 domain-containing extension, account for most of its nucleation activity. ^34,59^ To begin testing whether loss of this function is involved in the *Lmod1^SMKO^* CAD phenotype, we made a recombinant LMOD1 in which five basic amino acids in the LRR were substituted with five acidic residues; the five amino acid charge-reversal in mouse LMOD1 (K444D, H448D, R455D, R470D, R473D) targets highly conserved residues on the actin-binding surface of the LRR domain (**Figure 8A**). This recombinant protein (LMOD1^ND^) was purified in *E. coli* and found to be expressed equally with LMOD1^WT^ (**Figure 8A**). LMOD1^ND^ was anticipated to disrupt LMOD1 nucleation activity, as confirmed by polymerization assays where the nucleation rate of LMOD1^WT^ was over six-fold higher than that of LMOD1^ND^ (**Figure 8B**). We then generated a lentivirus carrying the same five amino acid substitutions in LMOD1^ND^ (*Lmod1^ND^* for nucleation deficient LMOD1) and transduced oxLDL-treated MASMC derived from *Lmod1^SMKO^* mice. These cells showed attenuated expression of LMOD1 and elevated THBS1, consistent with the inverse relationship of these two gene’s level of expression (see **Figure 7F, 7G**).

**Figure 8.**
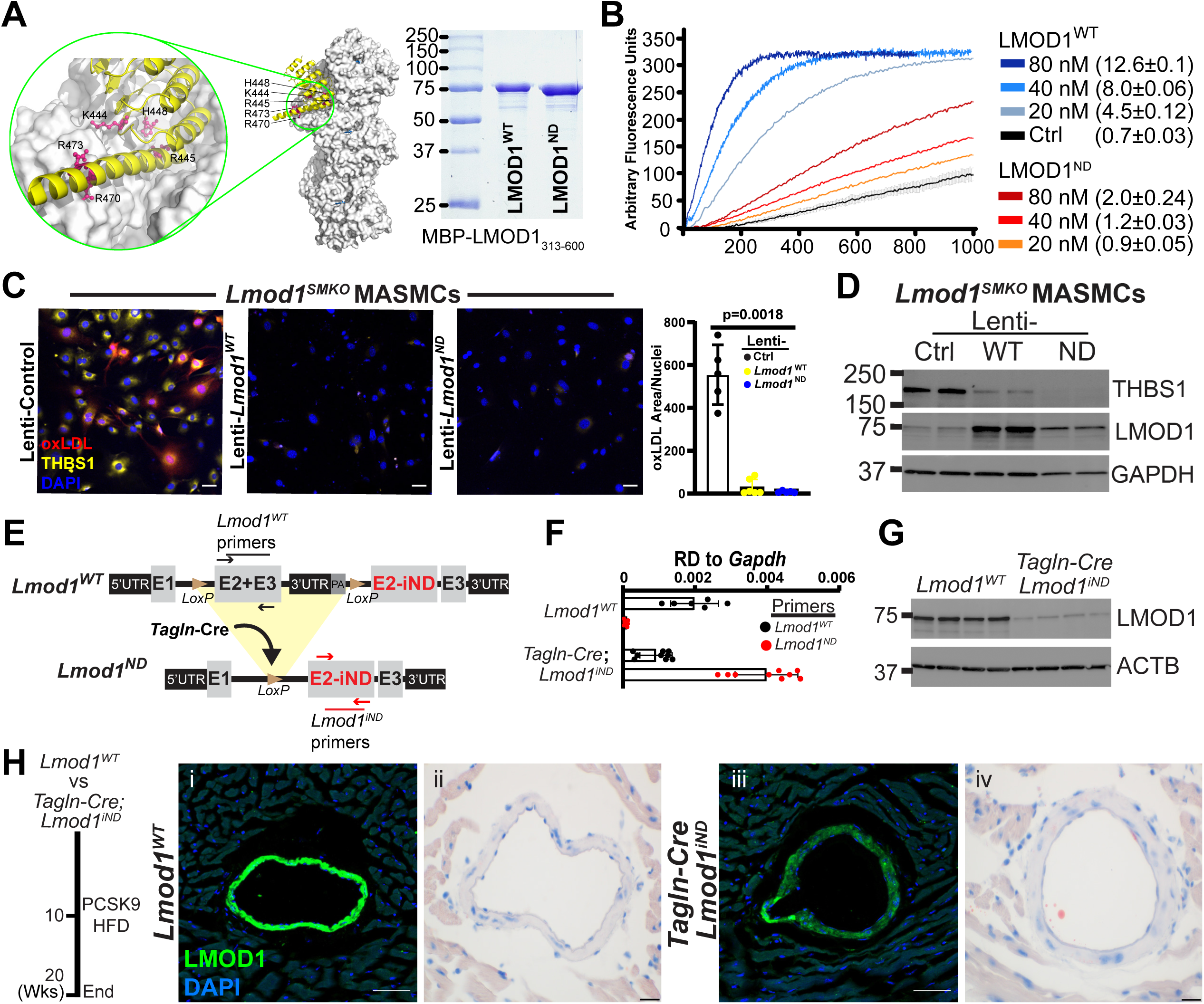
Role of LMOD1 actin nucleation domain on CAD phenotype in *Lmod1^SMKO^* mice. (**A**) LRR domain of mouse LMOD1 bound to an actin subunit at the pointed end of the actin filament, showing the mutated amino acids in LMOD1^ND^ (K444D, H448D, R455D, R470D, and R473D). Model produced based on superimposition of PDB codes 4Z79, 5WFN, and 8F8S. ^34,85^ Also shown at right is an SDS-PAGE gel of *E. coli*-expressed human LMOD1^WT^ and LMOD1^ND^ (K449D, H453D, R460D, R475D, R478D). (**B**) Time course of polymerization of 2 μM actin (6% pyrene-labelled), measured as the fluorescence increase upon incorporation of pyrene-actin into filaments. Curves represent the average of three independent experiments, color-coded by protein construct and concentration, with standard deviations (SD) shown. (**C**) *Lmod1^SMKO^* MASMC incubated with oxLDL in the presence of Lentivirus carrying each indicated construct with quantitation of oxLDL area/nuclei. (**D**) Western blot of indicated proteins from similar experiment as in panel **C**. (**E**) Strategy for making the inducible *Lmod1^iND^* mouse model. The wildtype mouse (**top**) expresses normal levels of LMOD1 from the minigene placed in first intron which is recombined out upon Cre exposure (**bottom**) enabling expression of the mutated second exon (E2-iND) which contains the five amino acid substitutions. Validation studies demonstrate the loss of *Lmod1^WT^* and induction of *Lmod1^iND^* mRNA (**F**) and protein (**G**) with *Tagln-Cre*. (**H**) CIFM of LMOD1 (**Hi, Hiii**) and ORO staining of coronary artery (**Hii, Hiv**) in *Lmod1^WT^* and *Lmod1^iND^* (induced with *Tagln-Cre*) mice treated for 10 weeks with PCSK9/HFD. n ≥ 5 mice per genotype. Scale bars, 100 μm for CIFM and 20 μm for ORO staining.

While a control lentiviral shRNA had no effect on THBS1 induction or oxLDL staining, lentivirus carrying *Lmod1^WT^* or *Lmod1^ND^* completely abolished both (**Figure 8C**). The effect of *Lmod1^ND^* was of particular interest as its expression was weaker than the LMOD1 derived from *Lmod1^WT^* lentivirus (**Figure 8D**). To explore the basis for this attenuated expression, we performed a cycloheximide study and found that the half-life of LMOD1^ND^ was considerably shorter than LMOD1^WT^ (**Figure S26A, B**). Next, we generated a Tmx-inducible *Lmod1^ND^* mouse (hereafter referred to as *Lmod1^iND^*) carrying the same five amino acid substitutions (**Figure 8E**) and validated its fidelity by Sanger sequencing (**Figure S26C**) and a qRT-PCR assay (**Figure 8F**). Next. we subjected homozygous *Lmod1^iND^* mice ± *Tagln-Cre* ^60^ to the PCSK9/HFD regimen for 10 weeks. Similar to in vitro findings above, there was a pronounced reduction in LMOD1^ND^ protein compared to LMOD1^WT^ (**Figure 8G**). Remarkably, despite such low LMOD1^ND^ protein levels, there was negligible ORO staining in coronary arteries (**Figure 8H**). To confirm these findings, we crossed the *Lmod1^iND^* mouse with *Itga8Cre-ER^T2^* and subjected these mice to the same atherogenic regimen as above. *Itga8Cre-ER^T2^*-induced *Lmod1^iND^* mice were also refractory to atherogenic-induced CAD (**Figure S26D**) and showed a similar trend in reduced LMOD1 protein expression (**Figure S26E**). Of note, mice with *Tagln-Cre* or *Itga8CreER^T2^* induced *Lmod1^iND^* showed comparable hypercholesterolemia (**Figure S26F**). Collectively, these findings imply that a low level of LMOD1 or its very low nucleation deficiency (or both) is sufficient to impede atherogenesis.

## Discussion

Efforts to define the role of the CAD risk gene *Leiomodin1* (*Lmod1*) in vascular pathobiology have been stalled by a lethal neonatal visceral myopathy of the gastrointestinal tract. ^15^ This difficulty has now been overcome with the *Itga8-CreER^T2^* mouse bred to a floxed *Lmod1* strain, generating viable VSMC-restricted *Lmod1* knockout (*Lmod1^SMKO^*) animals. *Lmod1^SMKO^* mice displayed a rapidly manifested, fully penetrant, and non-sexual dimorphic CAD phenotype with both a PCSK9/HFD or *Apoe^-/-^*/WD regimen.

In contrast, floxed *Lmod1* mice carrying the popular *Myh11-CreER^T2^* driver displayed massive intestinal distention and death, probably from sepsis, within a week of Tmx administration. It is likely other SMC Cre drivers, with activity in both vascular and visceral SMC lineages, would similarly have shown limited utility here and in other contexts where a targeted gene is of critical importance for intestinal homeostasis. Even in the absence of acute intestinal dysmotility, the inactivation of genes in both vascular and visceral SMCs could obfuscate accurate interpretation of vascular phenotypes given the importance of the gut microbiome in vascular disease.^61,62^ Thus, the potential consequences of SMC-specific gene loss on visceral smooth muscle function should be carefully considered. The smooth muscle-specific *Myh11* gene is another CAD risk allele ^5^ whose global inactivation results in a lethal visceral myopathy. ^17^ It will therefore be of interest to assess whether a VSMC-restricted knockout of *Myh11* generates a CAD-like phenotype.

Coronary atherosclerosis is rare in mice and typically requires complex genetic perturbations, ^63^ superimposed injury, ^64^ or prolonged periods of time for lesions to become manifest ^65^ (**Table S9**). Here, using standard lipid staining, coronary lineage tracing, and electron microscopy, coronary VSMCs were found to populate the base (just luminal from the IEL) of early and late atheroma; many of these VSMCs contained lipid droplets though not as much as macrophages. We also found lipid-containing VSMCs in their native medial milieu as well as the fibrous cap. These findings align well with older literature using unique ultrastructural features of VSMC foam cells, such as the presence of a basement membrane and peripherally-oriented dense bodies, ^66–68^ both of which we noted in murine coronary arteries (see **Figure S5K**). Surprisingly, despite examining both early and late stages of disease using different assays, we were unable to detect proliferating VSMCs in the *Lmod1^SMKO^* model. Moreover, not a single mitotic VSMC was ever observed in our ultrastructural analysis of dozens of vessels and thousands of individual VSMCs over the course of several years. Early human experiments showed a very low rate of VSMC proliferation in coronary plaques, although the study was conducted primarily in advanced lesions. ^69^ More recently, lineage tracing studies have demonstrated consistent clonal expansion of VSMCs in various disease processes, suggesting that a small number of medial SMCs undergo replication to generate clonal “patches.” ^70–73^ The IEMLT assay developed here determined, for the first time, that ∼46% of plaque cells were of coronary VSMC origin. The early emergence of these VSMCs in nascent coronary fatty streaks, as early as six days following the atherogenic regimen, suggests an initial migratory wave rather than proliferation of a small number of SMCs. This interpretation is supported by several examples of medial VSMCs moving through the fenestra of the IEL. Such “transfenestral migration” was elegantly demonstrated in an endothelial cell-specific knockout of the *Mef2c* transcription factor. ^74^ Further work is needed in the *Lmod1^SMKO^* model (and other models) to more accurately ascertain whether clonal expansion occurs in mouse coronary arteries and, if so, when and where VSMC proliferation occurs since migrating medial SMCs would need to be replaced.

Coronary hemorrhage, plaque rupture or endothelial cell erosion, and occlusive thrombosis all contribute to coronary symptomatology and death. Mice lacking *Lmod1* showed no evidence of these events and myocardial infarction was never observed, though studies were limited to only a few months. Therefore, this model, like most coronary atherosclerosis mouse models (**Table S9**), does not currently recapitulate the terminal events of CAD found in humans. Moreover, there was negligible atherosclerosis in cerebral vessels of the brain and no evidence of stroke or paralysis. Nevertheless, ∼35% of *Lmod1^SMKO^* mice died during the course of these studies. Other than mild neutropenia and lymphocytosis, the latter of which might be expected in this model, there were no indications of toxicity in *Lmod1^SMKO^* mice. Further, although *Lmod1^SMKO^* mice were hypotensive, it is currently unclear how this physiological change could lead to death in *Lmod1^SMKO^* mice. On the other hand, the CAD phenotype in *Lmod1^SMKO^* mice was pervasive and included the occlusion of smaller coronary vessels (<50 μm; see **Figure S10D**). We therefore favor the hypothesis that *Lmod1^SMKO^* mortality arises from ischemic cardiomyopathy and arrhythmia, though much more work is needed including extensive cardiac functional studies, coronary flow and reactivity, as well as chronic instrumentation to monitor cardiac electrical activity.

Using a variety of transcriptomic platforms, we found the matricellular protein, thrombospondin (THBS1) to be consistently elevated in VSMCs. THBS1 is a stress-responsive gene that is rapidly induced following acute vascular injury ^75^ and it has been demonstrated in human atherosclerotic vessels, albeit at low levels. ^76^ Here, we show clear evidence of THBS1 protein in both human and mouse coronary atheromatous lesions. THBS1 was also found to colocalize with GFP positive plaque cells, representing dedifferentiated VSMCs. Further, we found an inverse relationship between THBS1 and LMOD1 expression both in vivo and in vitro, a finding that was previously reported in the aortic wall of *Thbs1* null mice. ^77^ The mechanism for this opposing pattern of expression is uncertain at this time. We also observed the redistribution of THBS1 protein from a diffuse cytosolic location in *Lmod1^WT^* mice to a more membrane-bound location in SMC of *Lmod1^SMKO^* mice. It is tempting to consider that such membrane-associated localization implies an adaptive response to the massive influx of lipid into the vessel wall of *Lmod1^SMKO^* mice.

The pathophysiological role of THBS1 in atherosclerosis is unclear. An initial report found that *Apoe*/*Thbs1* double knockout mice had an increased necrotic core and reduced SMC content at the fibrous cap, suggesting a protective role for THBS1 in atherogenesis. ^78^ Conversely, a more recent study found reduced lipid staining and plaque burden in similar double-knockout mice treated with the proatherogenic adipokine, leptin. However, in the absence of leptin administration, there was no difference in lipid burden at the aortic root. ^79^ Here, we show that a shRNA targeting *Thbs1* abolished lipid accumulation in VSMCs derived from *Lmod1^SMKO^* mice. These results are the first to formally demonstrate a role for THBS1 in VSMC lipid accumulation. Of note, THBS1-binding proteins, CD36 and CD47, were also induced in vascular tissue of *Lmod1^SMKO^* mice (**Table S10**). Further work is needed including the validation of these THBS1-binding proteins in coronary SMCs and their role in THBS1-dependent lipid accumulation in VSMCs of *Lmod1^SMKO^* mice. In addition, it will be important to conduct VSMC-restricted *Lmod1*/*Thbs1* double knockout mouse experiments under atherogenic conditions and assess the CAD phenotype.

As shown here and by others, ^13^ LMOD1 protein levels are attenuated in human vascular diseases such as atherosclerosis. Further, LMOD1 is a known CAD risk allele, harboring two SNVs located in the 45 kilobase first intron. ^6,54,55^ One of these SNVs (rs34091558) was recently demonstrated to disrupt a nearly 100% conserved FOXO3 binding site, leading to reduced LMOD1 expression in cultured human coronary artery SMCs. ^58^ Interestingly, the rs34091558 SNV is located three helical turns away from a highly conserved serum response factor (SRF) binding site known as a CArG box (see **Figure 5A**). SRF-binding CArG boxes are found in most SMC regulatory regions and help direct the normal program of SMC differentiation. ^80^ In this report, we generated an 11.35 kilobase deletion in mice encompassing the SRF-CArG binding site and the adjacent orthologous SNV region. This deletion reduced baseline LMOD1 protein levels to those seen in heterozygous *Lmod1^SMKO^* mice. However, despite ∼50% reduction in the level of LMOD1 protein, heterozygous *Lmod1^SMKO^* mice failed to develop CAD. These results support that *Lmod1^SMKO^* mice do not exhibit haploinsufficiency for the CAD phenotype. It would be interesting to impose an additional risk factor for CAD, as was done recently in another mouse model of coronary atherosclerosis, ^81^ to determine whether augmented stress provokes CAD in heterozygous *Lmod1^SMKO^* mice.

How might loss of a VSMC-restricted cytoskeletal gene (*Lmod1*) trigger the rapid onset of CAD under hypercholesterolemic conditions? One possibility is a mechanism that intersects with LMOD1’s only validated function – actin nucleation. To address this possibility, we generated a mutant LMOD1 defective for actin nucleation (LMOD1^ND^). Both LMOD1^ND^ and LMOD1^WT^ fully reversed THBS1 upregulation and lipid accumulation in *Lmod1^SMKO^* VSMCs. Further, mice expressing the nucleation deficient LMOD1 mutation showed virtually no evidence of lipid insudation in the vessel wall following 10 weeks of PCSK9/HFD. This would seem to imply that loss of LMOD1’s ability to efficiently nucleate actin is sufficient to protect against CAD and that other unknown mechanisms are at play. However, we found a pronounced decrease in expression of LMOD1^ND^, both in cultured aortic SMCs and in coronary arteries of mice carrying the inducible knockin mutation. Cycloheximide studies indicate that reduced LMOD1^ND^ protein expression is due to a much shorter half-life than that seen with LMOD1^WT^. In this context, a clinical study found a similar unstable LMOD1 in a compound heterozygous patient with mutations in the vicinity of the five amino acid substitutions generated here for the *Lmod1^iND^* mouse. It is remarkable that such low-level expression of the mutant LMOD1 confers near complete protection against CAD. Relatedly, a previous report on a compound heterozygous mouse with low-level LMOD1 failed to exhibit the visceral myopathy seen in homozygous null mice. ^15^ Further, similar findings were found with LMOD2, where as little as 15% of wild-type protein was sufficient to prevent the lethal cardiac failure observed with complete LMOD2 loss. ^82^ These findings suggest that a very low level of LMOD1 protein is sufficient to maintain VSMC homeostasis. On the other hand, low-level *Lmod1^ND^* fully rescued the lipid accumulation phenotype in vitro (**Figure 8C**), suggesting an actin nucleation-independent function of LMOD1. Of course, we cannot formally rule out the rescue being a function of very low actin nucleation activity.

There are some limitations of this study that should be noted. First, studies were limited to a short time interval of only three or four months, and the underlying cause of death remains to be determined. Longer term studies will provide insight into whether lesions evolve into complex plaques, precipitating myocardial infarction and/or heart failure. Second, the small number of coronary SMCs captured in spatial and scRNA-seq modalities prohibited sub-clustering of SMCs into known dedifferentiated cell states of interest.^10,11,83^ Further, the molecular changes in cardiomyocytes and ECs were not studied here. These cells may contribute to cardiac ischemia or plaque biology (eg, endothelial-to-mesenchymal transition). Future studies should include higher resolution spatial transcriptomics and enriching for VSMCs by dissecting coronaries and/or flow-mediated enrichment of coronary VSMCs to enable rigorous cell-cell communication analyses. In addition, there is need for comparative transcriptomic studies at different stages of human and mouse coronary atherosclerosis. A related limitation is our incomplete understanding of coronary SMC-derived plaque cells. Although THBS1 was co-localized with GFP cells of the plaque, there was a surprisingly low number of SPP1, LGALS3, and CD68 marked GFP cells. Further, the question of coronary SMC clonality remains unresolved, particularly with respect to the timing and location of SMC expansion in these vessels.

Finally, given that atherogenesis is initiated by EC inflammation, how LMOD1—a cytoskeletal regulator of actin nucleation—promotes massive lipid infiltration into the intima remains unknown.

Nevertheless, the findings here represent the first SMC-specific risk allele for CAD to be studied in adult mice under an atherogenic regimen. With the exception of *Nos3*, ^84^ *Lmod1* is the only non-lipid related CAD risk gene whose genetic knockout yields a CAD phenotype in mice. The rapid manifestation of coronary atherosclerosis in *Lmod1^SMKO^* mice, while not modeling the more indolent nature of the human disease, offers an opportunity to rapidly test therapeutics that mitigate disease progression or promote plaque stability.

## Acknowledgments

Studies were supported by grants from the National Institutes of Health (HL147476 and HL173111 to JMM; HL139794 to XL; HL164792 to GC; HL136865 to VN; DK132888 to RIVP; GM152412 to RD) and the Florida Department of Health (grant number 22K07 to RIVP). We gratefully acknowledge Yueh-Chiang Hu (University of Cincinnati Mouse Core) for generating the floxed *Lmod1* mouse; the University of Rochester Genomics Research Center; and the Augusta University Electron Microscopy and Histology Core. Mass spectrometry imaging was performed in the University of Texas at Austin Mass Spectrometry Imaging Facility supported by Cancer Prevention and Research Institute of Texas awards RP190617/RP240559.

## Nonstandard abbreviations and acronyms

AAV: adeno-associated virus
Apoe: apolipoprotein E
ARRIVE: animal research: reporting of in vivo experiments
BSA: bovine serum albumin
CA: coronary arterial
CIFM: confocal immunofluorescence microscopy
CM: cardiomyocyte
Cre: causes recombination
CRISPR: clustered regularly interspaced short palindromic repeats
DEGs: differentially expressed genes
EC: endothelial cell
Edu: 5-ethynyl-2′-deoxyuridine
ER^T2^: estrogen receptor ligand-binding domainTamoxifen, 2nd generation
FFPE: formalin-fixed, paraffin-embedded
GEM: gel beads-in-emulsion
GOBP: Gene Ontology Biological Process
GFP: green fluorescence protein
GWAS: genome wide association studies
HFD: high fat diet
IEL: internal elastic lamina
IEMLT: immunogold electron microscopy lineage tracing
Itga8: integrin alpha 8
LMOD1: leiomodin1
LoxP: locus of X-over P1
MASMC: mouse aortic SMC
mTmG: membrane tomato, membrane green fluorescent protein
Myh11: myosin heavy chain 11
mTmG: membrane tomato, membrane green fluorescence protein
ORO: oil-red-o
ox-LDL: oxidized low-density lipoprotein
PCSK9: proprotein convertase subtilisin/kexin type 9
PFA: paraformaldehyde
scATAC-seq: single cell assay for transposase-accessible chromatin using sequencing
SCPA: single cell pathway analysis
scRNA-seq: single cell RNA sequencing
SEM: standard error of mean
SRF: serum response factor
SMI: spatial mass spectrometry imaging
SMKO: smooth muscle knockout
SNV: single nucleotide variant
TEM: transmission electron microscopy
Thbs1: thrombospondin
Tmx: tamoxifen
TUNEL: terminal deoxynucleotidyl transferase–mediated dUTP nick end labeling
UMAP: uniform manifold approximation and projection
VSMC: vascular smooth muscle cell(s)
WD: western diet
WT: wild type

## Supplemental Figure Legends

**Figure S1.**
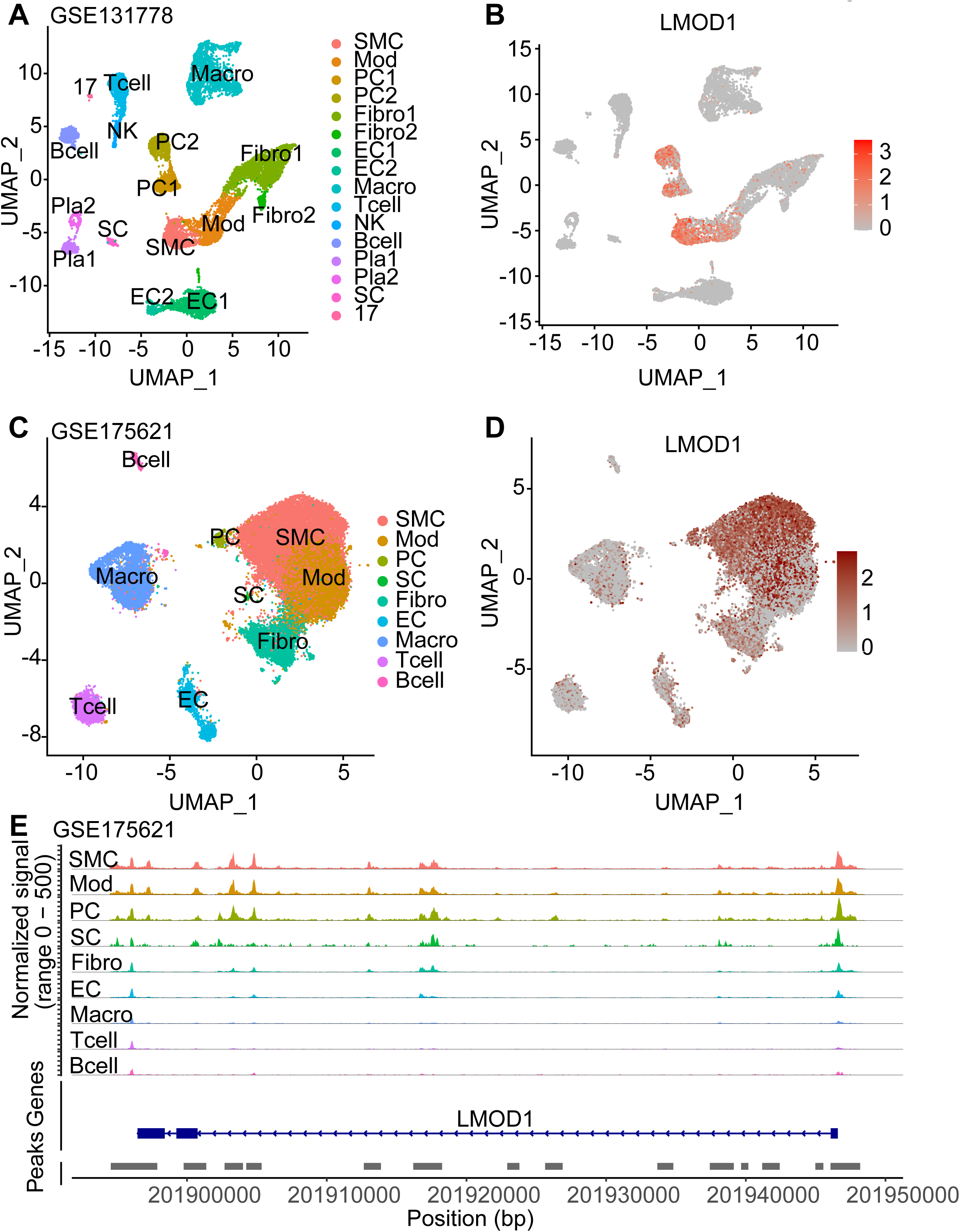
*LMOD1* omics analyses in human coronary artery atherosclerosis. UMAP (A) and attending feature plot of *LMOD1* (B) derived from indicated scRNA-seq study ^1^ of human coronary atherosclerosis. UMAP (C) and attending feature plot of *LMOD1* (D) derived from indicated scATAC-seq study ^2^ of human coronary atherosclerosis. (E) Open chromatin occupancy map across the *LMOD1* locus in indicated cell types.

**Figure S2.**
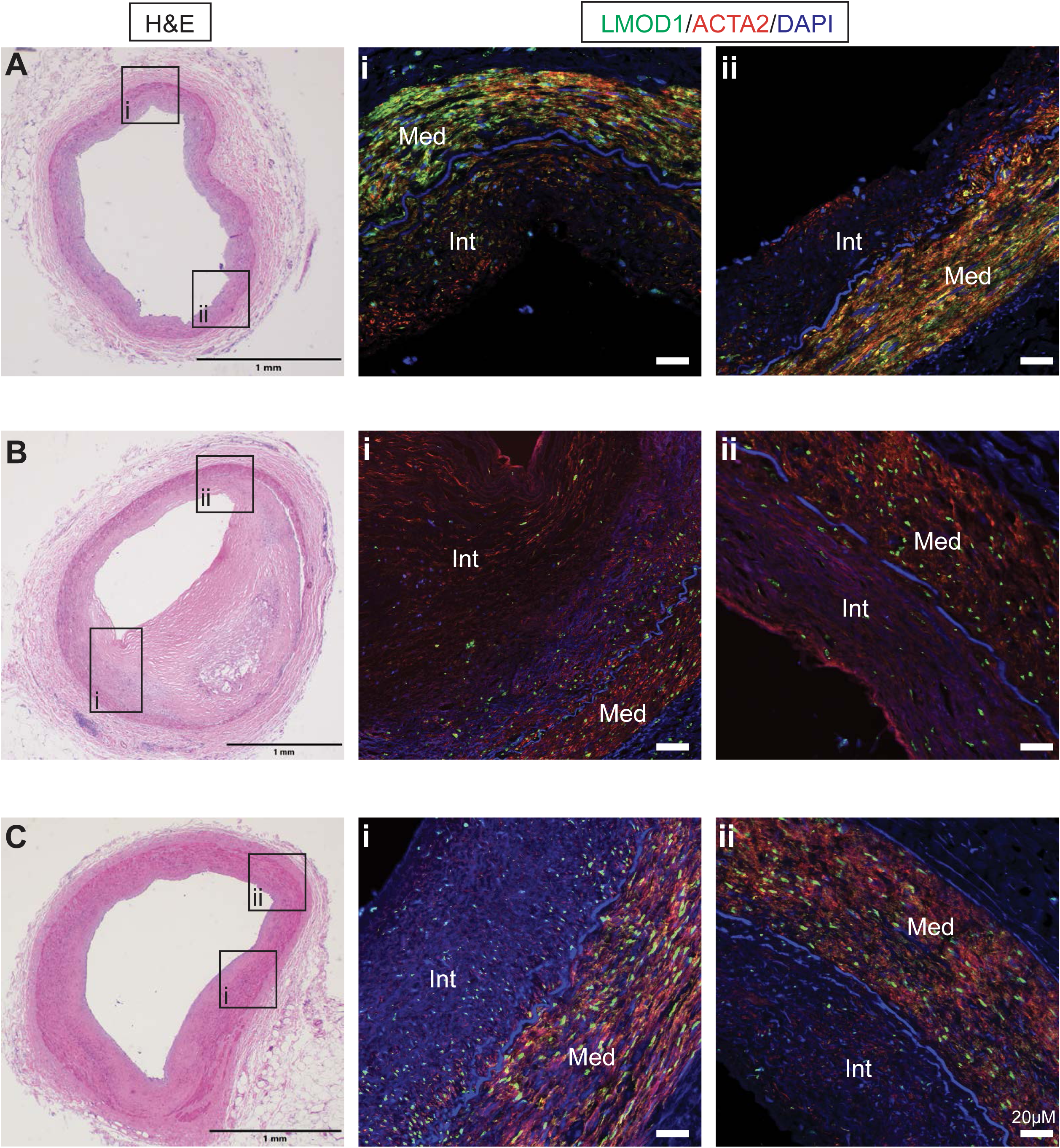
LMOD1 protein expression in human coronary atherosclerosis. Hematoxylin and eosin (H&E) staining of human coronary arteries of varying severity from male (**A, B**) and female (**C**) subjects with adjacent high magnification confocal immunofluorescence microscopy (CIFM) images of LMOD1 and ACTA2 from the boxed regions labeled i and ii in each H&E panel. Scale bars are 1 mm in the H&E images and 20 μm in the CIFM images. Int, intima; Med, media.

**Figure S3.**
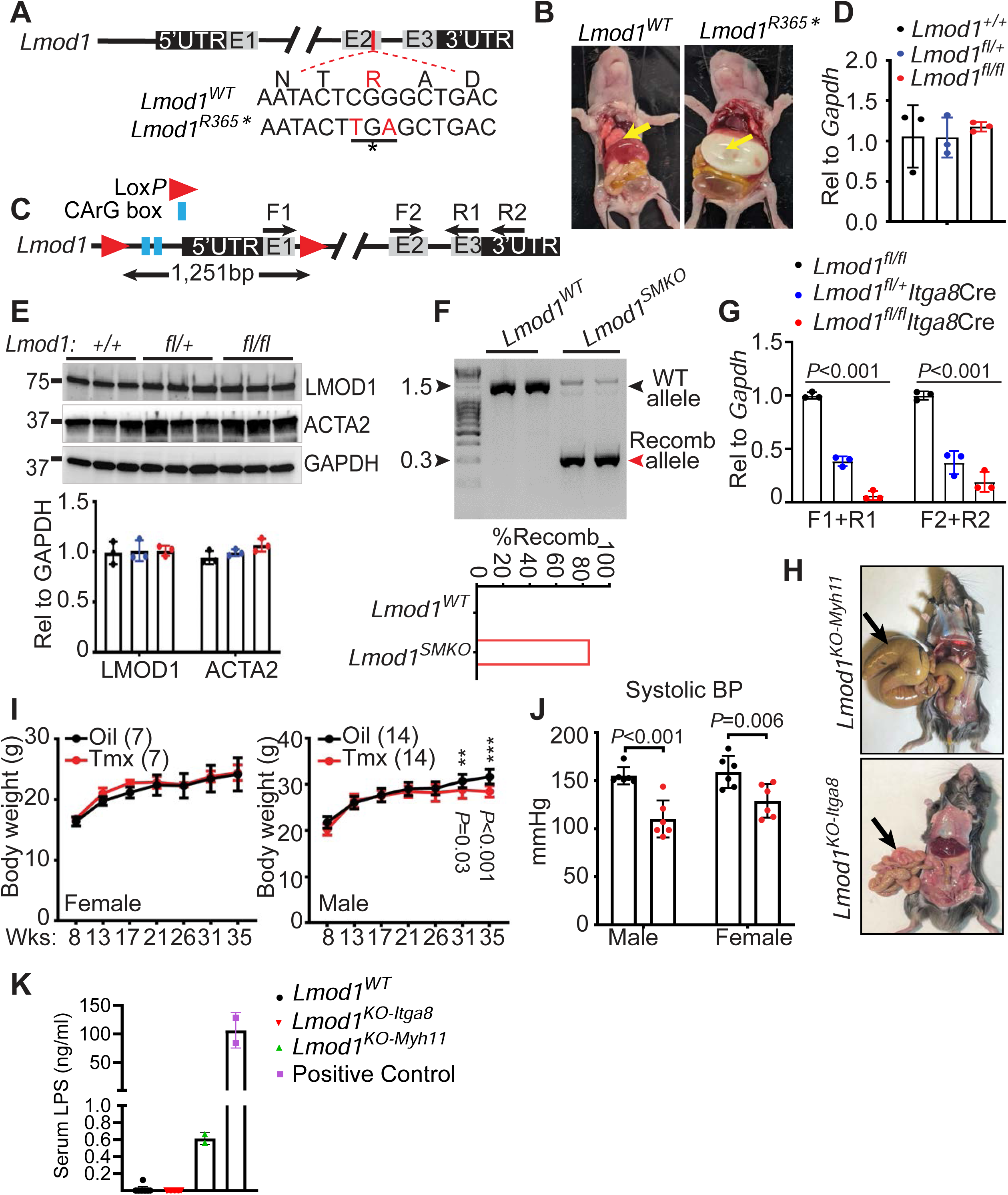
Baseline LMOD1 knockout phenotyping. (**A**) Nucleotide substitutions (red) in exon 2 of *Lmod1* generate the orthologous R365* premature stop codon previously reported in a human patient. ^3^ (**B**) Gastroparesis (yellow arrow) and megabladder (below) in *Lmod1^R365*^* mouse phenocopies the same visceral myopathy described in a human. ^3^ (**C**) Design of floxed *Lmod1* mouse. Quantitative *Lmod1* mRNA (**D**) and LMOD1 protein (**E**) expression in mice with indicated genotypes (n=3 mice). (**F**) Percent recombination of floxed *Lmod1* locus with *Itga8-CreER^T2^* driver (*Lmod1^SMKO^)* using conventional PCR (**top**) and quantitative PCR (**bottom**) of genomic DNA derived from mouse aorta. (**G**) Quantitative RT-PCR of *Lmod1* mRNA in aorta with primers depicted in panel **C** (n=3 mice). The reduced signal with primers F2+R2 signify the absence of an internal promoter yielding a mature *Lmod1* mRNA, thus validating this conditional knockout as a true null allele. ^4^ (**H**) Intestinal myopathy (arrow) in conditional knockout of *Lmod1* using *Myh11-CreER^T2^* (labeled as *Lmod1^KO-Myh11^*), but not in mice where the *Itga8-CreER^T2^* driver was used (labeled *Lmod1^KO-Itga8^*). (**I**) Body weights of homozygous floxed *Lmod1*/*Itga8Cre-ER^T2^* mice treated with oil (black) or tamoxifen (Tmx; red) beginning at eight weeks of age. The number of mice in each arm of the experiment is indicated in parentheses. (**J**) Systolic blood pressure, assessed by tail cuff method in oil (black) versus Tmx (red) homozygous floxed *Lmod1*/*Itga8Cre-ER^T2^* mice (n=6 mice). (**K**) Serum lipopolysaccharide (LPS) levels in conditional *Lmod1* knockout mice using the indicated Cre drivers.

**Figure S4.**
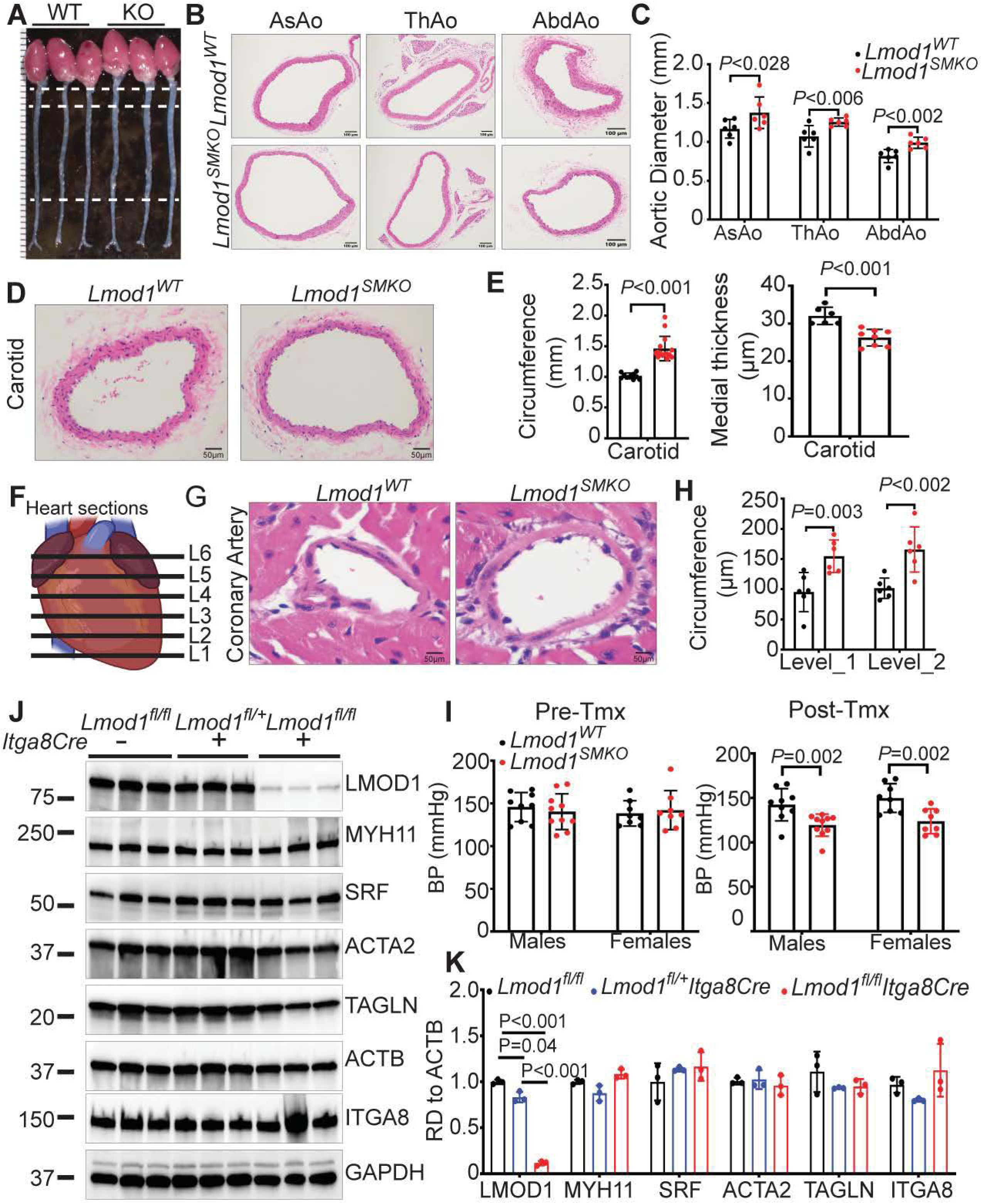
Baseline vascular phenotype in *Lmod1^SMKO^* mice. (A) Gross images of wildtype and *Lmod1^SMKO^* heart/aorta showing obvious thickened aortic wall in KO and areas (white dotted lines) sampled for thoracic and abdominal aortic sections. (B) H&E cross-sections of ascending aorta (AsAo), thoracic aorta (ThAo), and abdominal aorta (AbdAo) from *Lmod1^WT^* and *Lmod1^SMKO^* mice (n=3 mice per region). (C) Quantitation of aortic diameter from gross images of *Lmod1^WT^* and *Lmod1^SMKO^* aortae (n=6 mice per region). (D) H&E images of carotid artery and (E) quantitative measures of circumference and medial thickness from *Lmod1^WT^* and *Lmod1^SMKO^* carotids (n ≥ 6 mice). (F) Schematic of sampling (in 1mm levels) from base to apex of heart. (G) H&E images of coronary artery from *Lmod1^WT^* and *Lmod1^SMKO^* mice and (H) quantitative measures of circumference at Levels 1 and 2 (n=6 mice). (I) Baseline blood pressure before and 10 days after Tamoxifen (Tmx) administration in male and female WT and *Lmod1^SMKO^* mice (n=8 mice). (J) Western blots of indicated proteins of each genotype and (J) quantitative data (n=3 mice per genotype).

**Figure S5.**
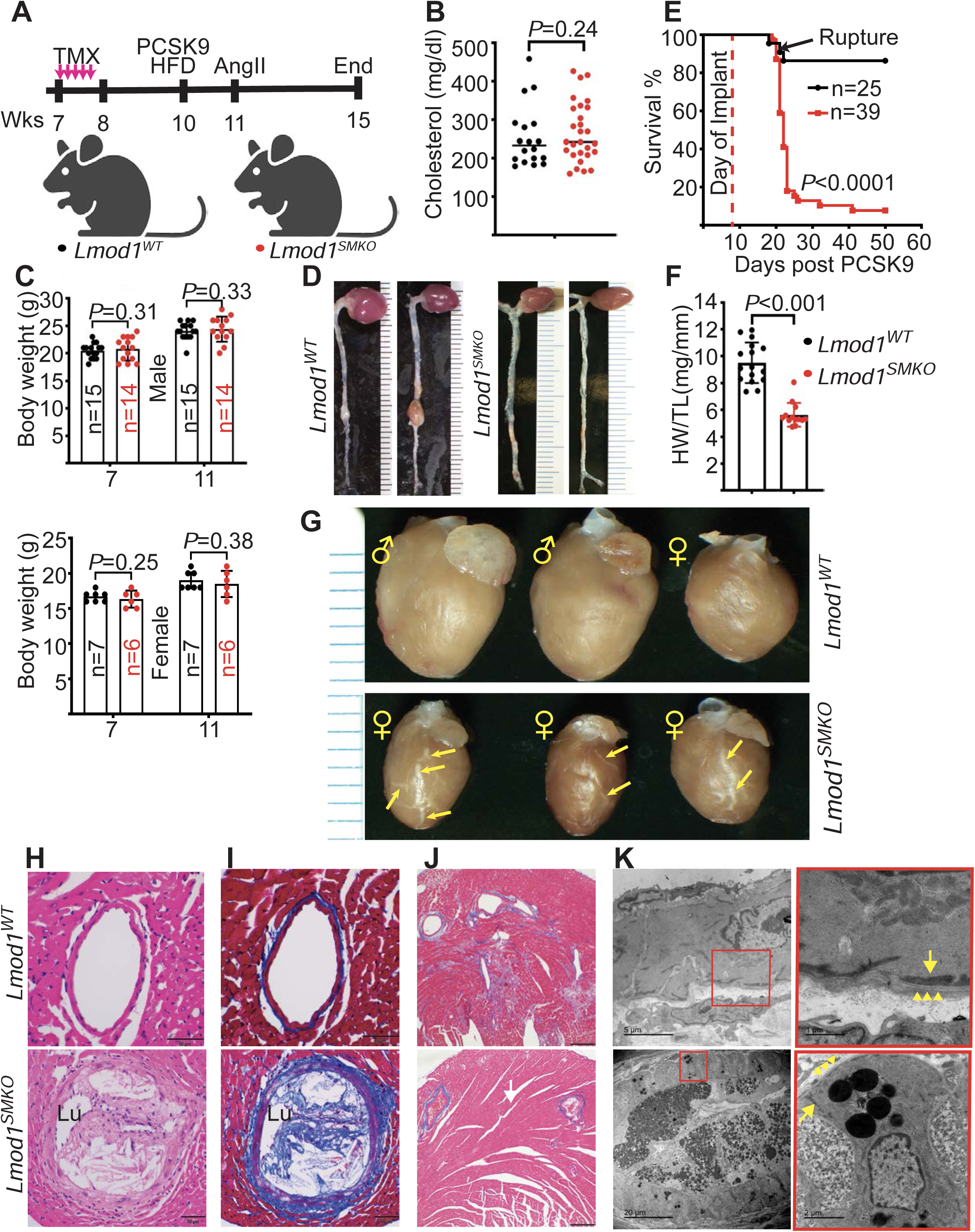
Cardiovascular changes in *Lmod1^SMKO^* mice subjected to an athero-aneurysm regimen. (**A**) Experimental design of study. Note, *Lmod1^WT^* mice here were of genotype *Lmod1^fl/fl^* and received the same schedule of PCSK9/HFD and angiotensin II as *Lmod1^SMKO^* mice. (**B**) Total cholesterol levels two weeks after the start of PCSK9/HFD in *Lmod1^WT^* (black dots, n=18) and *Lmod1^SMKO^* (red dots, n=28) mice. (**C**) Body weights in male (**top**) and female (**bottom**) mice of same genotype as above. (**D**) AAA in *Lmod1^WT^* but not *Lmod1^SMKO^* mice. (**E**) Survival curve of *Lmod1^WT^* versus *Lmod1^SMKO^* mice. (**F**) Heart weight (HW) to tibial length (TL) in *Lmod1^WT^* (n=16) and *Lmod1^SMKO^* (n=15) mice at the termination of study. (**G**) Gross heart images showing tofu-like coronary arteries (yellow arrows) in *Lmod1^SMKO^* mice. Hematoxylin (**H**) and Masson trichrome (**I**) stained coronary arteries; note small lumen (Lu) in *Lmod1^SMKO^* mice. (**J**) Masson trichrome stained heart tissue shows interstitial fibrosis in *Lmod1^WT^* but not *Lmod1^SMKO^* mice (n=3 mice). (**K**) Transmission electron microscopy of coronary arteries. Both *Lmod1^WT^* and lipid droplet-containing *Lmod1^SMKO^* coronary arteries show characteristic peripheral dense bodies (arrows) and basement membrane (arrowheads). Note the nearly 100% occluded *Lmod1^SMKO^* coronary artery with lipid-laden foam cells containing lipid droplets of variable color within residential SMC of the medial wall (magnified red boxed region at bottom).

**Figure S6.**
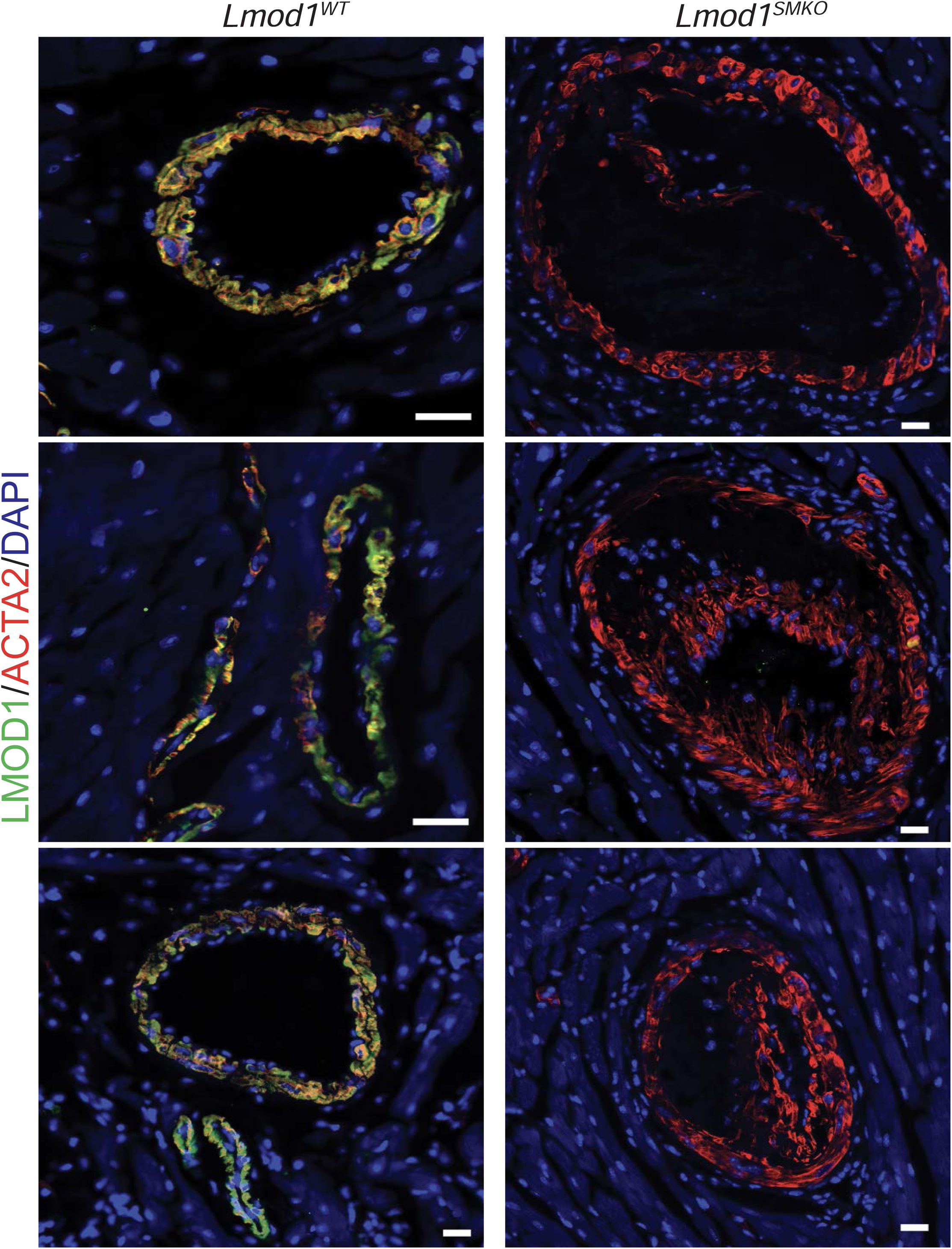
CIFM of LMOD1 and ACTA2 in coronary arteries. Images (from 74) of coronary arteries of *Lmod1^WT^* (n=8 mice total) and *Lmod1^SMKO^* (n=5 mice total) mice. Note the absence of LMOD1 staining and variable ACTA2 positive fibrous caps in the lesions of *Lmod1^SMKO^* mice. No such lesions were ever seen in *Lmod1^WT^* mice. Images from mice given the PCSK9/HFD regimen for 52 days. Scale bars, 20 μm.

**Figure S7.**
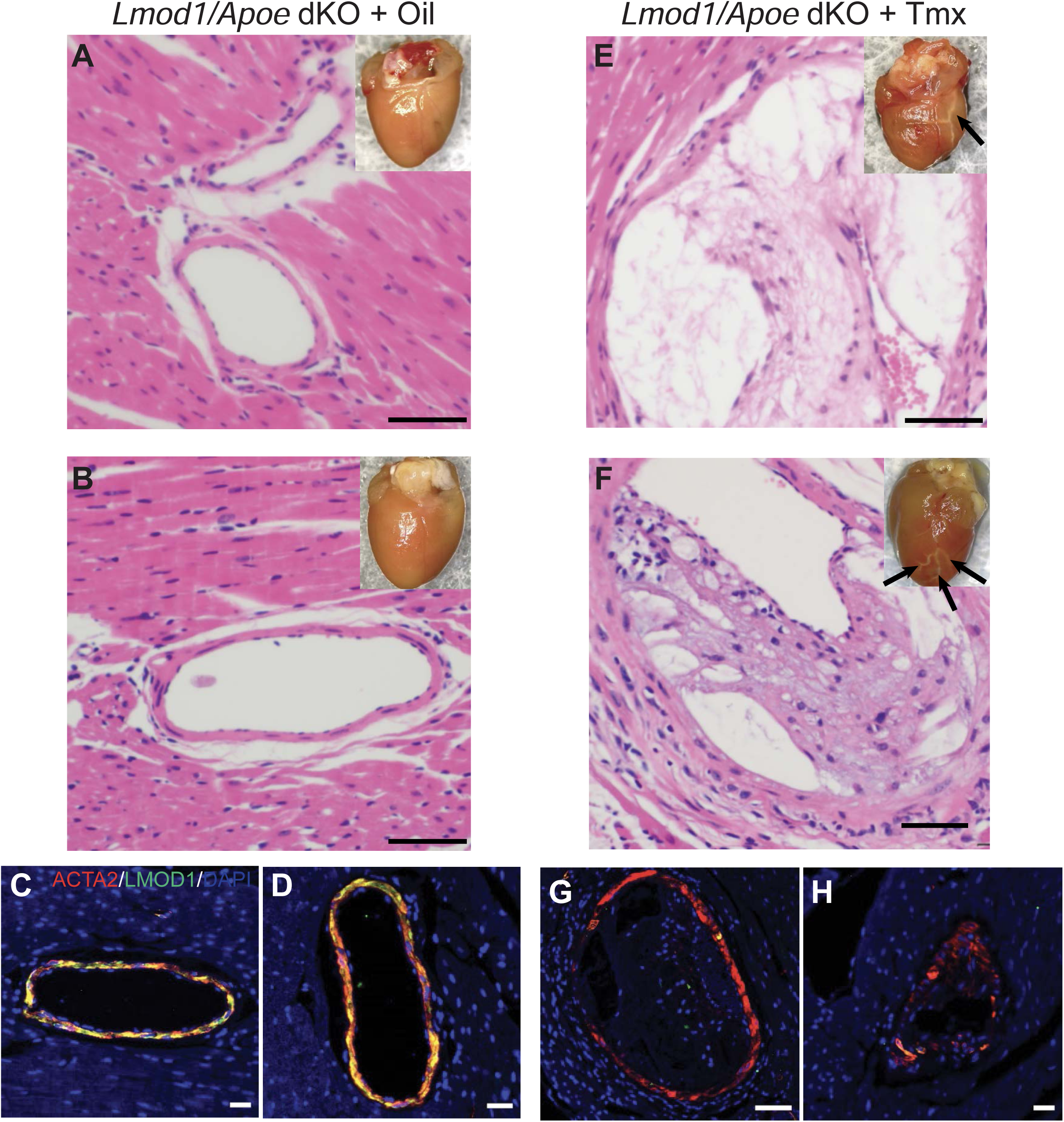
CAD phenotype in *Lmod1*/*Apoe* double knockout (dKO) mice. Pairs of control (A-D) and *Lmod1*/*Apoe* dKO (E-H) mice were subjected to a Western Diet for 11 weeks and hearts processed for imaging and staining with H&E (A, B, E, F) or CIFM imaging (C, D, G, H) for ACTA2 and LMOD1. Note tofu-like appearance of coronaries (black arrows in insets) of dKO hearts. These results were found in at least two additional pairs of mice. Scale bars are 50 μm for H&E and 20 μm for CIFM.

**Figure S8.**
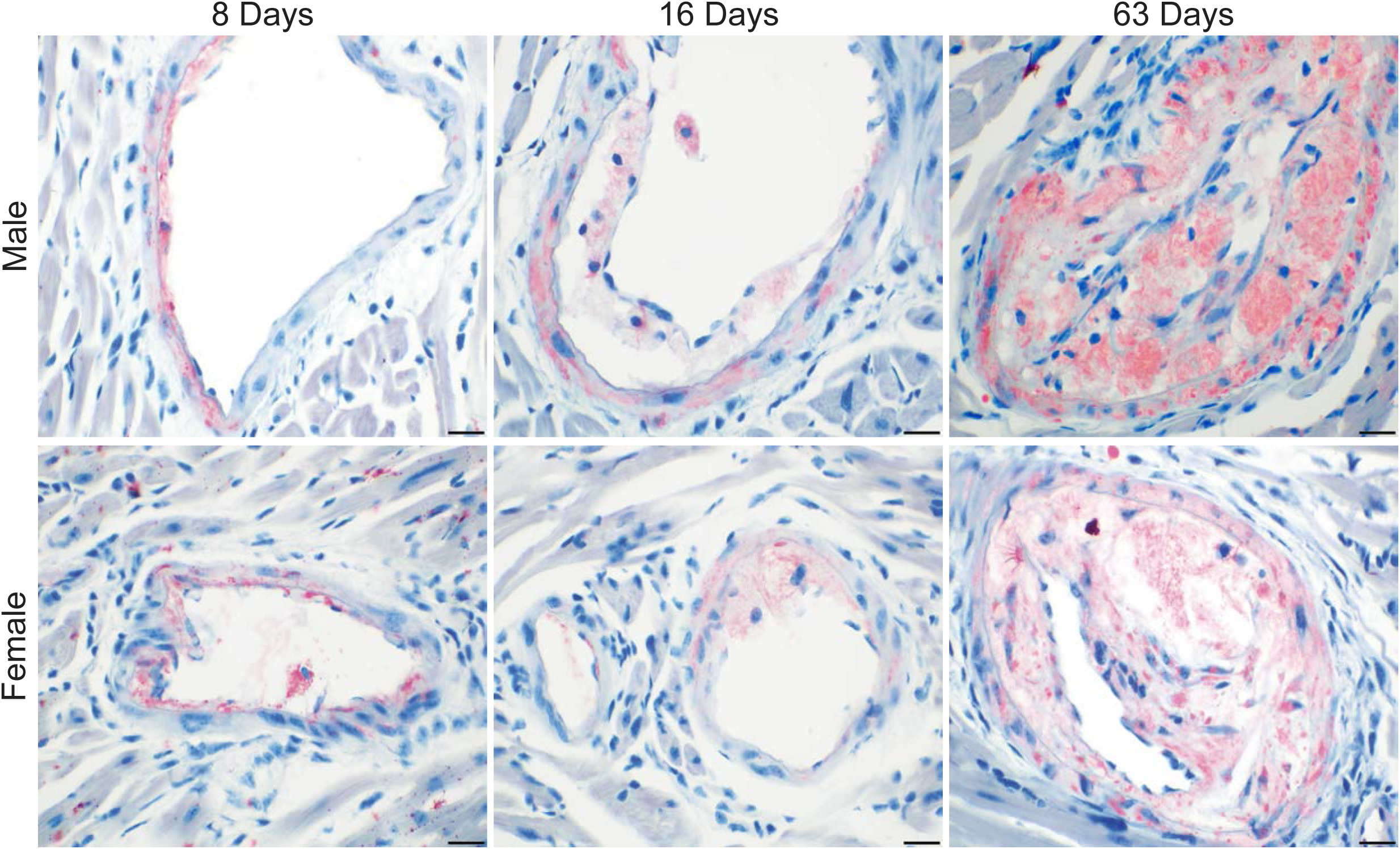
ORO-positive coronary lesions over time in *Lmod1^SMKO^* mice. Both male (top row) and female (bottom row) coronary arteries stained with ORO at the indicated times following PCSK9/HFD treatment. Scale bars are 20 μm.

**Figure S9.**
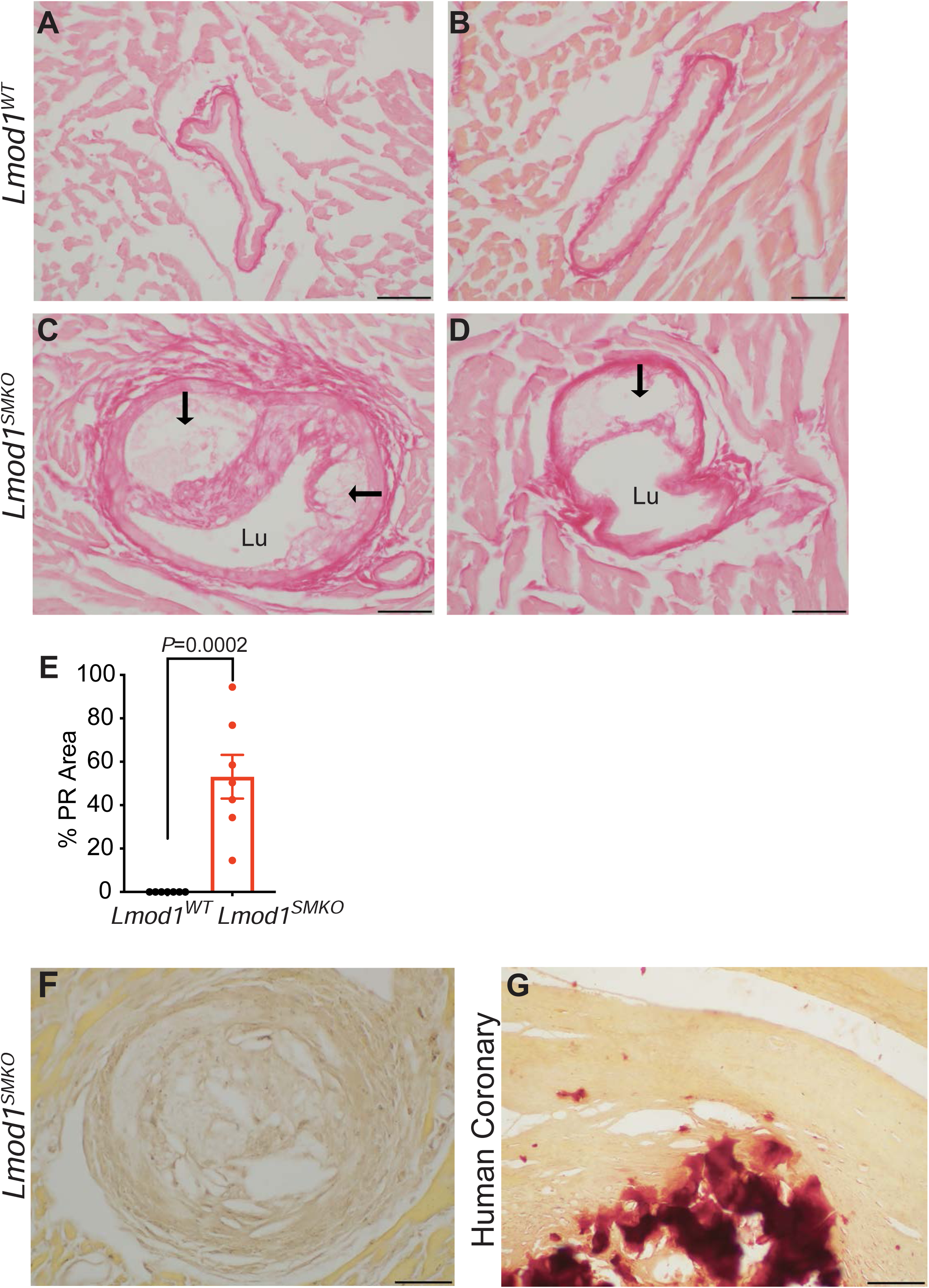
Picrosirius red and Alizarin red staining of coronary arteries. Picrosirius red stained coronaries from *Lmod1^WT^* (**A, B**) and *Lmod1^SMKO^* (**C, D**) mice. Note lumen (Lu) and necrotic core of atheroma (arrows) in *Lmod1^SMKO^* coronary arteries. (**E**) Percent plaque area stained for picrosirius red (PR) in *Lmod1^WT^* and *Lmod1^SMKO^* coronary arteries. Data represent measures of coronary arteries from seven independent mice per genotype. Alizarin red staining was not observed in plaques of *Lmod1^SMKO^* mice (**F**), but was readily observed in a calcified human coronary artery (**G**). All scale bars are 50 μm.

**Figure S10.**
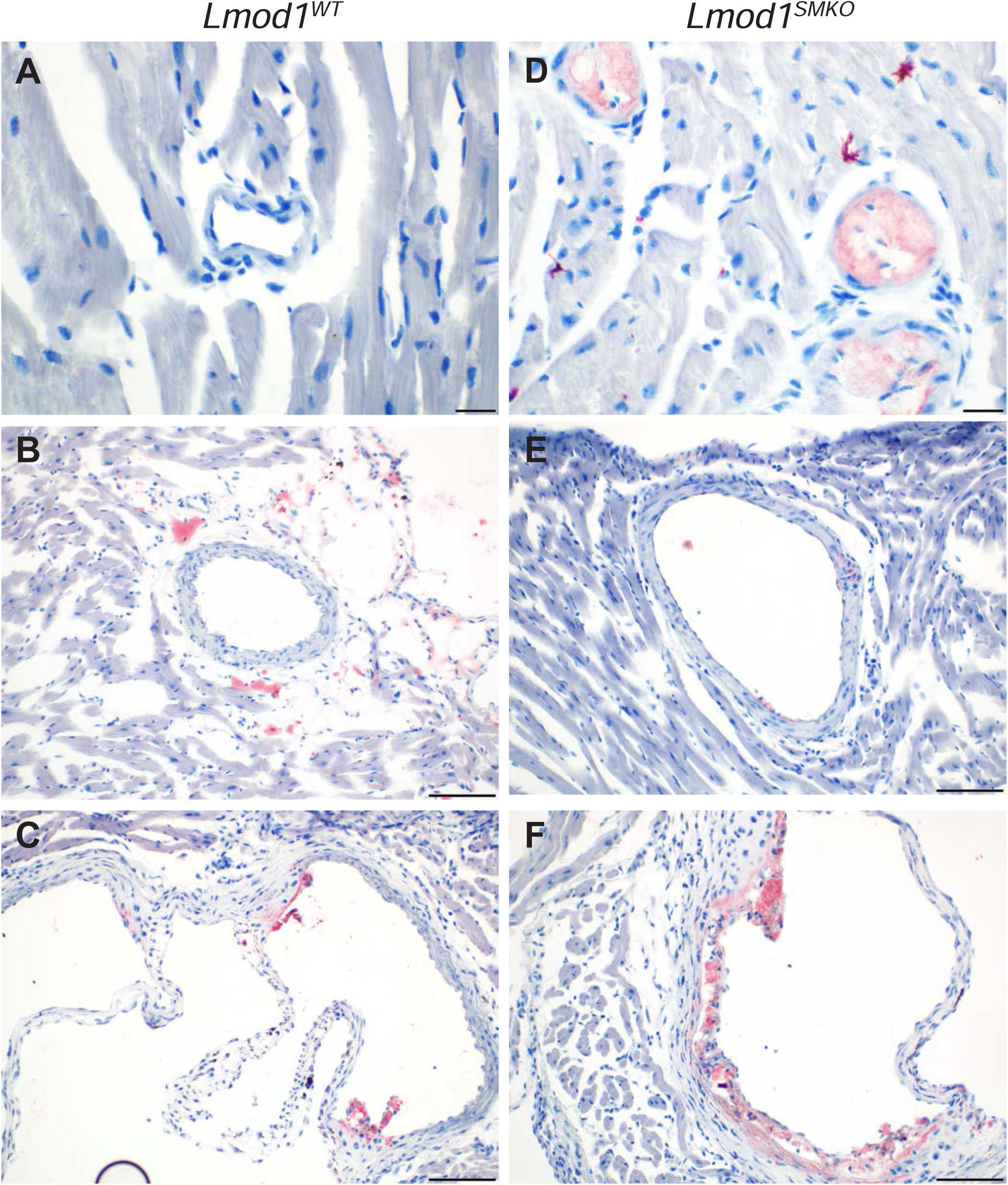
ORO-staining of aortic root and large versus small coronary arteries. *Lmod1^WT^* (A-C) and *Lmod1^SMKO^* (D-F) microvascular coronaries (A, D), branches of large coronary arteries near base of heart (B, E), and aortic root (C, F) stained for ORO. Each panel is representative of similarly-stained vessels (>50) in at least 10 mice of each genotype. Scale bars are 20 μm for panels A and D and 100 μm for panels B, C, E, and F.

**Figure S11.**
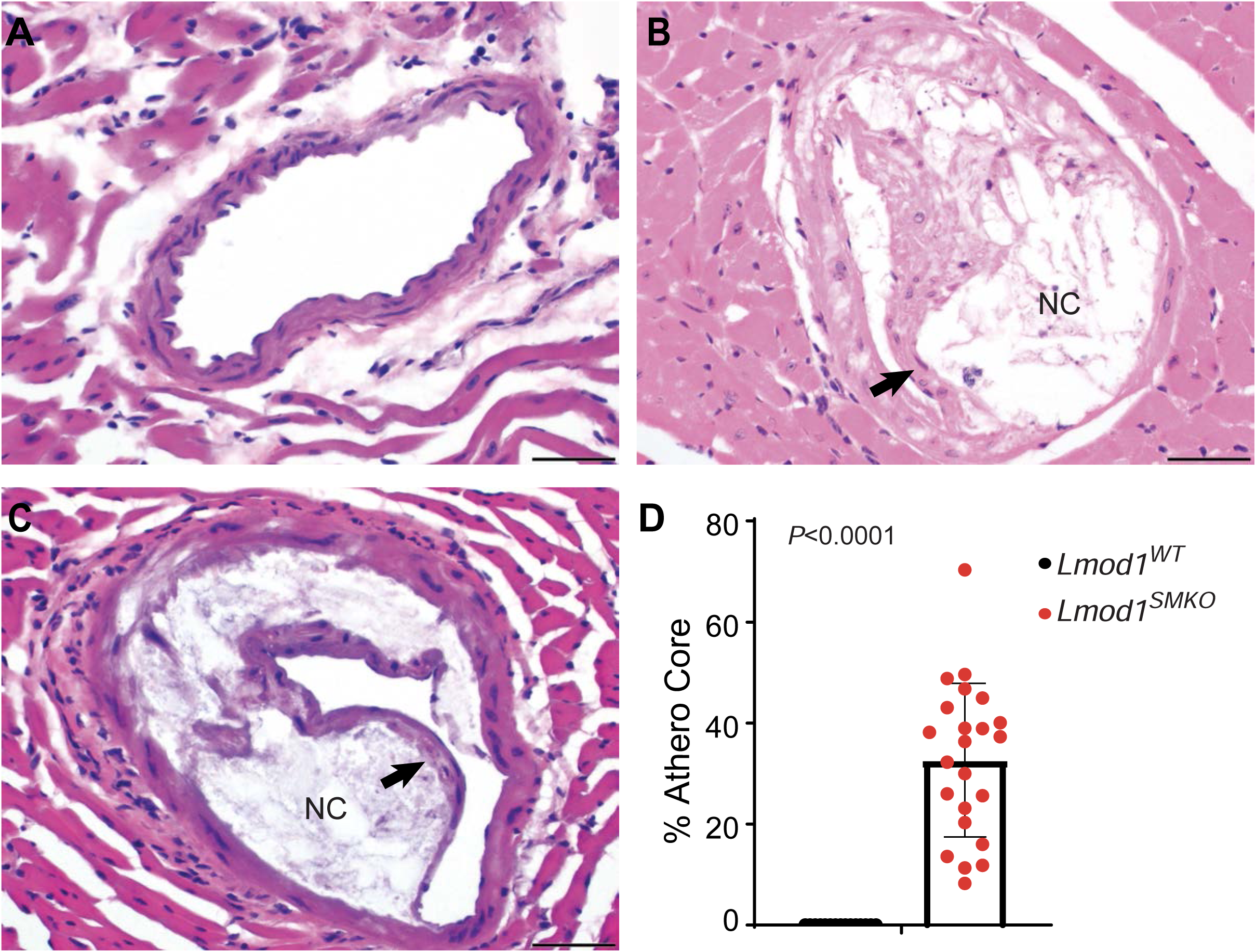
Percent necrotic core in *Lmod1^SMKO^* mice. H&E images of coronary arteries of *Lmod1^WT^* (**A**) and *Lmod1^SMKO^* male (**B**) and female (**C**) mice. (**D**) Percent of plaque occupied by necrotic core (NC) as defined by the mostly acellular region located below the fibrous cap (arrows). Data represents 15 lesions from six *Lmod1^WT^* mice and 23 lesions from 10 *Lmod1^SMKO^* mice. Scale bars are 50 μm.

**Figure S12.**
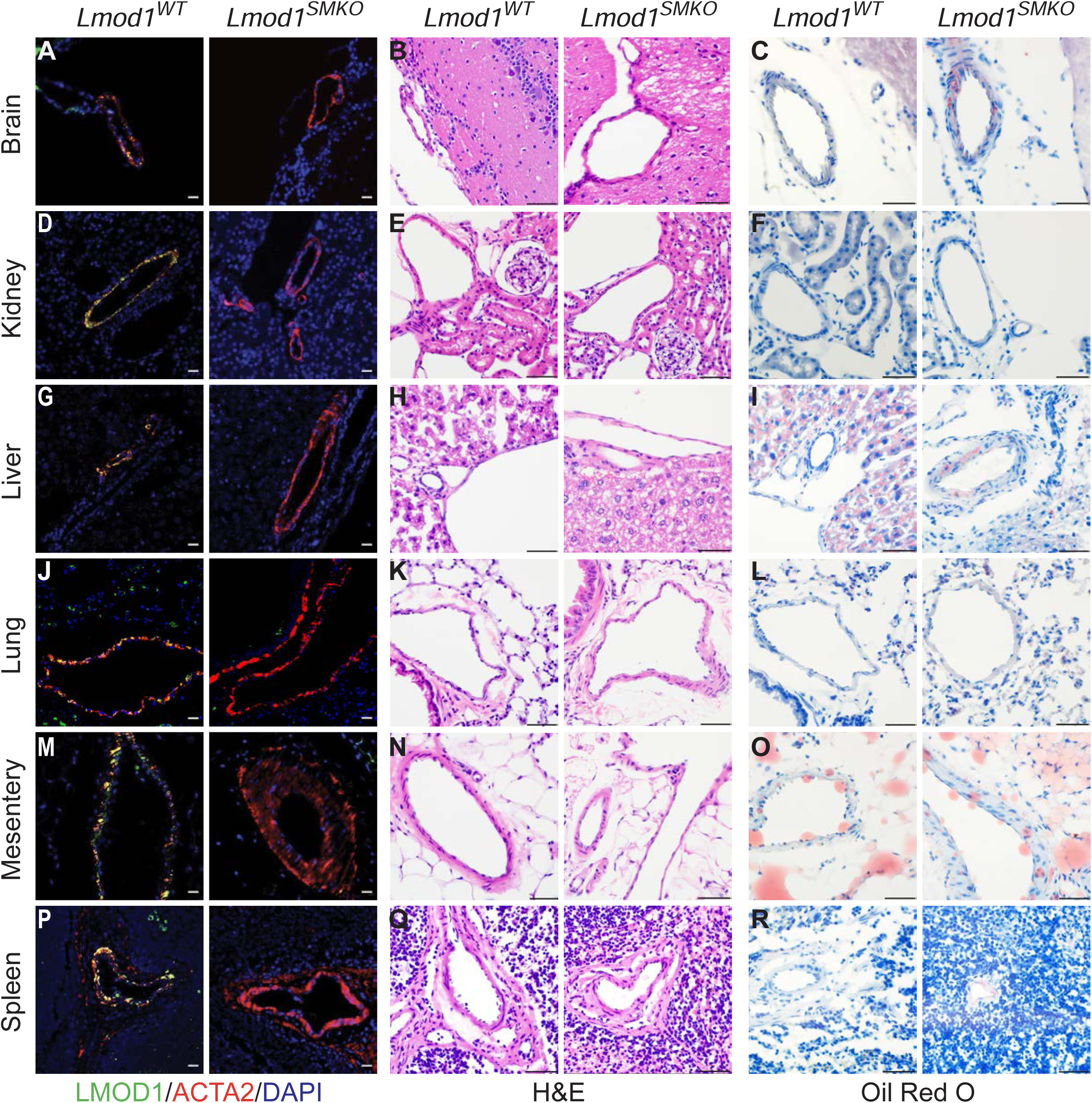
Minimal atherosclerosis in other organs of *Lmod1^SMKO^* mice. *Lmod1^WT^* and *Lmod1^SMKO^* vessels of Brain (**A-C**), Kidney (**D-F**), Liver (**G-I**), Lung (**J-L**), Mesentery (**M-O**), and Spleen (**P-R**) stained by CIFM for LMOD1 and ACTA2 (**A, D, G, J, M, P**), or H&E (**B, E, H, K, N, Q**), and ORO (**C, F, I, L, O, R**). Note that only brain (n=7 mice), liver (n=3 mice), and spleen (n=3 mice) showed occasional, mild ORO staining; most vessels were negative in these organ beds. No such mild lesions were ever seen in kidney (n=4 mice), lung (n=3 mice), or mesentery (n=3 mice) and none of the vessels from *Lmod1^WT^* mice (n=5 for brain and kidney; n=3 for liver, lung, mesentery, and spleen) showed evidence of atheromatous disease. Scale bars are 20 μm for CIFM images of LMOD1/ACTA2 and 50 μm for all other images.

**Figure S13.**
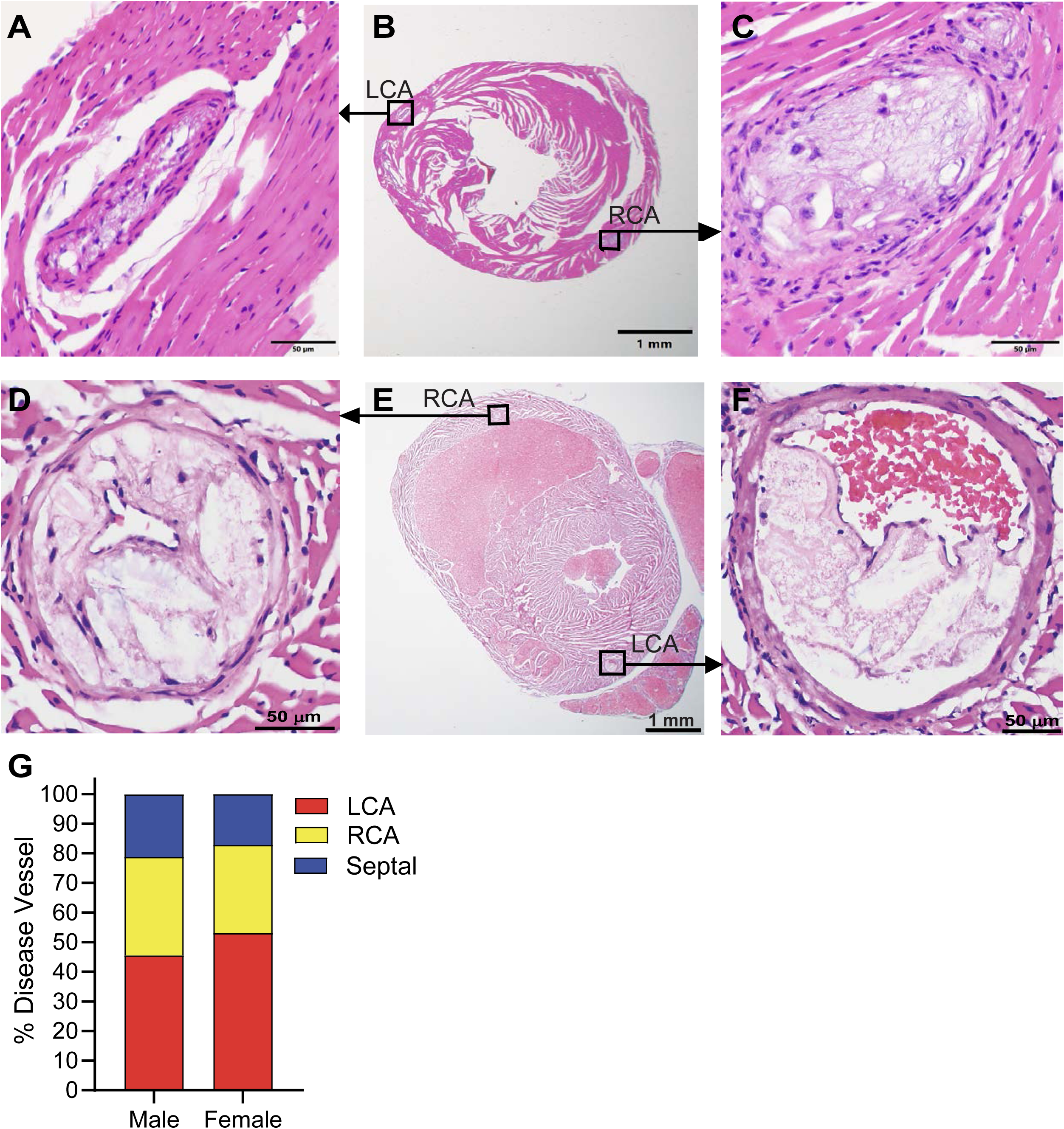
Distribution of lesions between different anatomical coronary arteries in male and female *Lmod1^SMKO^* mice. Indicated left coronary artery (LCA) and right coronary artery (RCA) of female (**A-C**) and male (**D-F**) *Lmod1^SMKO^* mouse hearts, nine weeks following PCSK9/HFD. (**G**) Distribution of lesions in each indicated coronary artery between males (92 sections from 22 mice) and females (64 sections from 12 mice).

**Figure S14.**
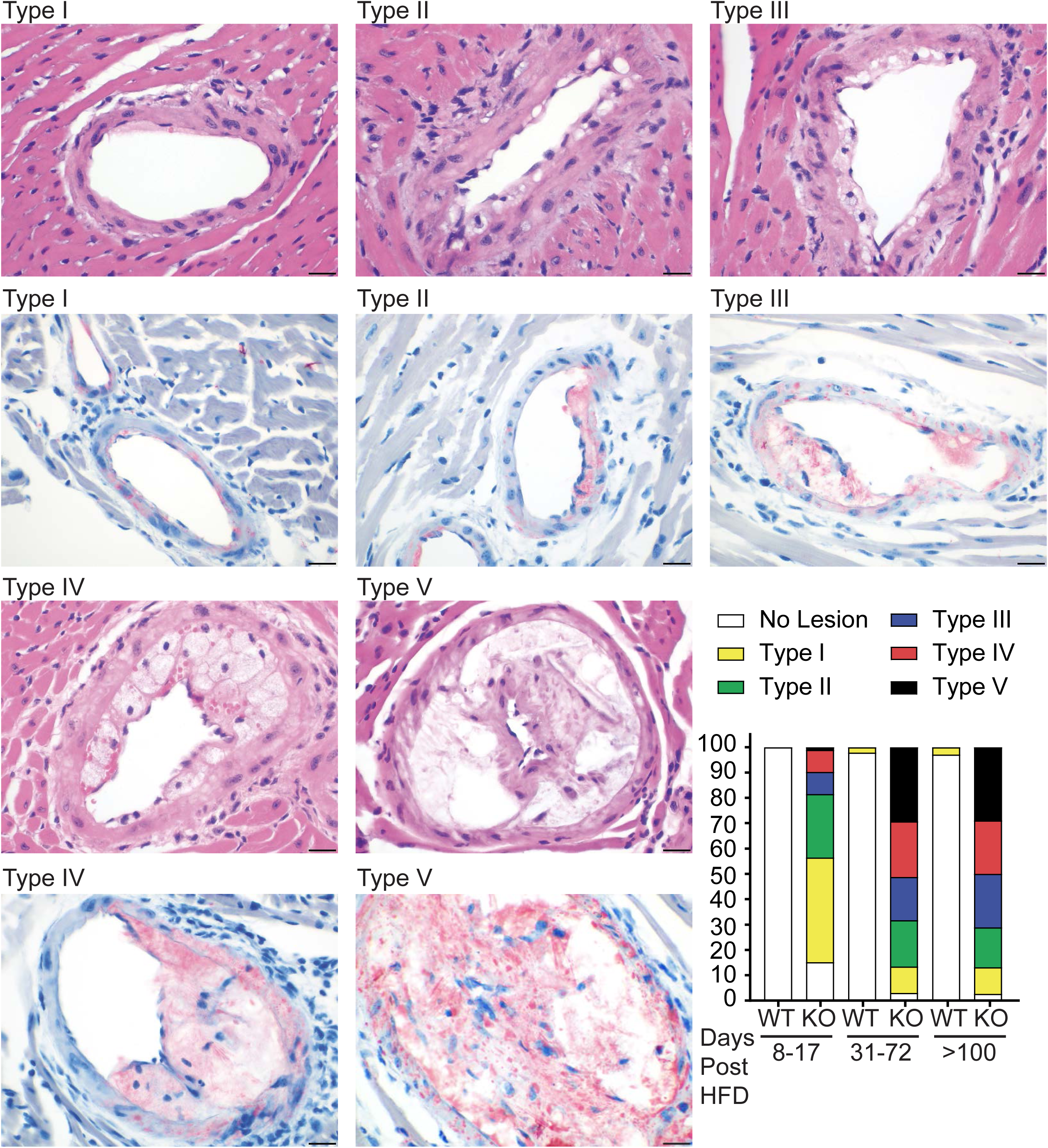
Stary classification of coronary lesions in *Lmod1^SMKO^* mice over time. H&E and ORO staining of representative lesion types in the *Lmod1^SMKO^* model, and the distribution of 504 lesions from 116 mice (combined sexes) at three different time intervals post-regimen. Scale bars, 20 μm. See also **Table S6**.

**Figure S15.**
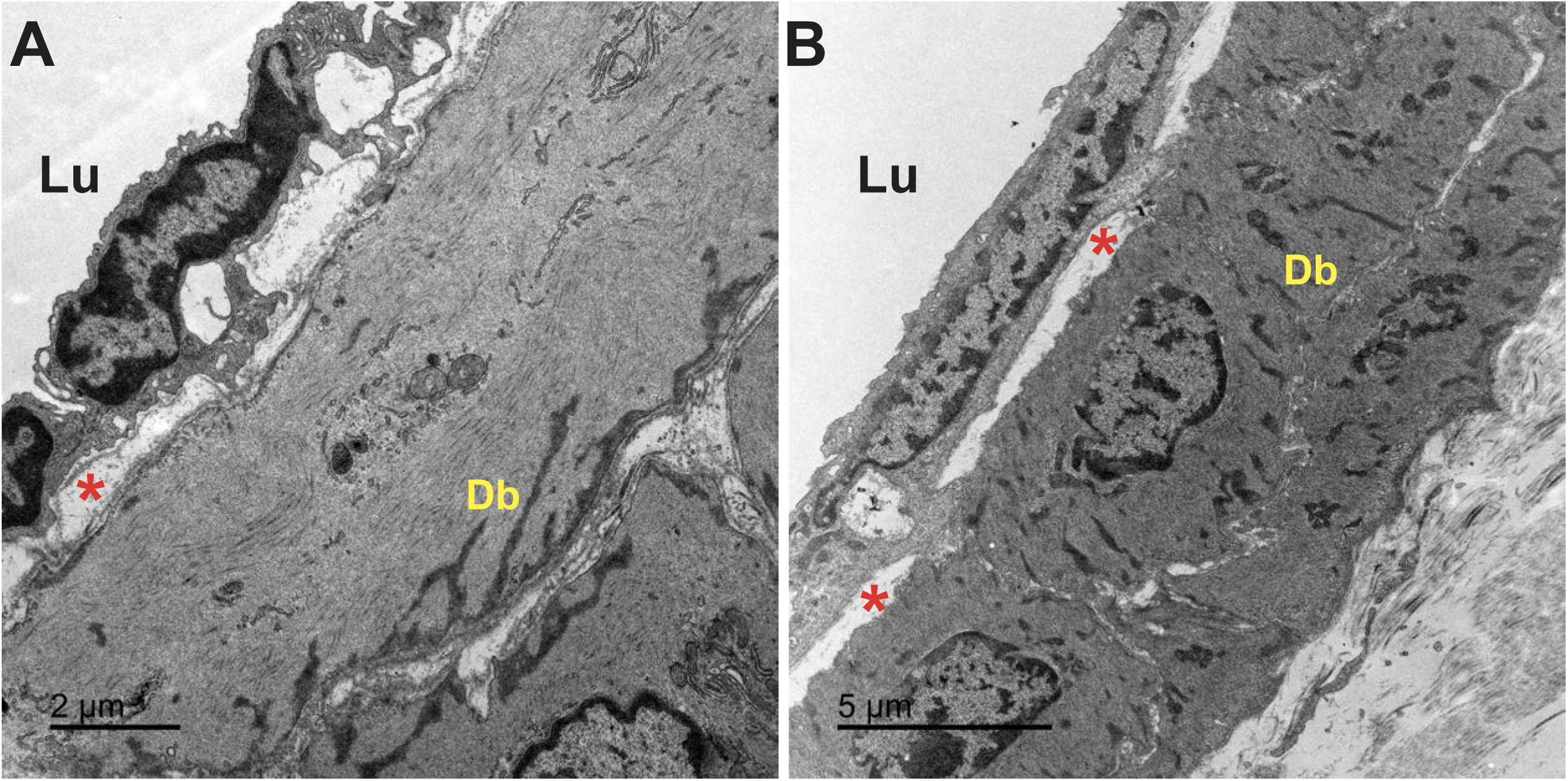
Electron microscopy of control *Lmod1^WT^* coronary arteries. (**A**) Coronary artery from an *Lmod1^WT^* mouse subjected to PCSK9/HFD for 37 days. Large dense bodies (Db) were concentrated at the abluminal surface of SMCs as described. ^5^ (**B**) Coronary artery from a C57BL6/J mouse fed a normal chow diet. Scale bars are 2 μm and 5 μm in panels **A** and **B**, respectively. Red asterisk, internal elastic lamina. Lu, lumen.

**Figure S16.**
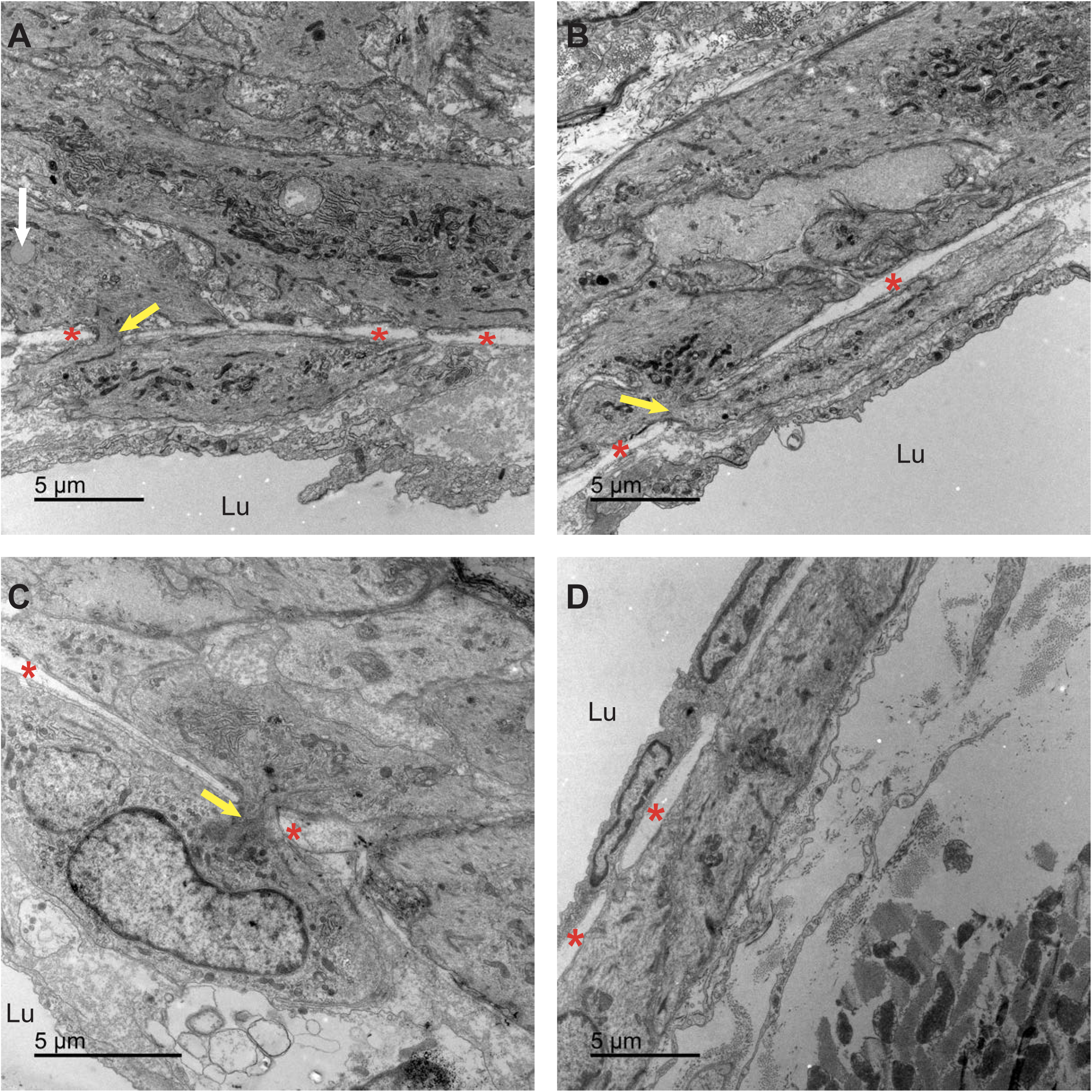
Migrating medial SMC in *Lmod1^SMKO^* mice. (A-C) Resident coronary artery SMC (defined by peripherally oriented dense bodies) undergoing migration through fenestra (yellow arrows) of the internal elastic lamina (red asterisks). Three different vessels from two *Lmod1^SMKO^* mice treated for eight days with PCSK9/HFD. Note the lipid droplet (white arrow at far left in panel A) in a migrating medial SMC. (D) A coronary artery from an *Lmod1^WT^* mouse treated eight days with PCSK9/HFD. Lu, lumen.

**Figure S17.**
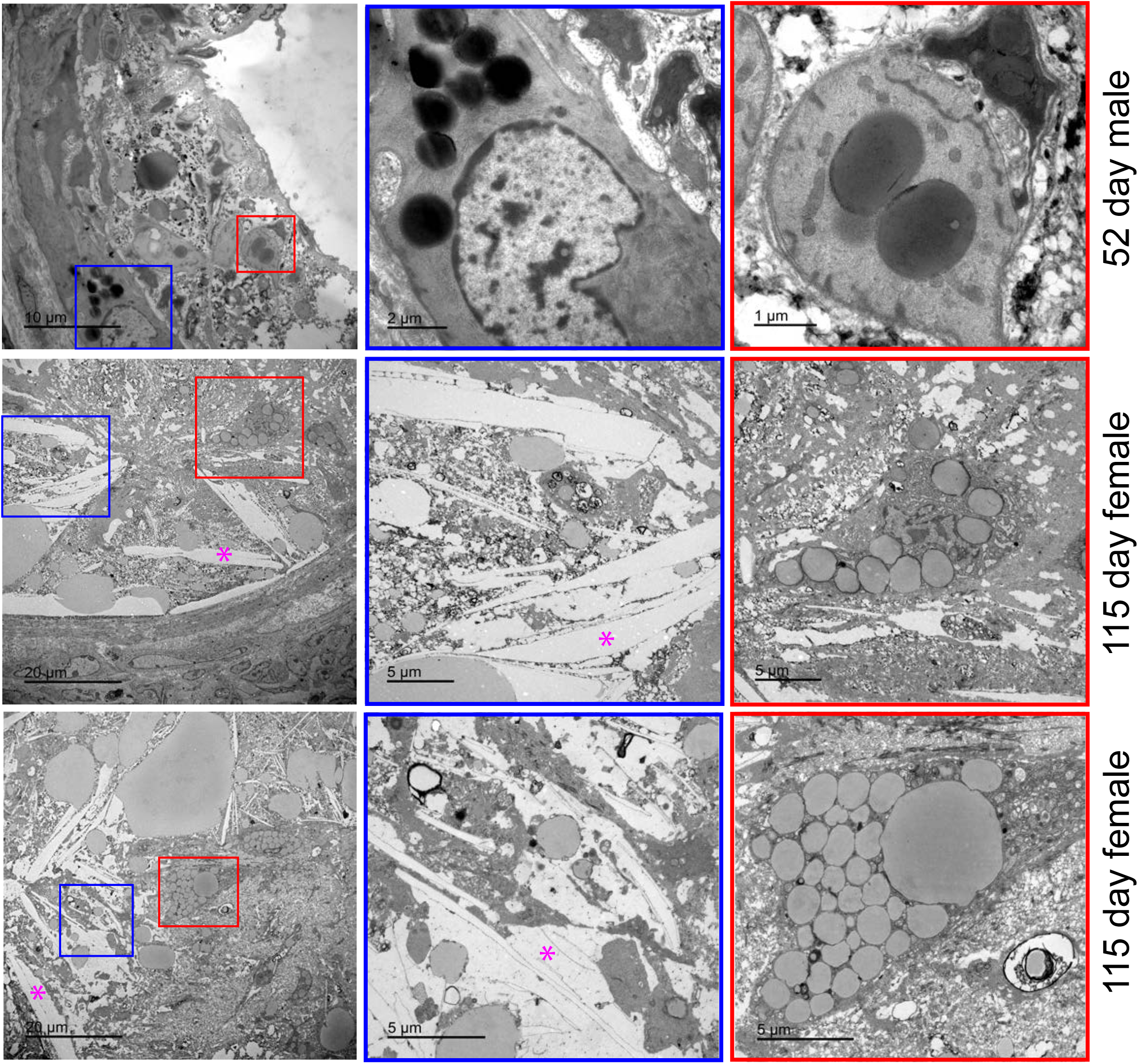
Lipid droplet heterogeneity and cholesterol crystals in coronary lesions of *Lmod1^SMKO^* mice. Electron micrographs of atheromas in a male mouse treated for 52 days (**top row**) and more advanced atheromas in coronary arteries from two females treated for 115 days (**middle, bottom rows**). Note the variable colors of lipid droplets, from white to gray and black, as well as the cholesterol crystals (magenta asterisks). The intimal foam cell at upper right panel is likely a SMC based on the appearance of dense bodies.

**Figure S18.**
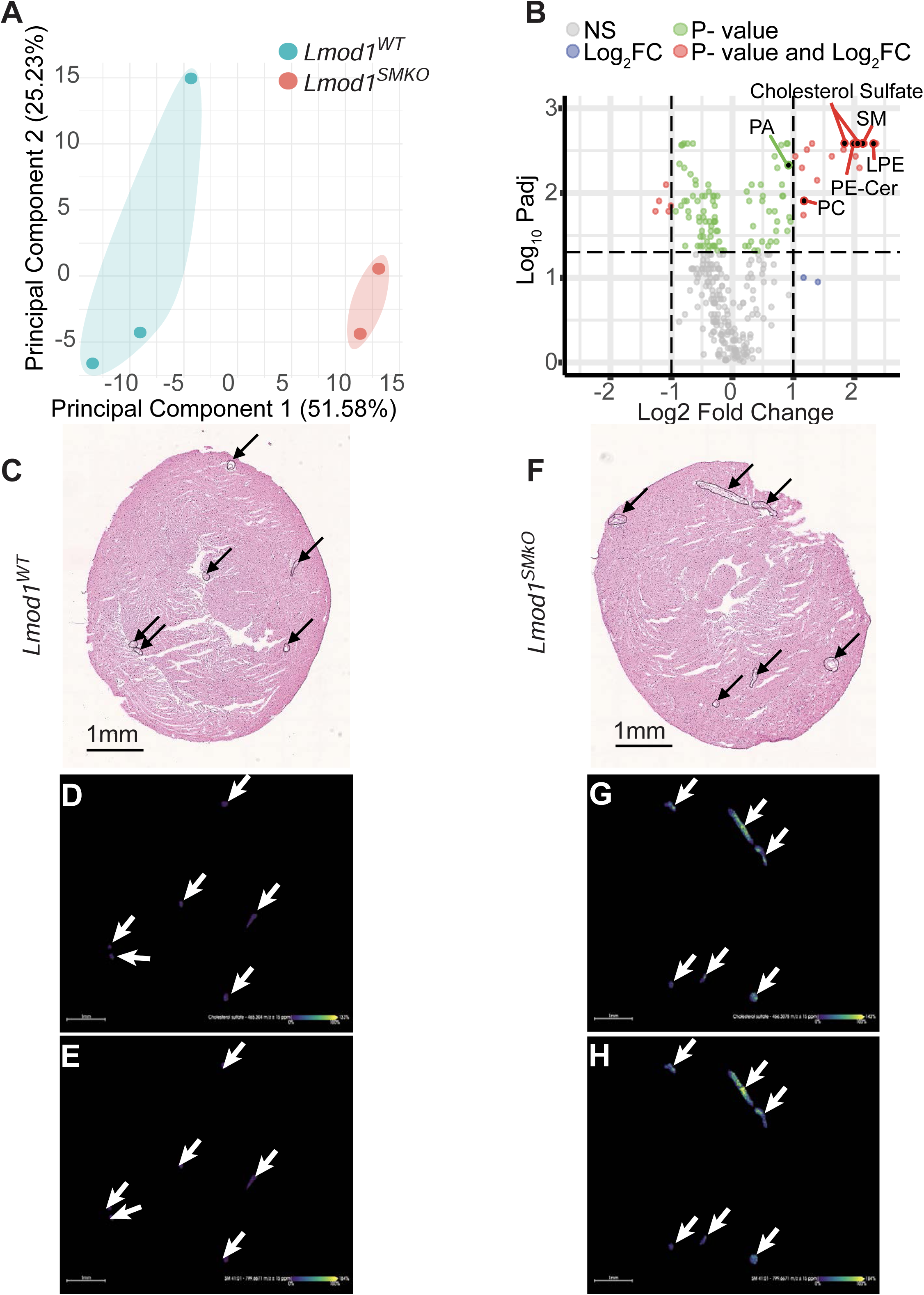
Spatial mass spectrometry imaging of coronary arteries from *Lmod1^WT^* and *Lmod1^SMKO^* mice. (**A**) Principal component analysis of coronary metabolomics demonstrates non-overlapping clustering between genotypes. (**B**) Volcano plot showing upregulated lipid molecules within coronary arteries of *Lmod1^SMKO^* mice. H&E stained *Lmod1^WT^* (**C**) and *Lmod1^SMKO^* (**F**) hearts with coronary arteries analyzed (arrows). Images of cholesterol sulfate (**D, G**) and membrane-associated sphingomyelin (**E,H**) in *Lmod1^WT^* (**D, E**) and *Lmod1^SMKO^* (**G, H**). The white arrows correspond to those in black in panels **C** and **F**. Images here are not in perfect scale with the H&E images due to compression and slight rotation. LPE, lyosphosphatidylethanolamine; PA, phosphatidic acid; PC, phosphatidylcholine; PE-Cer; phosphatidylethanolamine ceramide; SM, sphingomyelin species.

**Figure S19.**
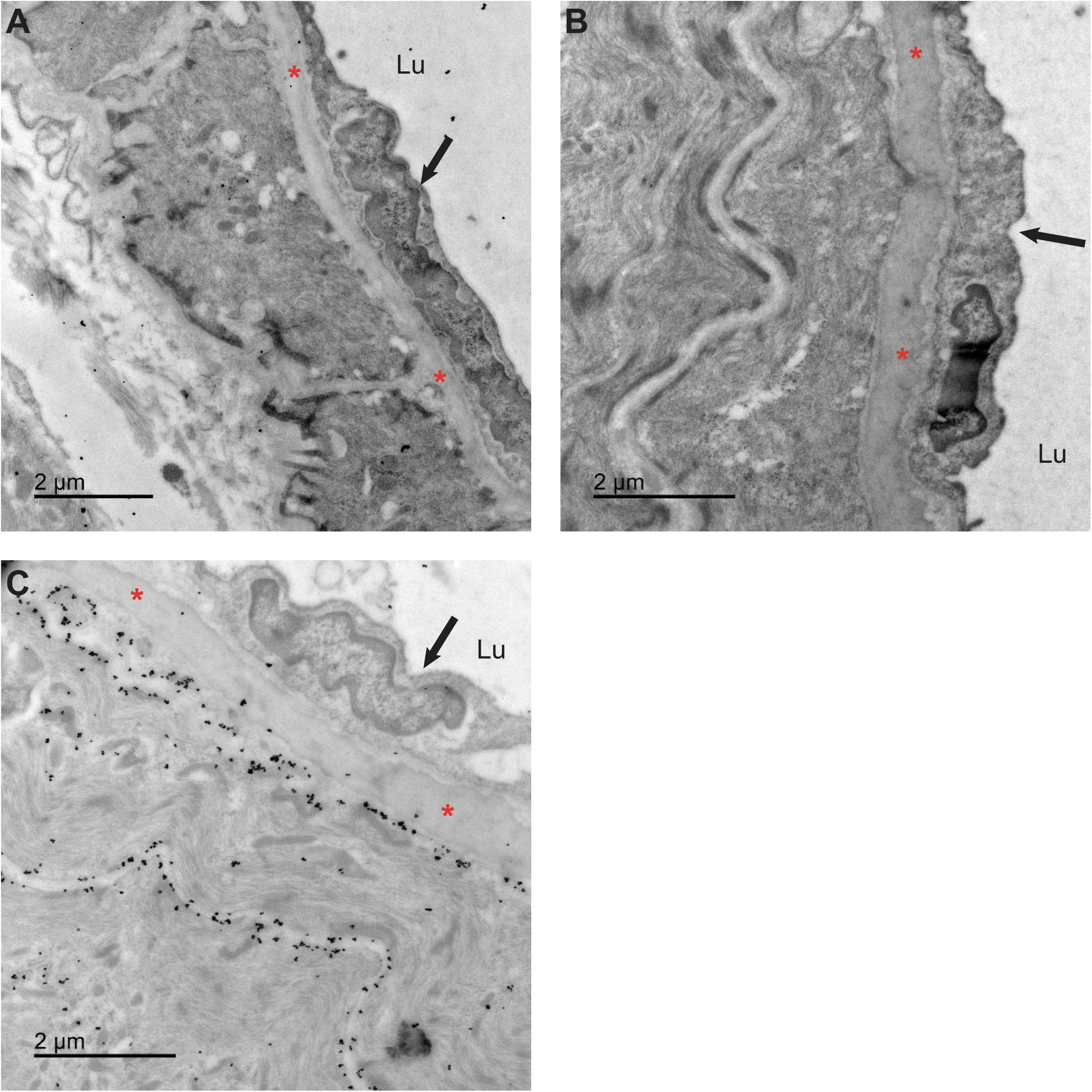
Control studies to optimize immunogold EM lineage tracing (IEMLT) of coronary arteries. Coronary arteries from *Lmod1^WT^* mice carrying the mTmG reporter and treated for six weeks with the PCSK9/HFD. Sections were processed for immunogold EM per methods using a primary antibody to GFP which was then detected with a secondary antibody conjugated to gold particles. (**A**) No gold particles were seen when GFP antibody was used on a coronary artery from a mouse that did not receive tamoxifen, as a test for leakiness or non-specific binding. (**B**) Similarly, no gold particles were seen when the GFP antibody was omitted in a mouse that received tamoxifen and therefore recombined out membrane tomato, thus allowing for membrane GFP expression. (**C**) Membrane associated gold particles observed in medial coronary SMC of a tamoxifen-treated mouse. Arrow and red asterisk in each panel indicate an endothelial cell and the internal elastic lamina, respectively. Lu, lumen.

**Figure S20.**
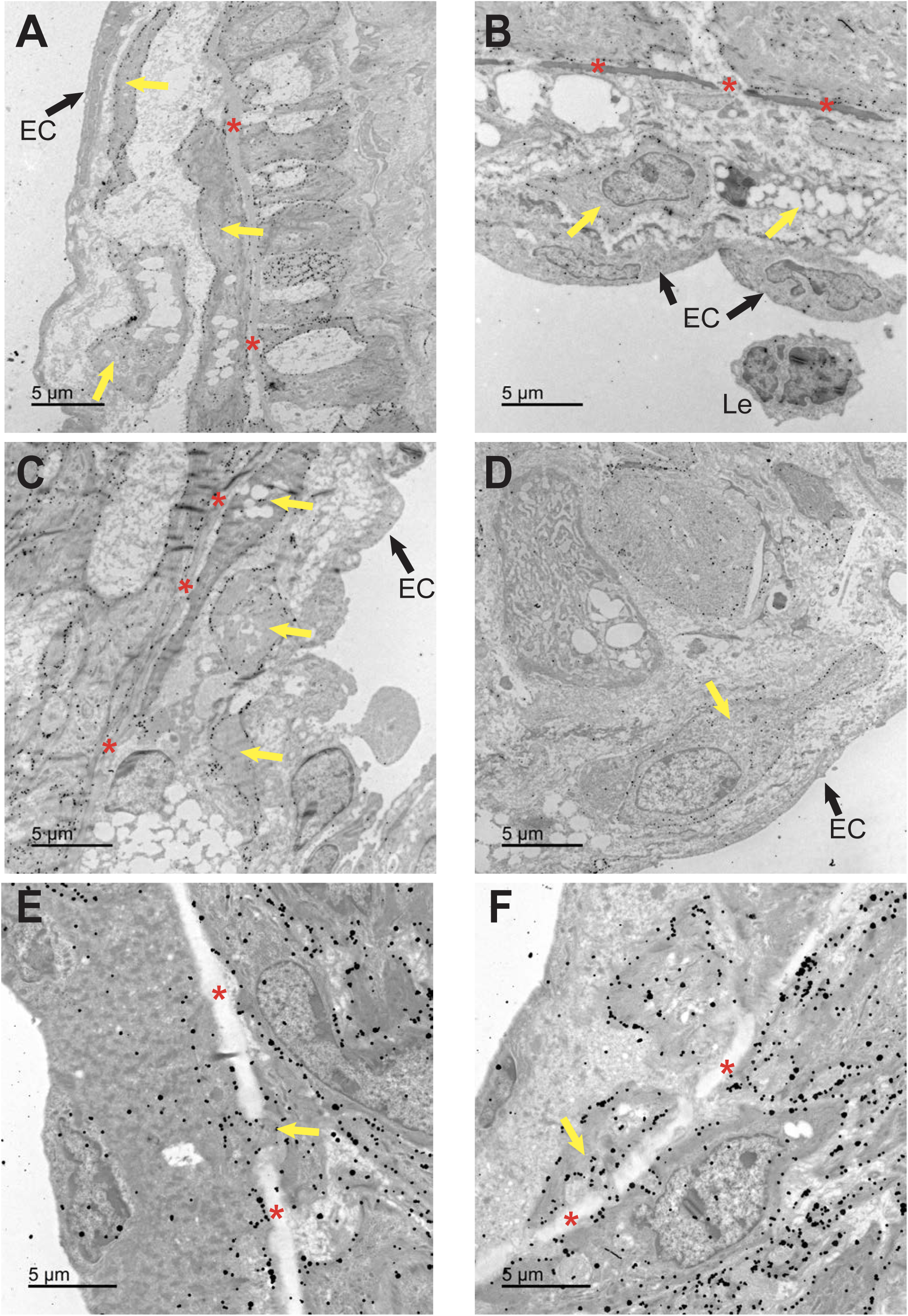
Additional IEMLT in *Lmod1^SMKO^* mice. *Lmod1^SMKO^* coronary arteries from four independent female mice subjected to PCSK9/HFD for six weeks showing (A) three GFP+ SMC-derived intimal cells (yellow arrows pointing to membrane-bound gold particles here and below) with one located just below a GFP-EC (black arrow) in an evolving cap and two containing lipid droplets in the core; (B) two intimal cells near the cap with one GFP+ (left yellow arrow) and the other GFP-(right yellow arrow), both located subjacent to two negative EC (black arrows); (C) three SMC-derived intimal cells (yellow arrows), with one (upper) containing lipid droplets; and (D) a non-lipid containing SMC-derived cap cell (yellow arrow) subjacent to a GFP-EC (black arrow). In a separate cohort of mice treated for only six days with PCSK9/HFD (E, F), GFP+ SMC could be seen moving through the fenestra (yellow arrows) of the internal elastic lamina (red asterisks here and elsewhere). The more intense gold particle staining in panels E and F relates to the timing for enhancement of gold particles in this cohort (see Methods). EC, endothelial cell; Le, leukocyte.

**Figure S21.**
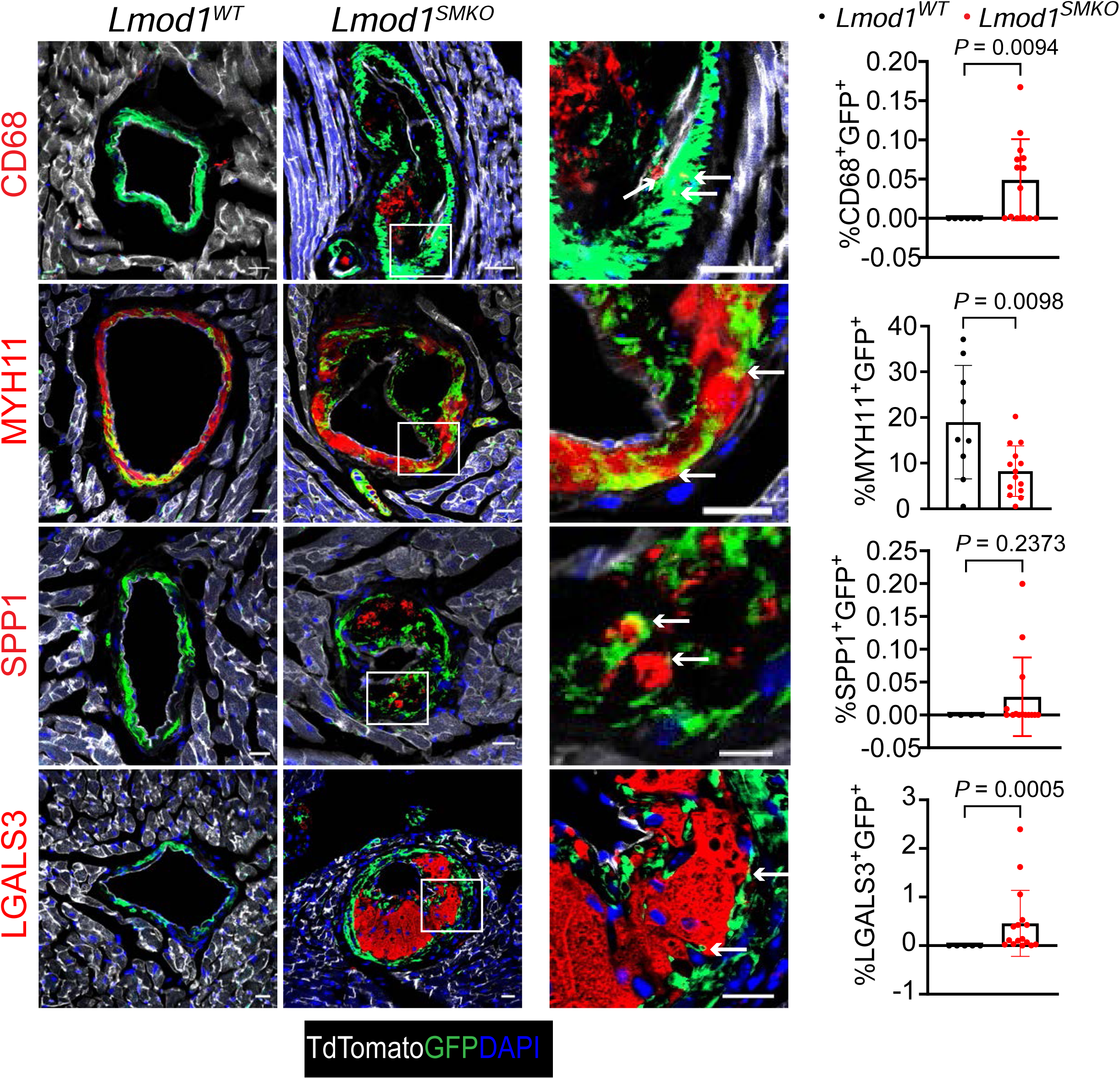
CIFM colocalization of GFP+ cells and markers of different cell states. Lineage-traced *c*oronary arteries from *Lmod1^WT^* and *Lmod1^SMKO^* mice after 63-days of PCSK9/HFD. GFP+ signals (green) colocalizing with each indicated marker (red) were imaged and quantitated (**far right**). The tdTomato signal predominating in surrounding cardiomyocytes was pseudo-colored white. Arrows point to colocalization of GFP and each marker. The number of replicates in graphs represents individual vessels from at least three mice of each genotype. Scale bars are 20 μm.

**Figure S22.**
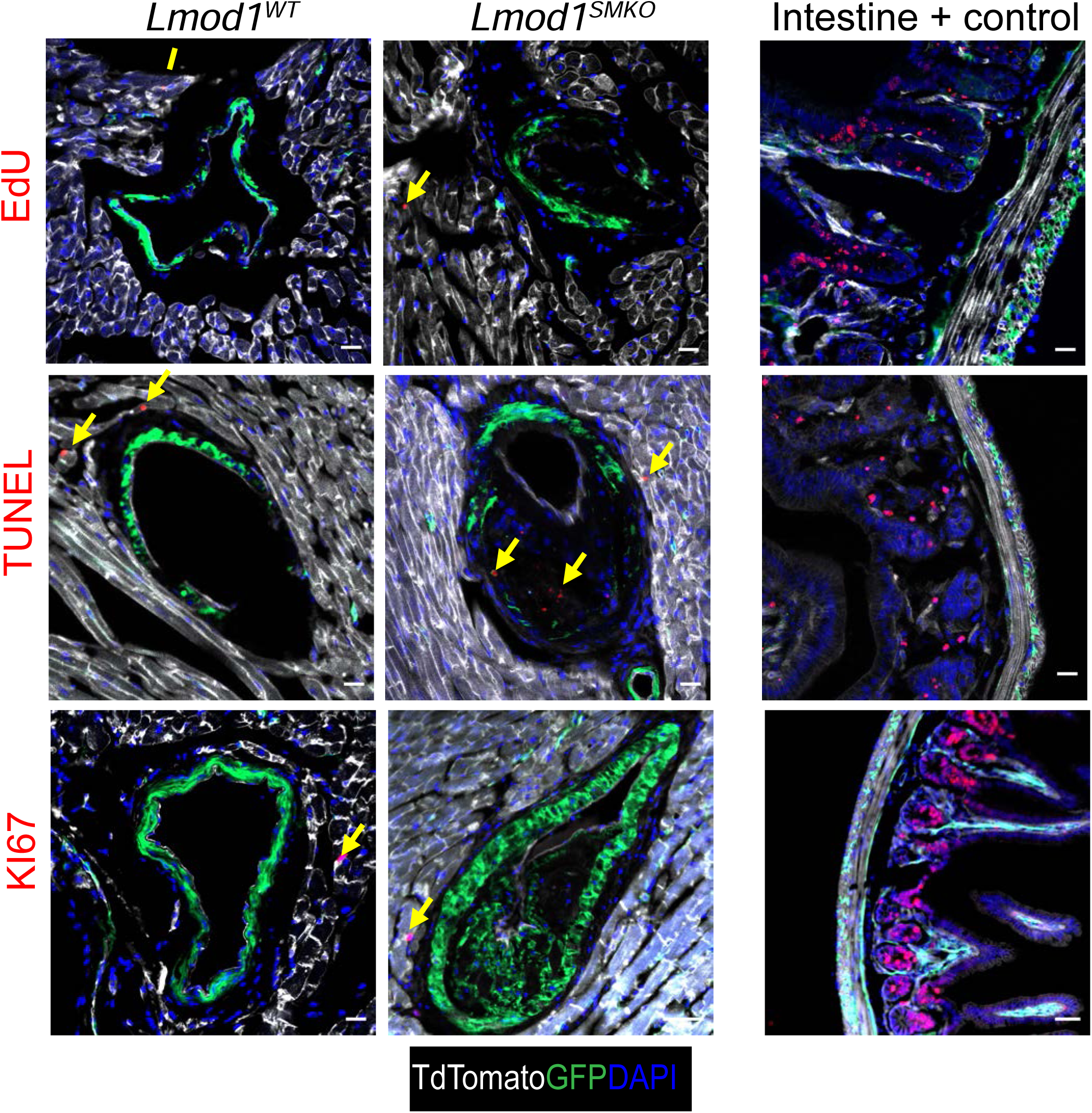
CIFM colocalization of GFP+ cells with growth and apoptotic markers. Lineage-traced coronary arteries from *Lmod1^WT^* and *Lmod1^SMKO^* mice after 63-days of PCSK9/HFD. The percentage of GFP+ signal (green) colocalizing with each indicated marker was imaged. Yellow arrows in each panel represent positive signals outside of GFP+ cells. Note TUNEL positive cells in the necrotic core of the *Lmod1^SMKO^* coronary artery. Each assay was validated with positive control tissue (intestine). Tomato signal was pseudo-colored white to allow easy identification of positive (red) signals. Scale bars are 20 μm.

**Figure S23.**
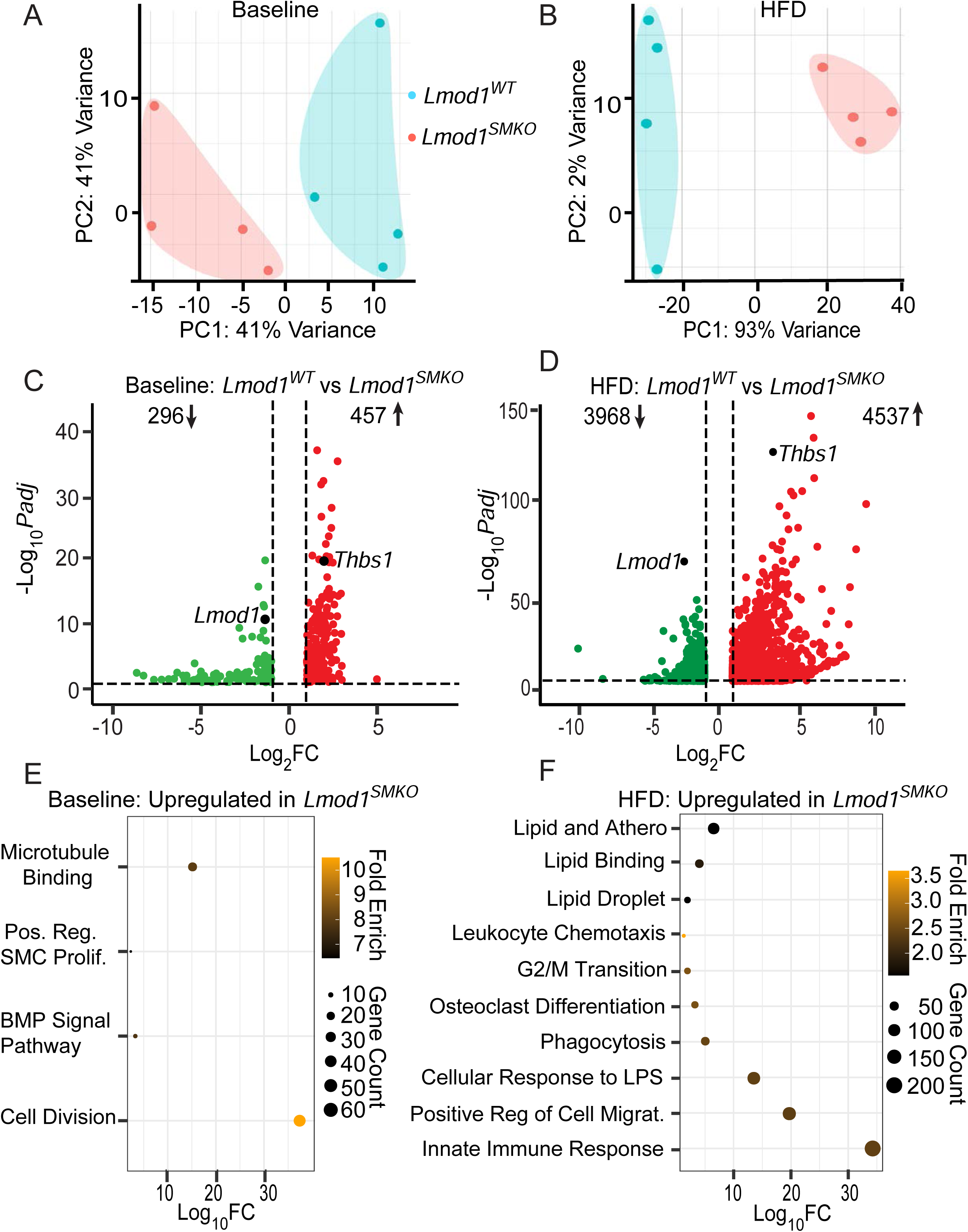
Baseline and PCSK9/HFD bulk RNA-seq of aorta. A, PCA plot displaying variance of wild type animals compared with *Lmod1^SMKO^* on chow diet and, B, high fat diet (HFD). Volcano plot showing differentially expressed genes (DEGs) under C, baseline and, D, HFD. Overrepresentation pathway analysis with upregulated pathways in *Lmod1^SMKO^* over *Lmod1^WT^* mice shown under E, chow diet and, F, HFD.

**Figure S24.**
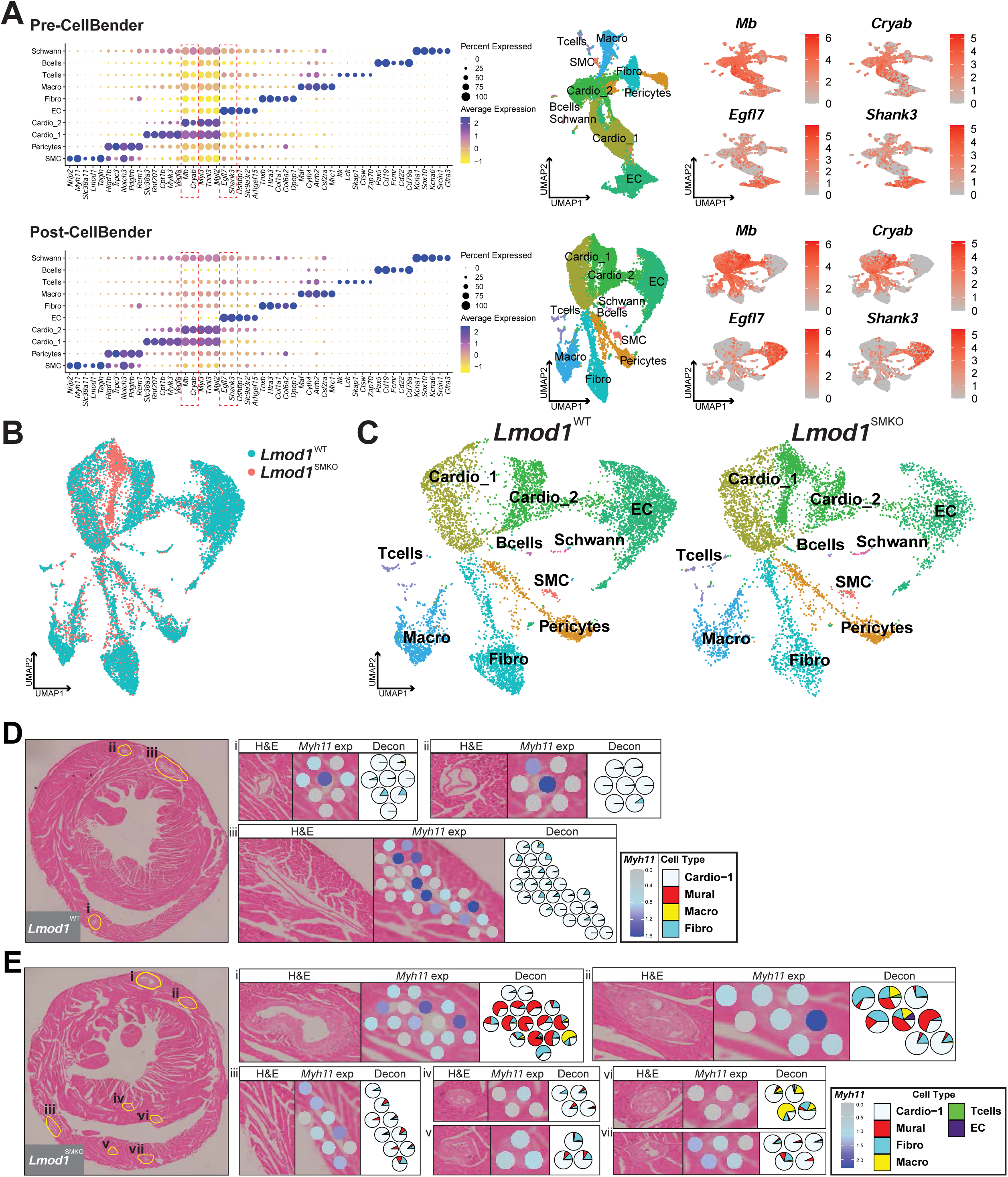
Multi-modal transcriptomic workflow for ambient RNA correction, dataset integration, and spatial deconvolution. **(A)** Pre-CellBender (**top**) dot plot of annotated cell-type markers with representative UMAPs (cardiomyocytes: *Mb*, *Cryab*; endothelial cells: *Egfl7*, *Shank3*) demonstrates ambient RNA artifacts in scRNA-seq data, indicated by red dotted boxes and diffuse low-level expression across clusters. Post-CellBender (**bottom**) ambient RNA correction restricts marker expression to appropriate cell types in both dot plots and UMAPs. **(B)** UMAP visualization of integrated scRNA-seq data with samples overlaid and **(C)** split by genotype, demonstrating correction for batch effects. **(D, E)** Genotype-matched spatial voxel deconvolution for *Lmod1^WT^* (three coronary arterial regions, CA) and *Lmod1^SMKO^* (seven CA regions) shown with corresponding H&E-stained cross sections, with CA regions highlighted in yellow. Each CA region has *Myh11* spatial feature plots paired with deconvolution voxels, done by genotype-matched scRNA-seq and spatial modalities.

**Figure S25.**
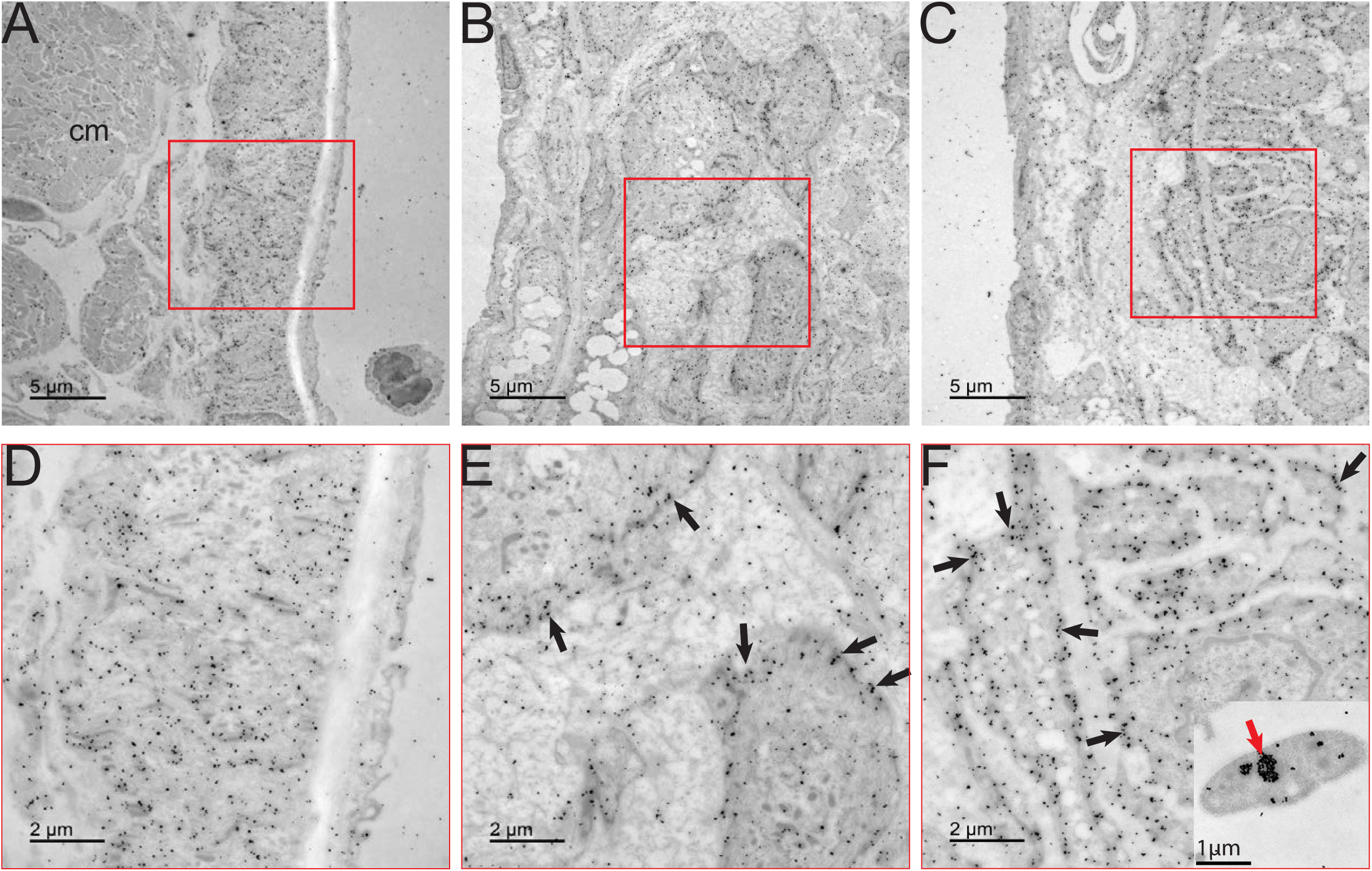
THBS1 immunogold EM in coronary arteries of *Lmod1^SMKO^* mice. Low (A-C) and high (D-F) magnification images of *Lmod1^WT^* (A, D) and *Lmod1^SMKO^* (B, E, C, F) coronary arteries stained for THBS1 (black dots). Note the diffuse pattern of gold particles in the *Lmod1^WT^* coronary (D) versus a more membrane-associated localization in *Lmod1^SMKO^* (arrows in E, F). Red arrow represents a positive internal control for THBS1 in alpha granule of an adjacent platelet, a rich source of THBS1. ^6^

**Figure S26.**
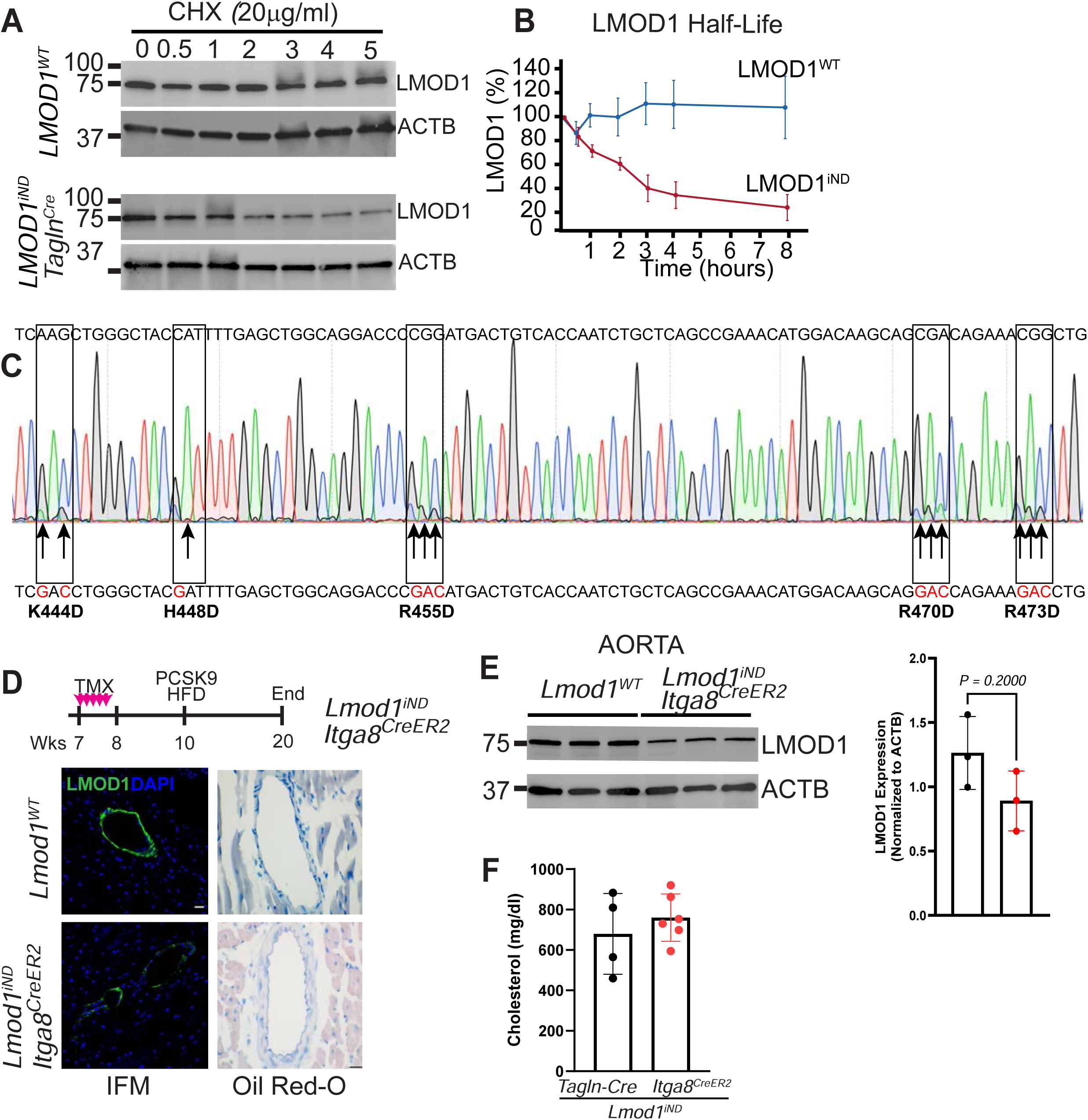
Effect of *Lmod1^iND^* on CAD phenotype in *Lmod1^SMKO^* mice. (A) Western blot of LMOD1 protein t1/2 in *Lmod1^WT^* (top) and *Lmod1^iND^* (bottom) MASMCs following various times of Cycloheximide (CHX) exposure. (B) Quantitation of LMOD1 bands in panel A (n=3 independent experiments). (C) Electropherogram demonstrating sequence fidelity of the five DNA sequence-encoding amino acid substitutions (indicated in red at bottom with arrows). (D) IFM of LMOD1 and ORO staining of coronary arteries from each mouse model after 10 weeks of PCSK9/HFD. (E) Western blotting and quantitation of aortic LMOD1 from each genotype. (F) Total cholesterol measures in *Lmod1^iND^* mice crossed with *Tagln-Cre* (black dots) or *Itga8-CreER^T2^* (red dots) to induce the *Lmod1^iND^* mutant.

