## Supplemental Table 9 for "Loss of the Coronary Artery Disease Risk Gene *Leiomodin1* in Vascular Smooth Muscle Cells Triggers Rapid Onset Coronary Atherosclerosis"

| **Genetic Model** | **Diet-Intervention** | **Coronary Lesion** | **PMID** |
| --- | --- | --- | --- |
| *Apoe* KO | 40w WD | Occlusive lesion; fibrosis; no MI | 8274468 |
| *Rora* KO | 9w 15% butter/1.25% cholesterol/0.5% cholate | Fatty streaks; Female>Male | 9851961 |
| *Apoe*/*Ldlr* dKO | 7m WD (21% fat/0.2% cholesterol) | Advanced plaque, MI, ECG abnormal | 10359814 |
| *Apoe*/*Nos3* dKO | >16w WD (21% fat/0.2% cholesterol) | Occlusive lesion; myocardial fibrosis | 11468208 |
| *Apoe*/*Scarb1* dKO | 8w chow diet | Advanced plaque; MI; death 5-8w | 11861414 |
| *Apoe^-/-^*/*Plau^Mac-Tg^* | 10-15w HFD (21% fat/0.2% cholesterol) | Occlusive; MI; death at 11w | 15096455 |
| *Apoe* KO | 60w chow diet | Intramyocardial foam cells | 15914296 |
| *Apoe^-/-^*/*Olr1^Tg^* | 3w 20% casein/15% butter/1.25% cholest/0.5% cholate | Intramyocardial foam cells | 15961718 |
| *Scarb1^-/-^/Apoe^T61R^* | 4w 7.5% butter/15.8% fat/1.25% cholest/0.5% cholate | Occlusive; death by MI in 7w | 15967843 |
| *Apoe*/*Akt1* dKO | 14w HFD (20% fat/1.25% cholesterol)* | Foam cells; some death by 14w | 18054314 |
| *Ldlr*/*Apob48* dKO | 2-3m WD (21% fat/0.2% cholesterol) | Intramyocardial foam cells | 18243216 |
| *Nos1/Nos2/Nos3* tKO | 3-11m chow diet | Occlusive; MI from 3-11m | 18413498 |
| *Apoe*/*Pdzk1* dKO | 12w 7.5% butter/15.8% fat/1.25% cholest/0.5% cholate | Advanced plaque; MI within 4w | 19956623 |
| *Ldlr^-/-^*/*Lrp6^R611C^* | 4m 40% fat/1.25% cholesterol/0.5% cholate | Mild lesions | 26489464 |
| *Cd38* KO | 12w WD (21% fat/0.2% cholesterol) | Mild lesions | 26818887 |
| *Apoe^-/-^*/*Fbn1^C1039G^* | 35w WD (21% fat/0.2% cholesterol) | Occlusive; Fibrosis and MI; death 20w | 24553721 |
| *Apoe/Pdgfrb^SMC-D849V^* | 16w WD (21% fat/0.2% cholesterol) | Advanced lesions and death 12-19w | 26183159 |
| *Apoe* KO | 4w chow diet plus TAC | Fibrofatty lesions within 4w of TAC | 26800562 |
| *Ldlr*/*Scarb1* dKO | 10w 20% lard/0.5% cholesterol | Advanced plaque/MI; all death at 20w | 27373983 |
| *Scarb1^ΔCT^/Apoe^-/-^* | 9w chow diet | Advanced plaque/MI; median death at 9w | 27694217 |
| *Ldlr^C699Y^*/*Hprt^EC-ALPL^* | 16w 21% fat/1.25% cholesterol/0.5% cholate | Advanced lesions at 16w | 29023576 |
| *Apoe^-/-^*/*Lmna^G609G^* | 14w chow diet | Occasional 3/9 lesions and death | 29490993 |
| *Apoe^-/-^*/*Lmna^SMC-LCS^* | 43w chow diet | Frequent 8/9 lesions/calcification and death | 29490993 |
| *Apoe^-/-^*/*Ins^C96Y^* | 25w WD (21% fat/0.2% cholesterol) | Intramyocardial lesions; death at 25w | 29879686 |
| *Ldlr*/*Gpihbp1* dKO | 3m STZ+LFD (5% lard/0.05% cholesterol) | Mild ORO lesions/death with HFD | 30721842 |
| *Scarb1^Liver^* KO | 6-8w PCSK9/HFD (20% fat/1.25% cholesterol)* | Fibrofatty lesions | 31019307 |
| *Apoe*/*Timp1* dKO | 36w HFD (21% lard/0.15% cholesterol) | Advanced lesions; MI and death | 34848763 |
| *Apoe/Apoa1* dKO | 30w chow diet | ORO-positive coronary lesions | 35587694 |
| *Scarb1 ^ΔCT^/Ldlr^-/-^* | 26w WD (21% fat/0.2% cholesterol) | MI/stroke; median death by 20w | 38881440 |
| *Ccn2^SMC-^* ^KO^ | 24w PCSK9 + WD | Coronary lesions; insignificant deaths | 39167826 |
| *Scarb1* KO | 12w HFD (15% fat/1.25% chol/0.5% cholate) | Occlusive and death in 40w fed mice | 40403091 |
| *Apoe^SA/SA^* | 4w WD+Doxycycline to induce AngII | CAD in 4 weeks with death from MI | 40485474 |

**Table S9. Mouse models of CAD**
